# Assessing the impact of the threatened crucian carp (*Carassius carassius*) on pond invertebrate diversity - a comparison of conventional and molecular tools

**DOI:** 10.1101/2020.03.30.015677

**Authors:** Lynsey R. Harper, Lori Lawson Handley, Carl D. Sayer, Daniel S. Read, Marco Benucci, Rosetta C. Blackman, Matthew J. Hill, Bernd Hänfling

## Abstract

Fishes stocked for recreation and angling can damage freshwater habitats and negatively impact biodiversity. The pond-associated crucian carp (*Carassius carassius*) is rare across Europe and stocked for conservation management in England, but impacts on pond biota are understudied. Freshwater invertebrates contribute substantially to aquatic biodiversity, encompassing many rare and endemic species, but their small size and high abundance complicates their assessment. Practitioners have employed sweep-netting and kick-sampling with microscopy (morphotaxonomy), but specimen size/quality and experience can bias identification. DNA and eDNA metabarcoding offer alternative means of invertebrate assessment. We compared invertebrate diversity in ponds (*N* = 18) with and without crucian carp using morphotaxonomic identification, DNA metabarcoding, and eDNA metabarcoding. Five 2-L water samples and 3-minute sweep-net samples were collected at each pond. Inventories produced by morphotaxonomic identification of netted samples, DNA metabarcoding of bulk tissue samples, and eDNA metabarcoding of water samples were compared. Alpha diversity was greatest with DNA or eDNA metabarcoding, depending on whether standard or unbiased methods were considered. DNA metabarcoding reflected morphotaxonomic identification, whereas eDNA metabarcoding produced markedly different communities. These complementary tools should be combined for comprehensive invertebrate assessment. Crucian carp presence minimally reduced alpha diversity in ponds, but positively influenced beta diversity through taxon turnover (i.e. ponds with crucian carp contained different invertebrates to fishless ponds). Crucian carp presence contributes to landscape-scale invertebrate diversity, supporting continued conservation management in England. Our results show that molecular tools can enhance freshwater invertebrate assessment and facilitate development of more accurate and ecologically effective pond management strategies.

## 1. Introduction

Freshwater ecosystems comprise <1% of the Earth’s surface, but represent major biodiversity hotspots and provide vital ecosystem services (Dudgeon et al., 2006). Ponds (see definition in Biggs, von Fumetti, & Kelly-Quinn, 2016) in particular provide critical habitat for biodiversity in a fragmented landscape (Céréghino, Biggs, Oertli, & Declerck, 2008), supporting many rare and specialist species not found in other waterbodies (Biggs et al., 2016; Wood, Greenwood, & Agnew, 2003). These highly diverse and species-rich ecosystems contribute considerably to regional-scale diversity due to high habitat heterogeneity and associated species turnover (Davies et al., 2008; Williams et al., 2003). Aquatic invertebrates are a crucial and abundant component of this diversity, and occupy the vast range of microhabitats made available in ponds by their broad-ranging physicochemical properties and habitat complexity (Davies et al., 2008; Williams et al., 2003; Wood et al., 2003).

In Europe, lakes and ponds are commonly stocked with fish for angling and recreation purposes, despite often having negative effects on invertebrates and amphibians (Gledhill, James, & Davies, 2008; Wood, Greenwood, Barker, & Gunn, 2001). Fish can alter community structure (Schilling, Loftin, & Huryn, 2009a, 2009b; Wood et al., 2001), reduce diversity (Lemmens et al., 2013; Wood et al., 2001), and reduce the abundance and biomass (Marklund, Sandsten, Hansson, & Blindow, 2002; Schilling et al., 2009a) of invertebrates. These effects may manifest through direct predation by fish, altered water quality and loss of macrophyte diversity and cover via foraging activity of fish, or management practices (e.g. commercial farming, artificial feeding, pond depth and bank alteration, removal of aquatic vegetation and sediment) associated with angling activity (Lemmens et al., 2013; Maceda-Veiga, López, & Green, 2017; Schilling et al., 2009a; Wood et al., 2001). However, the impact of fish stocking can be negligible or even beneficial to invertebrate diversity, particularly at a regional-scale, provided that fish species are carefully selected and managed (Gee, Smith, Lee, & Griffiths, 1997; Hassall, Hollinshead, & Hull, 2011; Lemmens et al., 2013; Stefanoudis et al., 2017).

The crucian carp (*Carassius carassius*) is one of few fish species strongly associated with small ponds in Europe, but has suffered heavy declines and local extinctions in the last century (Copp & Sayer, 2010; Sayer et al., 2011) due to habitat loss, species displacement by the invasive goldfish (*Carassius auratus*) and gibel carp (*Carassius gibelio*) (Copp, Warrington, & Wesley, 2008; Sayer et al., 2011), and genetic introgression through hybridisation with goldfish, gibel carp, and common carp (*Cyprinus carpio*) (Hänfling, Bolton, Harley, & Carvalho, 2005). In 2010, the crucian carp was designated as a Biodiversity Action Plan (BAP) species in the county of Norfolk in eastern England (Copp & Sayer, 2010). A key objective of this BAP is to increase the number of viable crucian carp populations across Norfolk through pond restoration and species reintroduction. Many Norfolk ponds have since been restored and stocked with crucian carp to realise this objective (Sayer et al., 2020), but maintaining populations and continued stocking requires justification in light of genetic evidence that indicates the crucian carp is not native to the UK (Jeffries et al., 2017). Nonetheless, there is support for UK conservation efforts to continue to protect the genetic integrity of the crucian carp at the European level, and provide a natural stronghold for the species (Harper et al., 2019; Jeffries et al., 2017; Stefanoudis et al., 2017) in the face of persistent declines throughout its native range of Northwest and Central Europe (Copp et al., 2008; Sayer et al., 2011; Sayer et al., 2020).

The impact of stocking crucian carp on lentic biodiversity has been largely ignored, and little is known about its interactions with other pond species. Existing research suggests crucian carp are characteristic of ponds rich in invertebrates with extensive macrophyte cover (Copp et al., 2008; Sayer et al., 2011), and play an important ecological role (along with other pond-associated fishes) by increasing landscape-scale diversity across pond networks (Stefanoudis et al., 2017). Yet to our knowledge, only one study has assessed biodiversity (specifically macrophytes, zooplankton, and water beetles) in ponds with and without pond-associated fishes in the UK (Stefanoudis et al., 2017), and no studies have specifically focused on crucian carp. Consequently, there is a need to survey and compare fishless ponds to ponds containing crucian carp to assess the impact of maintaining existing crucian carp populations and conservation-based stocking of this species on invertebrate diversity more broadly.

Assessments of invertebrate diversity in pond networks have relied upon morphotaxonomic identification using sweep-netting together with microscopy, but this approach requires extensive resources and taxonomic expertise for accurate species-level identification (Briers & Biggs, 2003; Haase, Pauls, Schindehütte, & Sundermann, 2010; Hill et al., 2018). Metabarcoding offers a rapid, high-resolution, cost-effective approach to biodiversity assessment, where entire communities can be identified using High-Throughput Sequencing (HTS) in conjunction with community DNA from bulk tissue samples (DNA metabarcoding), or environmental DNA (eDNA) from environmental samples (eDNA metabarcoding), such as soil or water (Deiner et al., 2017; Taberlet, Coissac, Pompanon, Brochmann, & Willerslev, 2012). DNA metabarcoding of aquatic invertebrate samples has proven relatively successful, with applications in biomonitoring (Andújar et al., 2017; Elbrecht, Vamos, Meissner, Aroviita, & Leese, 2017a; Emilson et al., 2017; Lobo, Shokralla, Costa, Hajibabaei, & Costa, 2017). Use of eDNA metabarcoding for invertebrate assessment in freshwater rivers (Blackman et al., 2017; Carew, Kellar, Pettigrove, & Hoffmann, 2018; Deiner, Fronhofer, Mächler, Walser, & Altermatt, 2016; Klymus, Marshall, & Stepien, 2017; Leese et al., 2020), streams (Macher et al., 2018), and lakes (Klymus et al., 2017) is also gaining traction, but there are currently few published studies that have used metabarcoding for small lake or pond invertebrates (Beentjes, Speksnijder, Schilthuizen, Hoogeveen, & van der Hoorn, 2019).

In this study, we assessed invertebrate diversity in UK ponds with and without crucian carp using metabarcoding in conjunction with morphotaxonomic identification. The entire taxon inventories generated by each monitoring tool (standard methods) were compared to evaluate which tool provided the most holistic assessment of invertebrate diversity. We also compared the taxon inventories produced by each method when potential biases were removed, i.e. taxa with no reference sequences and meiofauna that would not be captured by a 1 mm mesh net (unbiased methods). The effect of crucian carp presence on invertebrate diversity was then determined using the taxon inventories from the standard or unbiased methods individually and combined. We hypothesised that: 1) DNA metabarcoding and morphotaxonomic identification would produce congruent invertebrate communities (e.g. Carew et al., 2018; Emilson et al., 2017; Nichols et al. 2020), whereas eDNA metabarcoding would reveal taxa not identified by these tools (e.g. Macher et al., 2018; Mächler et al., 2019); 2) DNA and eDNA metabarcoding would enable species resolution data for problematic taxa that cannot be morphotaxonomically identified to species-level using standard keys (e.g. Blackman et al., 2017; Elbrecht et al., 2017a; Lobo et al., 2017; Nichols et al., 2020); and 3) alpha diversity would be lower in ponds with crucian carp due to direct predation or altered habitat quality (e.g. Lemmens et al., 2013; Maceda-Veiga, López, & Green, 2017; Schilling et al., 2009a; Wood et al., 2001), but beta diversity would be enhanced due to community heterogeneity induced by crucian carp presence in the pond network (e.g. Stefanoudis et al., 2017). The third pattern was expected regardless of the monitoring tool used. We provide recommendations for the application of molecular tools to freshwater invertebrate assessment as well as impact assessment and conservation management of the non-native crucian carp alongside pond biodiversity.

## 2. Methods

### 2.1 Study sites

We surveyed nine ponds containing crucian carp across Norfolk (*n* = 6) and East Yorkshire (*n* = 3), and nine fishless ponds in Norfolk (Fig. S1). Crucian carp were either present in ponds as well-established populations, or individuals that were restocked between 2010 and 2017 as part of the species’ BAP (Copp & Sayer, 2010) because ponds contained populations between 1970-1980 according to local anglers (Sayer et al., 2011; Sayer et al., 2020). The fish status of ponds was confirmed by prior fyke-net surveys (Sayer et al., 2011; Sayer et al., 2020). For ponds containing crucian carp, the last fyke-net survey undertaken was 2016 (*n* = 7), 2015 (*n* = 1), and 2013 (*n* = 1). Roach (*Rutilus rutilus*) and rudd (*Scardinius erythrophthalmus*) were the only other fishes that occurred in three and one of the Norfolk ponds respectively, but at much lower density (see Table S1). All study ponds were selected to be similar in terms of morphometry and habitat structure, being <1 ha in area, <5 m in depth, dominated by submerged and floating-leaved macrophytes, with a largely open-canopy and thus minimal shading (≤20%) of the water surface. Ponds were mainly surrounded by arable fields, excluding one pond located in woodland. As invertebrate diversity often peaks in autumn (Hill, Sayer, & Wood, 2016) but large and more easily identifiable odonate larvae are typically present in summer prior to emergence (Brooks & Cham, 2014), invertebrate specimens were collected for morphotaxonomic identification and DNA metabarcoding alongside water samples for eDNA metabarcoding in late August 2016. Data on physical (area, depth, estimated percentages of perimeter with emergent vegetation, emergent macrophyte cover, submerged macrophyte cover, and shading) and chemical (conductivity measured with a HACH HQ30d meter) properties of the ponds were collected between May and August from 2010 to 2017.

### 2.2 Sweep-netting and morphotaxonomic identification

Sweep-netting was performed in accordance with the UK National Pond Survey methodology (Biggs, Fox, & Nicolet, 1998), using a 1 mm mesh long-handled net (0.3 m square bag), to generate a conventional taxonomic inventory of lentic macroinvertebrates. Sampling time at each pond comprised a 3 min sweep-net sample and 1 min hand search (see Supporting Information). Samples were deposited in a 1.2 L sterile Whirl-Pak® stand-up bag (Cole-Palmer, London) and stored at -20 °C until laboratory processing and sorting. Samples were thawed and passed through sieves of 8 mm, 2 mm, and 0.25 mm in succession to remove large items of vegetation and detrital matter. For each sample, invertebrate specimens were sorted into three body size categories on laminated millimetre graph paper: small (<2.5 × 5 mm), medium (2.5 × 5 mm up to 5 × 10 mm), and large (>10 mm and up to 10 × 20 mm) (Elbrecht, Peinert, & Leese, 2017b). Where possible, size-sorted specimens were identified under a light microscope to species-level using Freshwater Biological Association publications (Bass, 1998; Brooks & Cham, 2014; Edington & Hildrew, 1995; Elliott, 2009; Elliott & Humpesch, 2010; Elliott & Dobson, 2015; Friday, 1988; Gledhill, Sutcliffe, & Williams, 1993; Macan, 1960; Savage, 1989; Wallace, Wallace, & Philipson, 1990). Specimens that could not be identified to species-level were identified to genus or family-level, except Hydrachnidea, Tabanini, Tanytarsini, Oligochaeata, and Collembola which were recorded as such. Terrestrial taxa and empty cases/shells were discarded. The laminated paper was sterilised with 50% v/v chlorine-based commercial bleach solution (Elliot Hygiene Ltd, UK) and 80% v/v ethanol solution between measuring specimens from different ponds to minimise cross-contamination risk. Specimens were preserved in sterile 15 mL falcon tubes (SARSTED, Germany) containing 100% ethanol according to size category and pond sampled, and stored at -20 °C until DNA extraction.

### 2.3 DNA samples

We followed the DNA extraction procedure from Blackman *et al*. (2017). Size categories from each pond were dried overnight on FisherBrand cellulose filter paper (Fisher Scientific UK Ltd, UK) in sterile glass funnels and conical flasks to remove excess ethanol. Size categories were lysed (3 × 30 sec) using a TissueLyser II (Qiagen®, Germany) with DigiSol (50 mM Tris, 20 M EDTA, 120 mM NaCl and 1% SDS). The adapter sets held 1.5 g of dried tissue and corresponding volume of DigiSol. If the dry tissue weight of any size category exceeded 1.5 g, we processed the size category in batches until all tissue had been lysed. The lysates from all batches were pooled to recreate size categories. The size categories were incubated overnight at 55 °C with SDS and Proteinase K (Bioline®, UK), following which 200 μL of lysate from each size category was used for extraction with the DNeasy Blood & Tissue Kit (Qiagen®, Germany) according to the manufacturer’s protocol. Specimens collected from two ponds were categorised as medium or large only, whereas specimens collected from other study ponds were categorised as small, medium, and large (Table S2). Consequently, 16 bulk tissue samples were each represented by three DNA extracts and two bulk tissue samples each represented by two DNA extracts that were stored at -20 °C until metabarcoding.

### 2.4 eDNA samples

We repurposed eDNA samples from Norfolk ponds that were collected by Harper et al. (2019) to validate a quantitative PCR (qPCR) assay for crucian carp. Additional samples were taken from East Yorkshire ponds for the present study. Details of water sample collection, filtration, and DNA extraction are provided in the Supporting Information. Briefly, five 2 L surface water samples were collected at equidistant points around the pond perimeter where access permitted. For each pond, a full process blank (1 L molecular grade water) was taken into the field, transported, filtered, and extracted alongside samples to identify contamination. Filters were stored at -20 °C until DNA extraction with the PowerWater® DNA Isolation Kit (MO BIO Laboratories, CA, USA) following the manufacturer’s protocol. eDNA extracts were stored at -20 °C until metabarcoding.

### 2.5 Metabarcoding workflow

Our metabarcoding workflow is fully described in the Supporting Information. In short, a comprehensive list of UK invertebrate species living in or associated with freshwater habitats (expected fauna) was used to create custom, curated reference databases for UK aquatic invertebrates, excluding Diptera (Appendix 1). Public GenBank records for Diptera were missing record features (e.g. ‘gene’ or ‘CDS’) and/or names were not in the format required for custom reference database construction using the selected bioinformatic tools. Assignments to Diptera were therefore made using the entire NCBI nucleotide database (see below). For the remaining taxonomic groups, 1483 species were represented in our custom reference database (78.97% of the expected fauna), although representation varied strongly across groups (Fig. S2). Species without database representation are listed in Table S3 (see also Figs. S2, S3). Published metazoan primers mICOIintF (Leray et al., 2013) and jgHCO2198 (Geller, Meyer, Parker, & Hawk, 2013), which amplify a 313 bp fragment of the mitochondrial cytochrome c oxidase subunit I (*COI*) gene, were validated *in silico* using ecoPCR software (Ficetola et al., 2010) against the reference databases. Parameters set allowed a 250-350 bp fragment and maximum of three mismatches between each primer and each reference sequence (Appendix 2). Primers were validated *in vitro* against 38 invertebrate species, representing 38 families and 10 major groups, and compared to published macroinvertebrate metabarcoding primers (Elbrecht & Leese, 2017; Vamos, Elbrecht, & Leese, 2017) (Figs. S4, S5, S6).

PCR conditions were optimised (Figs. S7, S8) before two independent libraries were constructed for DNA and eDNA metabarcoding using a two-step PCR protocol (Appendix 1). During the first PCR (performed in triplicate), the target region was amplified using oligonucleotides comprising the locus primers, sequencing primers, and pre-adapters. DNA from the exotic, terrestrial two-spotted assassin bug (*Platymeris biguttatus*) was used for PCR positive controls (tissue DNA *n* = 9, eDNA *n* = 11) as this species is not found in the UK, whilst sterile molecular grade water (Fisher Scientific UK Ltd, UK) substituted template DNA for PCR negative controls (tissue DNA *n* = 9, eDNA *n* = 11). PCR products were individually purified using a magnetic bead clean-up (VWR International Ltd, UK), following a double size selection protocol from Bronner et al. (2009). The second PCR bound Multiplex Identification (MID) tags to the purified products. PCR products were pooled according to PCR run and the pooled PCR product purified using a magnetic bead clean-up, following a double size selection protocol from Bronner et al. (2009). Each pooled purified PCR product was quantified on a Qubit(tm) 3.0 fluorometer using a Qubit(tm) dsDNA HS Assay Kit (Invitrogen, UK) and normalised according to concentration and sample number to produce a pooled volume of 20 μL.

The pooled libraries were quantified by qPCR using the NEBNext® Library Quant Kit for Illumina® (New England Biolabs® Inc., MA, USA). An Agilent 2200 TapeStation and High Sensitivity D1000 ScreenTape (Agilent Technologies, CA, USA) were used to verify removal of secondary products from libraries by bead purification and that a fragment of the expected size (531 bp) remained. A total of 52 bulk tissue subsamples sequenced in triplicate (*n* = 156), 12 extraction blanks, and 18 PCR controls were included alongside samples from other projects (*N* = 188) in the DNA library. A total of 90 eDNA samples, 18 full process blanks, and 22 PCR controls were included alongside samples from other projects (*N* = 140) in the eDNA library. The DNA and eDNA libraries were sequenced on an Illumina® MiSeq using 2 × 300 bp V3 chemistry with 10% and 20% PhiX Sequencing Control respectively (Illumina, Inc, CA, USA).

Raw sequences were converted to taxonomic assignments using metaBEAT v0.97.11, a custom pipeline for reproducible analysis of metabarcoding data (https://github.com/HullUni-bioinformatics/metaBEAT). After quality filtering, trimming, merging, chimera detection, and clustering, non-redundant query sequences were compared against our reference databases using BLAST (Zhang, Schwartz, Wagner, & Miller, 2000). Putative taxonomic identity was assigned using a lowest common ancestor (LCA) approach based on the top 10% BLAST matches for any query matching with at least 90% identity to a reference sequence across more than 80% of its length. Unassigned sequences were subjected to a separate BLAST against the complete NCBI nucleotide (nt) database at 90% identity to determine the source via LCA as described above. Bioinformatic settings (Appendix 1) were chosen based on comprehensive exploration of the parameter space and comparison of metaBEAT taxonomic assignments to morphotaxonomic inventories. Reference databases in GenBank or fasta format and the bioinformatic analysis have been deposited in a dedicated GitHub repository, which has been permanently archived for reproducibility (https://doi.org/10.5281/zenodo.3993125).

### 2.6 Data analysis

All analyses were performed in R v.3.6.3 (R Core Team, 2017), with data and R scripts deposited in the GitHub repository. Manipulation of the DNA and eDNA metabarcoding datasets for analysis is described in the Supporting Information. For each method, two presence-absence datasets (species-level and family-level) were generated using the *decostand* function in vegan v2.5-6 (Oksanen et al., 2018) for downstream analyses. This is because potential amplification bias during PCR can prevent reliable abundance or biomass estimation from sequence reads produced by DNA or eDNA metabarcoding (Elbrecht et al., 2017b). The entire species and family inventories produced by each method (hereafter standard methods) were compared. We also compared inventories from each method after correcting for known biases: taxa without reference sequences (i.e. those which cannot be detected with metabarcoding) were removed from the morphotaxonomic dataset, and meiofauna (i.e. taxa that would escape a 1 mm mesh net) were removed from the metabarcoding datasets (hereafter unbiased methods). The effect of crucian carp on invertebrate diversity was then assessed at species and family-level according to standard methods (main text) and unbiased methods (Supporting Information) individually and combined. Finally, the impact of crucian carp alongside environmental variables on invertebrate diversity was investigated using the combined standard method (main text) and unbiased method (Supporting Information) datasets. For species-level analyses, only species records were used, whereas family-level analyses included species, genus and family records grouped at the taxonomic rank of family.

We define alpha diversity as taxon richness (species or families) within individual ponds, and beta diversity as the difference between communities present at each pond whilst accounting for taxon identity (Baselga & Orme, 2012). Jaccard dissimilarity was used as a measure of beta diversity for all analyses. For each standard or unbiased dataset, the following analyses were performed. Alpha diversity (response variable) was obtained using the *specnumber* function in vegan v2.5-6 and modelled against sampling method (explanatory variable), then modelled against crucian carp presence-absence (explanatory variable). Using the combined method datasets, alpha diversity of the major invertebrate groups, i.e. orders, classes, or phyla (Dobson, Pawley, Fletcher, & Powell, 2012), in each pond (response variable), was modelled against crucian carp presence-absence (explanatory variable). All alpha diversity comparisons were made using negative binomial Generalized Linear Models (GLMs), and Pairwise Tukey’s HSD tests were used to assess significance of differences. We used betapart v1.5.1 (Baselga & Orme, 2012) to estimate total beta diversity, partitioned into nestedness-resultant and turnover components, across sampling methods and ponds using the *beta*.*multi* function. These three beta diversity components were estimated for inventories produced by each sampling method, and ponds with or without crucian carp using the *beta*.*pair* function (pairwise beta diversity scores). For each beta diversity component, we compared community heterogeneity in each group (sampling method or crucian carp presence-absence) by calculating homogeneity of multivariate dispersions (MVDISP) using the *betadisper* function from vegan v2.5-6. Variation in MVDISP was statistically tested using an analysis of variance (ANOVA), and pairwise Tukey’s HSD tests used to determine if there were significant differences between the groups. The effect of sampling method and crucian carp on each beta diversity component was visualised using Non-metric Multidimensional Scaling (NMDS) ordination plots with the *metaMDS* function, and tested statistically using permutational multivariate analysis of variance (PERMANOVA) with the *adonis* function in vegan v2.5-6. Pre-defined cut-off values were used for effect size, where PERMANOVA results were interpreted as moderate and strong effects if R^2^ > 0.09 and R^2^ > 0.25 respectively. These values are broadly equivalent to correlation coefficients of *r* = 0.3 and 0.5 which represent moderate and strong effects accordingly (Macher et al., 2018; Nakagawa & Cuthill, 2007).

We tested whether the species and family-level invertebrate communities produced by the combined standard or unbiased methods were influenced by pond properties in conjunction with crucian carp presence-absence. Redundancy Analysis (RDA) was selected for constrained ordination as it analyses variation in biological communities in response to explanatory variables (Legendre & Legendre, 2012). Principal Coordinate Analysis (PCoA) was performed using the *pcoa* function in ape v5.3 (Paradis & Schliep, 2018) on the turnover, nestedness-resultant, and total beta diversity matrices. The Lingoes correction was employed to account for negative eigenvalues (Legendre, 2014). The resultant PCoA eigenvectors (principle coordinates) for each distance matrix were used as the response variable in variance partitioning. Our variables were grouped as biotic (crucian carp presence-absence) or abiotic (pond conductivity, area, depth, percentages of perimeter with emergent vegetation, emergent macrophyte cover, submerged macrophyte cover, and shading). Abiotic variables were log10 transformed to eliminate their physical units (Legendre & Birks, 2012). Significant abiotic variables influencing each component of beta diversity were identified using the *ordiR2step* function in vegan v2.5-6 to perform separate RDA analyses under a forward selection procedure. Where applicable, the relative contributions of the biotic and abiotic variables on turnover, nestedness-resultant, and total beta diversity for the species and family-level invertebrate communities were assessed by variance partitioning (Borcard, Legendre, & Drapeau, 1992) using the *varpart* function in vegan v2.5-6. For each beta diversity component, RDA was performed using biotic and significant abiotic variables, and variance partitioning used to divide the total percentage of variation explained into unique and shared contributions for biotic and abiotic predictor groups. The *anova* function in vegan v2.5-6 was used to examine the statistical significance of the full model and the unique contributions of each predictor group. We report the adjusted R^2^-fractions as they are widely recommended and unbiased (Peres-Neto, Legendre, Dray, & Borcard, 2006).

## 3. Results

### 3.1 Comparison of methods for freshwater invertebrate assessment

Summaries of taxon detection by standard and unbiased methods are reported in the Supporting Information (Appendix 2; Tables S4, S5, S6, S7; Figs. S9, S10). Briefly, using microscopy, we identified 2,281 specimens belonging to 38 families, of which 1,404 specimens belonged to 91 species. With DNA metabarcoding, 2,906,869 sequence reads were assigned to 55 families, of which 2,448,078 sequence reads were assigned to 131 species. With eDNA metabarcoding, 813,376 sequence reads were assigned to 90 families, of which 346,163 sequence reads were assigned to 145 species. Only 19 species (Figs. 1ai, bi) and 22 families (Figs. 1aii, bii) were shared by all three methods (standard or unbiased). Standard methods detected more distinct taxa than unbiased methods, especially morphotaxonomic identification and eDNA metabarcoding. eDNA metabarcoding detected the most unique species and families, whereas DNA metabarcoding and morphotaxonomic identification were more comparable (Fig. 1). The number of taxa that morphotaxonomic identification shared with DNA and eDNA metabarcoding was unchanged using standard or unbiased methods. Detection of one family was exclusive to morphotaxonomic identification and eDNA metabarcoding, whereas detection of 36 species and 13 families was exclusive to morphotaxonomic identification and DNA metabarcoding (Fig. 1). Morphotaxonomic identification detected 19 species without reference sequences, whereas DNA and eDNA metabarcoding detected 4 and 41 meiofaunal species respectively (Fig. 1a). The metabarcoding approaches either shared 44 species and 16 families (standard methods, Fig. 1a) or 33 species and 8 families (unbiased methods, Fig. 1b).

**Figure 1.**
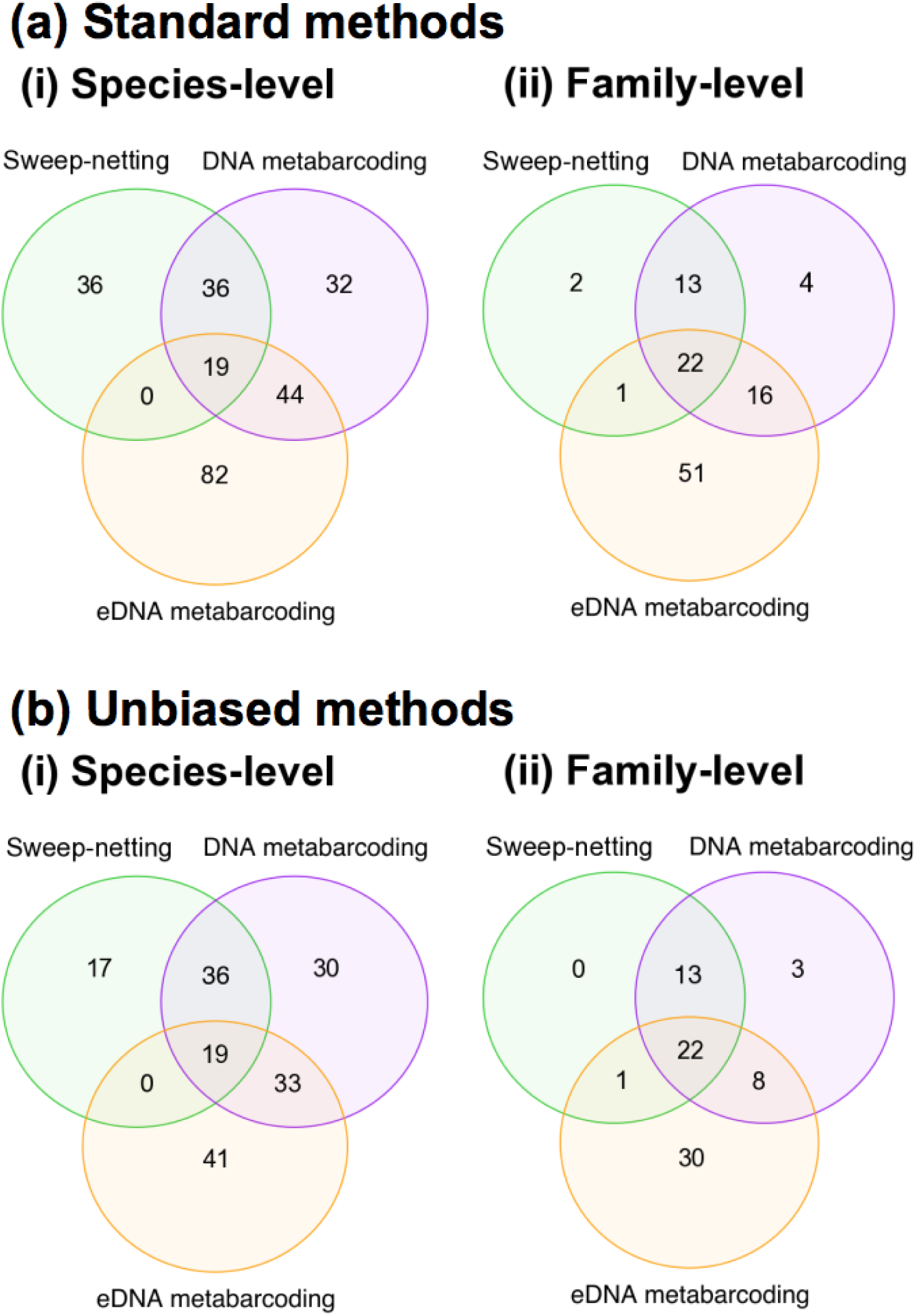
Venn diagrams summarising taxon detection by standard **(a)** and unbiased **(b)** methods of invertebrate assessment. The number of invertebrate species **(i)** and families **(ii)** detected across the 18 study ponds by sweep-netting and microscopy (green circle), DNA metabarcoding (purple circle), and eDNA metabarcoding (orange circle) is displayed. Overlap in species or family detections between methods is shown within circle intersections.

Standard sampling method influenced alpha diversity of ponds at species (GLM χ^2^_2_ = 26.058, *P* < 0.001) and family-level (GLM χ^2^_2_ = 48.238, *P* < 0.001). Significant differences (*P* < 0.05) between the alpha diversity means for morphotaxonomic identification and DNA metabarcoding, morphotaxonomic identification and eDNA metabarcoding, and DNA and eDNA metabarcoding (family-level only) were observed at species and family-level (Table 1).

**Table 1.** Summary of analyses (Pairwise Tukey’s HSD Test) statistically comparing alpha diversity at species-level and family-level between standard and unbiased methods.

|  | Species-level |  |  | Family-level |  |  |
| --- | --- | --- | --- | --- | --- | --- |
|  | Pairwise Tukey's HSD Test |  |  | Pairwise Tukey's HSD Test |  |  |
|  | Estimate ± SE | Z | P | Estimate ± SE | Z | P |
| **Standard methods** |  |  |  |  |  |  |
| Sweep-netting and microscopy :DNA metabarcoding | -0.399 ± 0.129 | -3.087 | **0.006** | -0.284 ± 0.108 | -2.630 | **0.023** |
| Sweep-netting and microscopy : eDNA metabarcoding | -0.645 ± 0.127 | -5.087 | **<0.001** | -0.686 ± 0.102 | -6.712 | **<0.001** |
| DNA metabarcoding : eDNA metabarcoding | 0.246 ± 0.121 | 2.030 | 0.105 | 0.402 ± 0.096 | 4.183 | **<0.001** |
| **Unbiased methods** |  |  |  |  |  |  |
| Sweep-netting and microscopy : DNA metabarcoding | -0.463 ± 0.128 | -3.632 | **<0.001** | -0.134 ± 0.106 | -1.267 | 0.414 |
| Sweep-netting and microscopy : eDNA metabarcoding | 0.128 ± 0.132 | -0.973 | 0.594 | 0.072 ± 0.110 | 0.650 | 0.792 |
| DNA metabarcoding : eDNA metabarcoding | -0.335 ± 0.125 | -2.671 | **0.021** | -0.206 ± 0.108 | -1.915 | 0.134 |

Conversely, unbiased sampling method influenced alpha diversity of ponds at species-level (GLM χ^2^_2_ = 14.614, *P* < 0.001), but not family-level (GLM χ^2^_2_ = 5.214, *P* = 0.074). Significant differences between the species-level alpha diversity means of morphotaxonomic identification and DNA metabarcoding, and DNA and eDNA metabarcoding were observed (Table 1). With standard methods, species and family-level alpha diversity was highest using eDNA metabarcoding, followed by DNA metabarcoding then morphotaxonomic identification (Figs. 2ai, aii). With unbiased methods, DNA metabarcoding detected more species than morphotaxonomic identification and eDNA metabarcoding, but alpha diversity was comparable at family-level (Figs. 2bi, bii). For beta diversity, MVDISP did not significantly differ between standard or unbiased methods for turnover, nestedness-resultant or total beta diversity at either taxonomic rank (Table S8). Both standard and unbiased sampling method had moderate and strong positive effects on turnover and total beta diversity at species (Figs. 3ai, aiii, ci, ciii) and family-level (Figs. 3aii, aiv, cii, civ) respectively, but not nestedness-resultant (Table S9).

**Figure 2.**
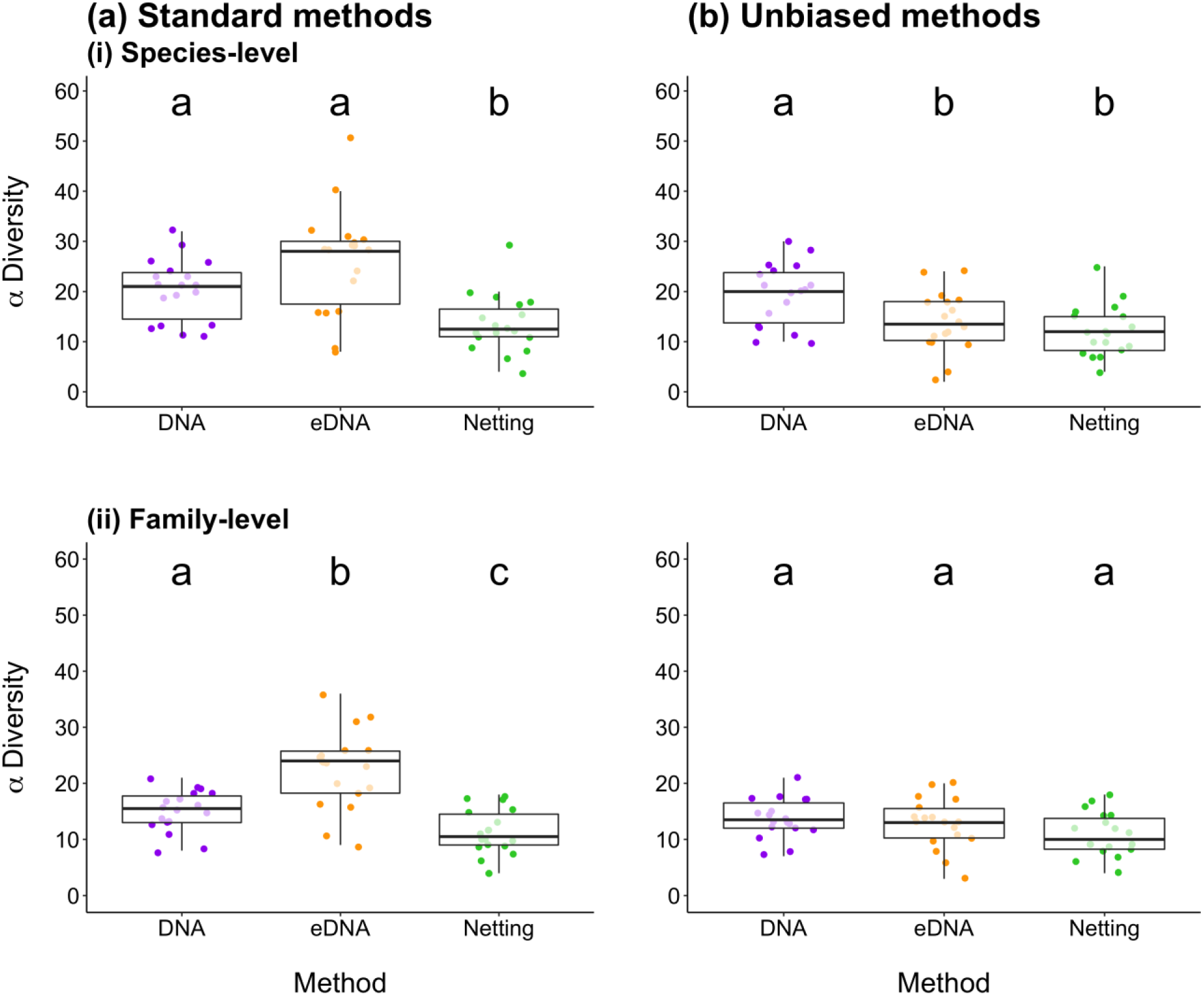
Mean alpha diversity of invertebrates in ponds across Norfolk and East Yorkshire. Alpha diversity is displayed according to standard **(a)** and unbiased **(b)** methods of invertebrate assessment. Species **(i)** and family **(ii)** richness generated by sweep-netting and microscopy (green points), DNA metabarcoding (purple points), and eDNA metabarcoding (orange points) is shown. Boxes show 25th, 50th, and 75th percentiles, and whiskers show 5th and 95th percentiles.

**Figure 3.**
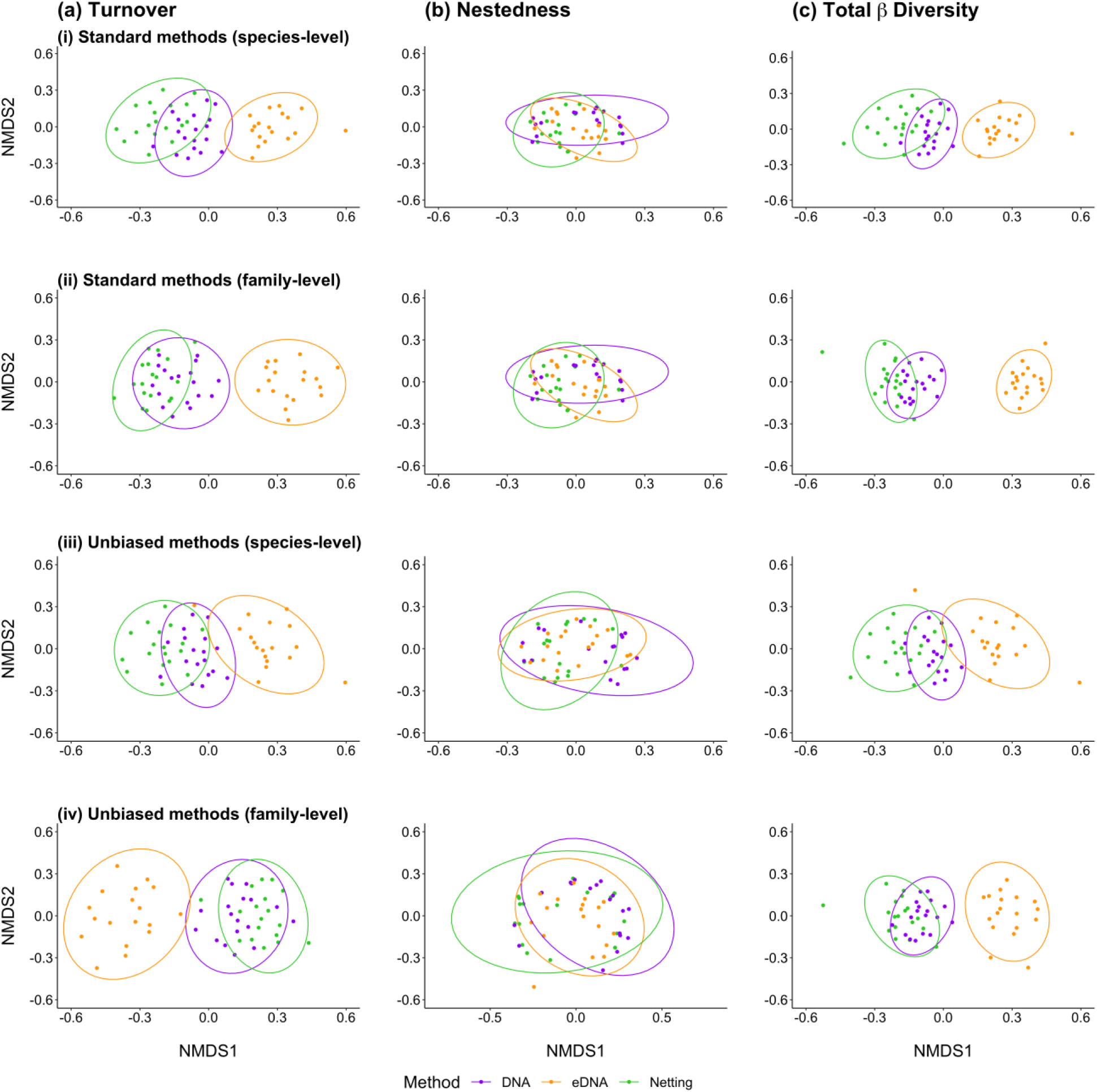
Non-metric Multidimensional Scaling (NMDS) plots of invertebrate communities (Jaccard dissimilarity) produced by sweep-netting and microscopy (green points/ellipse), DNA metabarcoding (purple points/ellipse), and eDNA metabarcoding (orange points/ellipse) for the 18 study ponds. The turnover **(a)** and nestedness-resultant **(b)** partitions of total beta diversity **(c)** are shown at species-level and family-level for standard **(i, ii)** and unbiased **(iii, iv)** methods.

### 3.2 Impact of crucian carp stocking on pond invertebrates

Here, we report the impact of crucian carp stocking assessed using standard methods as these provide the most comprehensive evaluation of invertebrate diversity. Crucian carp impact assessment using unbiased methods is reported in the Supporting Information (Appendix 2; Tables S10, S11, S12, S13; Figs. S11, S12, S13, S14). Independently and combined, standard methods showed alpha diversity of invertebrates to be marginally lower in ponds containing crucian carp at species (Figs. 4ai-iv) and family-level (Figs. 4bi-iv), but this difference was not statistically significant (*P* > 0.05, Table 2). Detailed examination of alpha diversity within the major invertebrate groups identified by all three methods combined revealed that Coleoptera and Mollusca diversity was reduced (Coleoptera borderline significant and Mollusca highly significant) in ponds with crucian carp at species-level (GLM χ^2^_21_ = 28.033, *P* = 0.139; Coleoptera -0.481 ± 0.254, *Z* = -1.897, *P* = 0.058; Mollusca -0.925 ± 0.278, *Z* = -3.324, *P* < 0.001), but not family-level (GLM χ^2^_22_ = 18.366, *P* = 0.684; Coleoptera -0.544 ± -0.296, *Z* = -1.834, *P* = 0.067; Mollusca -0.442 ± -0.302, *Z* = -1.462, *P* = 0.144). However, differences in alpha diversity between ponds with or without crucian carp were not significant for other invertebrate groups at either taxonomic rank (Fig. S15). Alpha diversity relationships held true for the unbiased methods (Table S10; Fig. S11, S12).

**Table 2.**
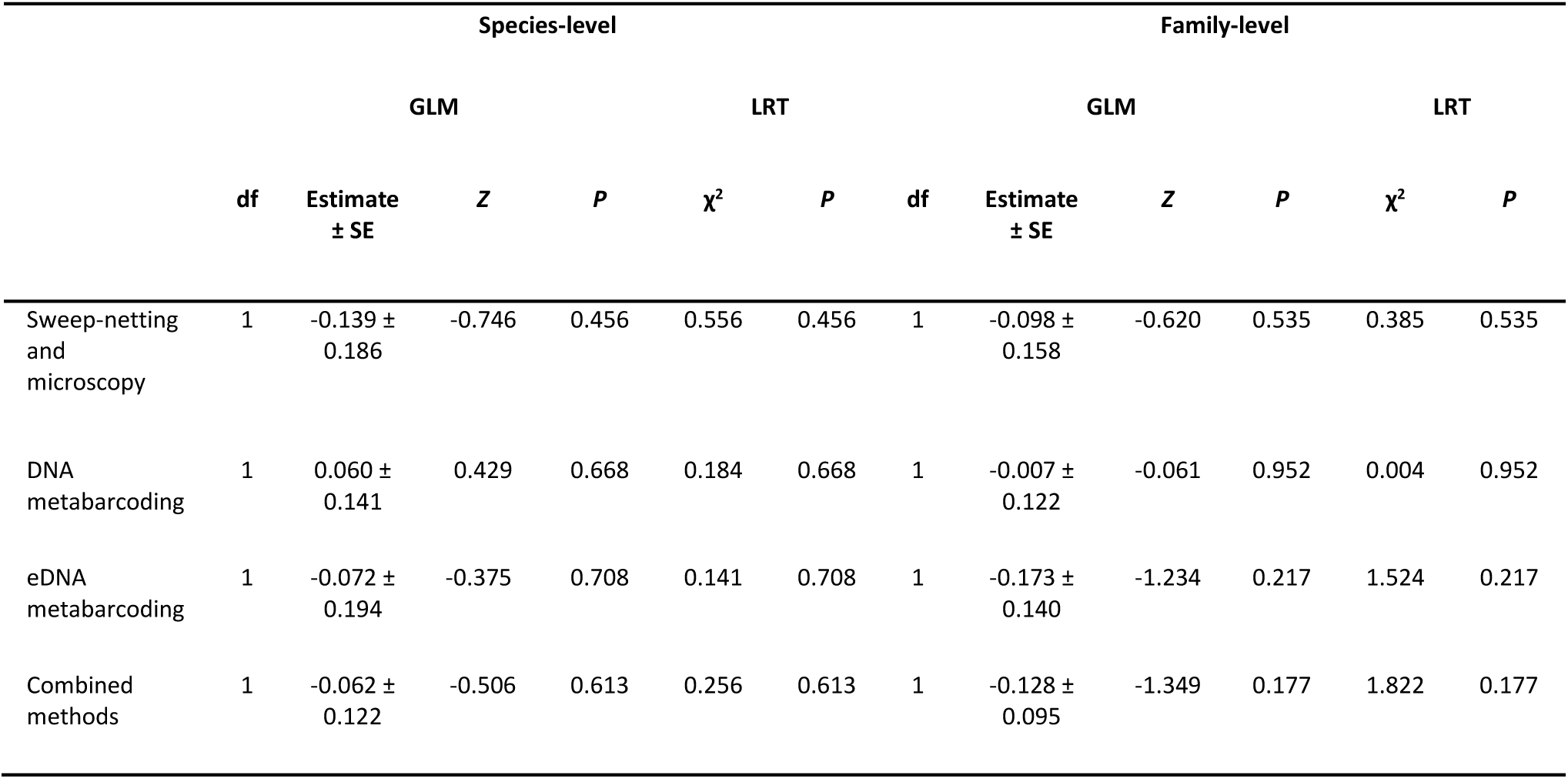
Summary of analyses (GLM) statistically comparing alpha diversity at species-level and family-level between ponds with and without crucian carp using independent and combined standard methods.

|  | Species-level |  |  |  |  |  | Family-level |  |  |  |  |  |
| --- | --- | --- | --- | --- | --- | --- | --- | --- | --- | --- | --- | --- |
|  | GLM |  |  | LRT |  |  | GLM |  |  | LRT |  |  |
| | df | Estimate<br>± SE | Z | P | $\chi^2$ | P | df | Estimate<br>± SE | Z | P | $\chi^2$ | P |
| Sweep-netting<br>and<br>microscopy | 1 | -0.139 ±<br>0.186 | -0.746 | 0.456 | 0.556 | 0.456 | 1 | -0.098 ±<br>0.158 | -0.620 | 0.535 | 0.385 | 0.535 |
| DNA<br>metabarcoding | 1 | 0.060 ±<br>0.141 | 0.429 | 0.668 | 0.184 | 0.668 | 1 | -0.007 ±<br>0.122 | -0.061 | 0.952 | 0.004 | 0.952 |
| eDNA<br>metabarcoding | 1 | -0.072 ±<br>0.194 | -0.375 | 0.708 | 0.141 | 0.708 | 1 | -0.173 ±<br>0.140 | -1.234 | 0.217 | 1.524 | 0.217 |
| Combined<br>methods | 1 | -0.062 ±<br>0.122 | -0.506 | 0.613 | 0.256 | 0.613 | 1 | -0.128 ±<br>0.095 | -1.349 | 0.177 | 1.822 | 0.177 |

**Figure 4.**
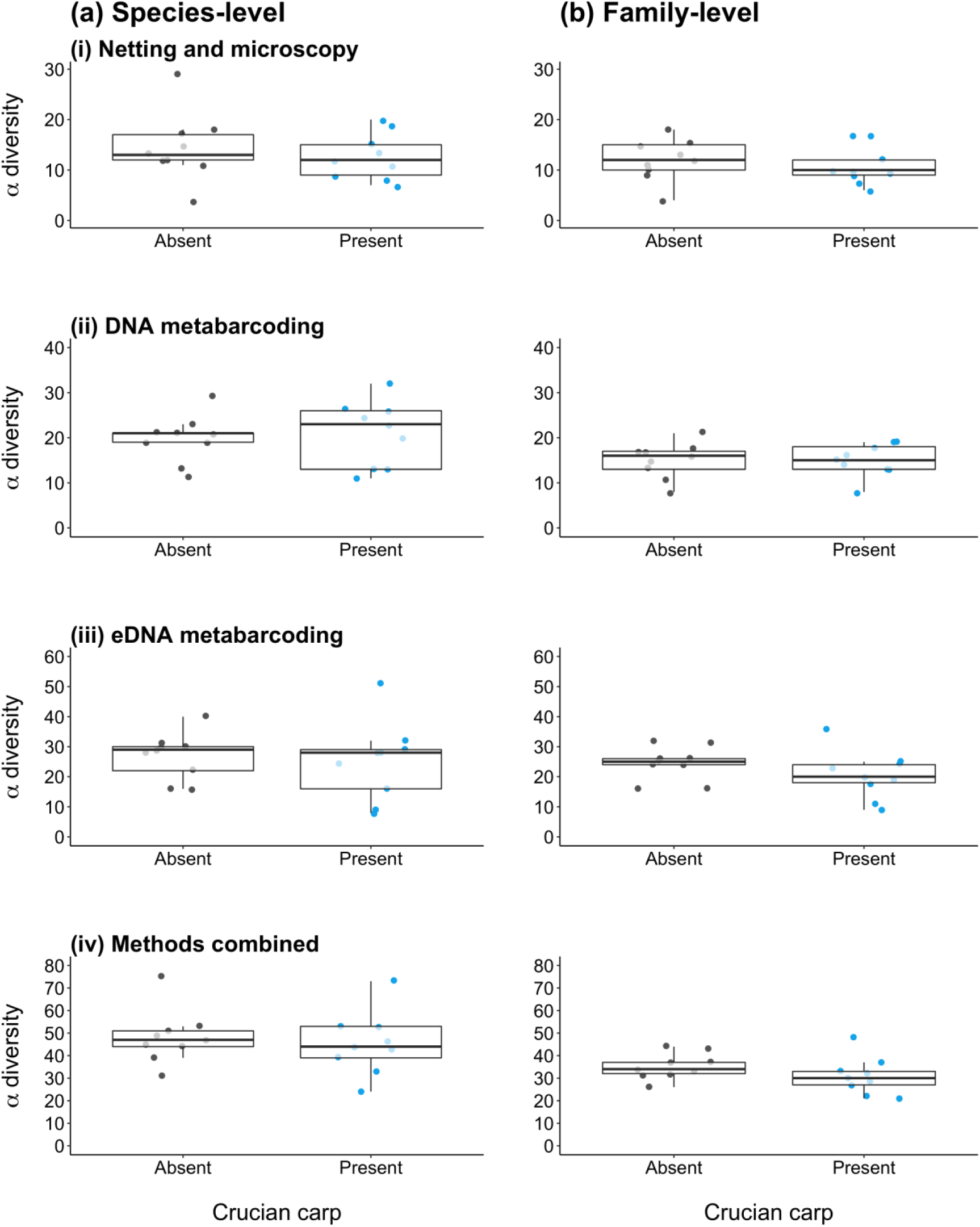
Mean alpha diversity of invertebrates in ponds with crucian carp (blue points) and without fish (grey points) across Norfolk and East Yorkshire. Alpha diversity at species-level **(a)** and family-level **(b)** is shown according to standard methods of invertebrate assessment: sweep-netting and microscopy **(i)**, DNA metabarcoding **(ii)**, eDNA metabarcoding **(iii)**, and all methods combined **(iv)**. Boxes show 25th, 50th, and 75th percentiles, and whiskers show 5th and 95th percentiles.

Total beta diversity of ponds was consistently high at species and family-level for independent and combined methods (standard or unbiased). Variation in invertebrate community composition was driven by turnover rather than nestedness-resultant (Tables 3, S11). MVDISP differed between ponds for species-level total beta diversity with eDNA metabarcoding, and species and family-level total beta diversity with all methods combined. Fishless ponds had significantly lower dispersion than ponds with crucian carp (Table S14). MVDISP patterns were not consistent across standard and unbiased methods (Table S12). At species-level, morphotaxonomic identification, eDNA metabarcoding, and all methods combined revealed a weak or moderate positive influence of crucian carp presence on turnover (Table 4; Figs. 5ai, iii, iv) and total beta diversity (Table 4; Figs. 5ci, iii, iv) between ponds, but not nestedness-resultant (Table 4; Figs. 5bi, iii, iv). At family-level, eDNA metabarcoding and all methods combined revealed a weak positive influence of crucian carp presence on total beta diversity (Table 4; Fig. S16ciii, iv) between ponds, but not turnover (Table 4; Fig. S16aiii, iv) or nestedness-resultant (Table 4; Fig. S16biii, iv). Crucian carp presence did not influence any beta diversity component produced by DNA metabarcoding (Table 4; Figs. 5aii, bii, cii). Similar beta diversity patterns were obtained using unbiased eDNA metabarcoding and combined methods (Table S13; Figs. S13, S14).

**Table 3.** Relative contribution of taxon turnover and nestedness-resultant to total beta diversity (Jaccard dissimilarity) using independent and combined standard methods. A value of 1 corresponds to all sites containing different species.

|  | Species-level |  |  | Family-level |  |  |
| --- | --- | --- | --- | --- | --- | --- |
|  | Turnover | Nestedness-resultant | Total beta diversity | Turnover | Nestedness-resultant | Total beta diversity |
| Sweep-netting and microscopy | 0.935<br>(98.01%) | 0.019<br>(1.99%) | 0.954<br>(100%) | 0.867<br>(94.75%) | 0.048<br>(5.25%) | 0.915<br>(100%) |
| DNA metabarcoding | 0.938<br>(98.53%) | 0.014<br>(1.47%) | 0.952<br>(100%) | 0.885<br>(96.83%) | 0.029<br>(3.17%) | 0.914<br>(100%) |
| eDNA metabarcoding | 0.917<br>(97.24%) | 0.026<br>(2.76%) | 0.943<br>(100%) | 0.871<br>(95.29%) | 0.043<br>(4.71%) | 0.914<br>(100%) |
| Combined methods | 0.927<br>(98.41%) | 0.015<br>(1.59%) | 0.942<br>(100%) | 0.871<br>(96.46%) | 0.032<br>(3.54%) | 0.903<br>(100%) |

**Table 4.** Summary of analyses (PERMANOVA) statistically examining variation in community composition between ponds with and without crucian carp at species-level and family-level using independent and combined standard methods.

| Community similarity (PERMANOVA) |  |  |  |  |  |  |  |  |
| --- | --- | --- | --- | --- | --- | --- | --- | --- |
|  | Species-level |  |  |  | Family-level |  |  |  |
|  | df | F | R<sup>2</sup> | P | df | F | R<sup>2</sup> | P |
| **Sweep-netting and microscopy** |  |  |  |  |  |  |  |  |
| Turnover | 1 | 1.673 | 0.095 | **0.018** | 1 | 1.198 | 0.070 | 0.358 |
| Nestedness-resultant | 1 | -3.454 | -0.275 | 0.949 | 1 | -0.103 | -0.007 | 0.620 |
| Total beta diversity | 1 | 1.369 | 0.079 | **0.033** | 1 | 1.136 | 0.066 | 0.306 |
| **DNA metabarcoding** |  |  |  |  |  |  |  |  |
| Turnover | 1 | 1.305 | 0.075 | 0.137 | 1 | 1.668 | 0.094 | 0.098 |
| Nestedness-resultant | 1 | -0.578 | -0.038 | 0.664 | 1 | -0.116 | -0.007 | 0.717 |
| Total beta diversity | 1 | 1.163 | 0.068 | 0.176 | 1 | 1.277 | 0.074 | 0.197 |
| **eDNA metabarcoding** |  |  |  |  |  |  |  |  |
| Turnover | 1 | 2.374 | 0.129 | **0.001** | 1 | 1.551 | 0.088 | 0.097 |
| Nestedness-resultant | 1 | -2.515 | -0.187 | 0.967 | 1 | 0.586 | 0.035 | 0.560 |
| Total beta diversity | 1 | 1.790 | 0.101 | **0.004** | 1 | 1.502 | 0.086 | **0.041** |
| **Combined methods** |  |  |  |  |  |  |  |  |
| Turnover | 1 | 1.999 | 0.111 | **0.001** | 1 | 1.452 | 0.083 | 0.136 |
| Nestedness-resultant | 1 | -1.663 | -0.116 | 0.966 | 1 | 3.103 | 0.162 | 0.128 |
| Total beta diversity | 1 | 1.717 | 0.097 | **0.001** | 1 | 1.474 | 0.084 | **0.027** |

**Figure 5.**
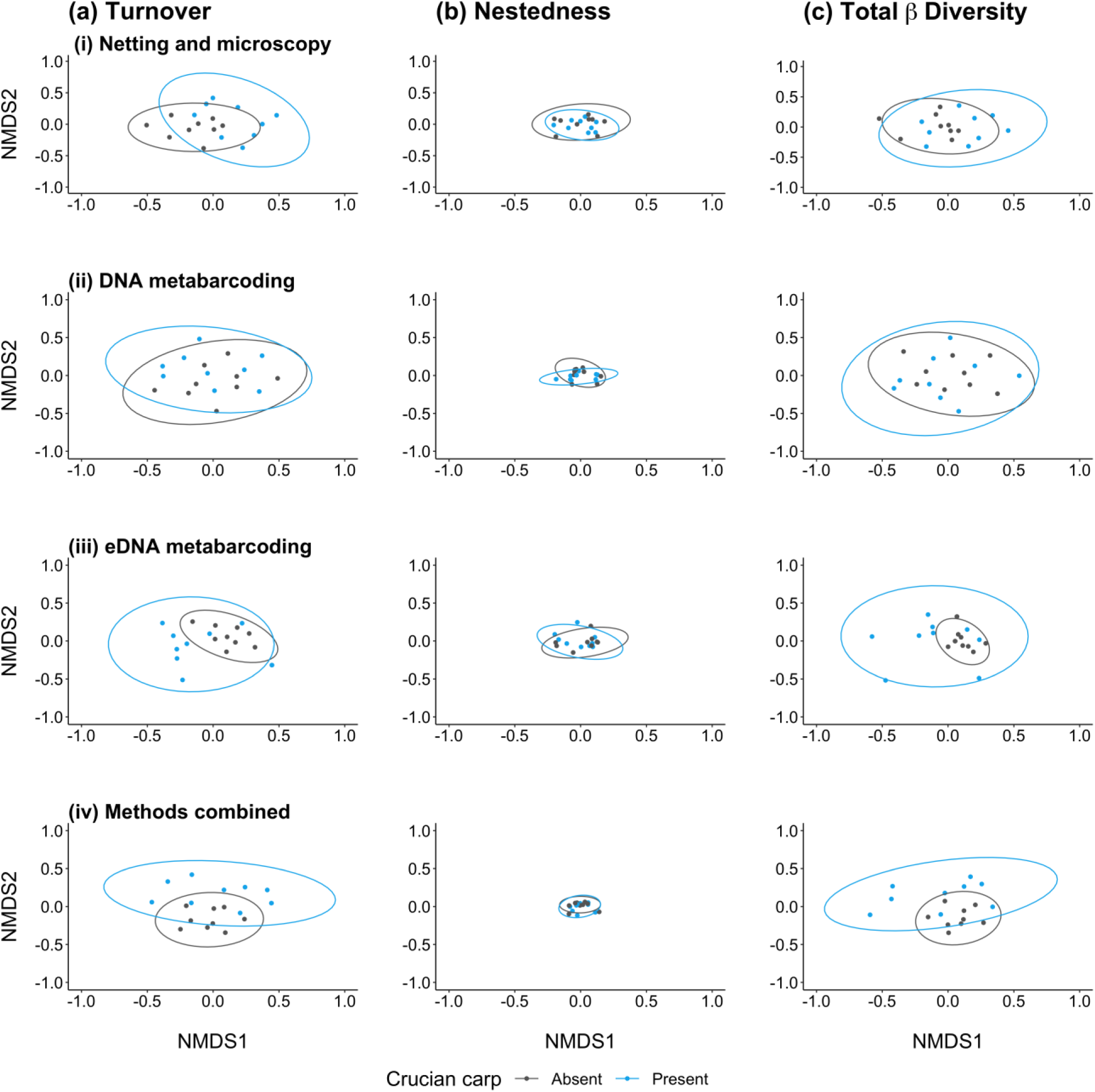
Non-metric Multidimensional Scaling (NMDS) plots of species-level invertebrate communities (Jaccard dissimilarity) from ponds with crucian carp (blue points/ellipse) and without fish (grey points/ellipse) across Norfolk and East Yorkshire. The turnover **(a)** and nestedness-resultant **(b)** partitions of total beta diversity **(c)** are shown according to standard methods of invertebrate assessment: netting and microscopy **(i)**, DNA metabarcoding **(ii)**, eDNA metabarcoding **(iii)**, and all methods combined **(iv)**.

Additional analyses undertaken on the combined method datasets supported an effect of crucian carp presence and excluded the influence of abiotic variables on invertebrate diversity. At species-level, only pond area was identified by forward selection as a significant abiotic variable for turnover and total beta diversity. Variance partitioning analysis was undertaken for turnover and total beta diversity using crucian carp presence-absence and pond area. Using the adjusted R^2^ values, biotic and abiotic variables explained 7.04% and 4.75% of the total variation in species turnover and total beta diversity respectively (Fig. 6). Crucian carp presence-absence significantly contributed to turnover (Fig. 6a; adjusted *R*^2^ = 5.55%, *F*_1_ = 2.031, *P* = 0.001) and total beta diversity (Fig. 6b; adjusted *R*^2^ = 4.05%, *F*_1_ = 1.730, *P* = 0.001), whereas pond area explained less variance and did not influence turnover (Fig. 6a; adjusted *R*^2^ = 2.94%, *F*_1_ = 0.340, *P* = 0.852) or total beta diversity (Fig. 6b; adjusted *R*^2^ = 1.93%, *F*_1_ = 1.118, *P* = 0.218). RDA of nestedness-resultant without abiotic data indicated no influence of crucian carp presence-absence (*F*_1_ = 0.340, *P* = 0.850). At family-level, forward selection did not identify any significant abiotic variables for turnover, nestedness-resultant, or total beta diversity. Therefore, variance partitioning was not undertaken for any component of beta diversity. RDA of each beta diversity component minus abiotic data showed that crucian carp presence-absence influenced total beta diversity (*F*_1_ = 1.474, *P* = 0.034), but not turnover (*F*_1_ = 1.399, *P* = 0.106) or nestedness-resultant (*F*_1_ = 1.970, *P* = 0.141). Variance partitioning results from the unbiased methods are presented in Appendix 2.

**Figure 6.**
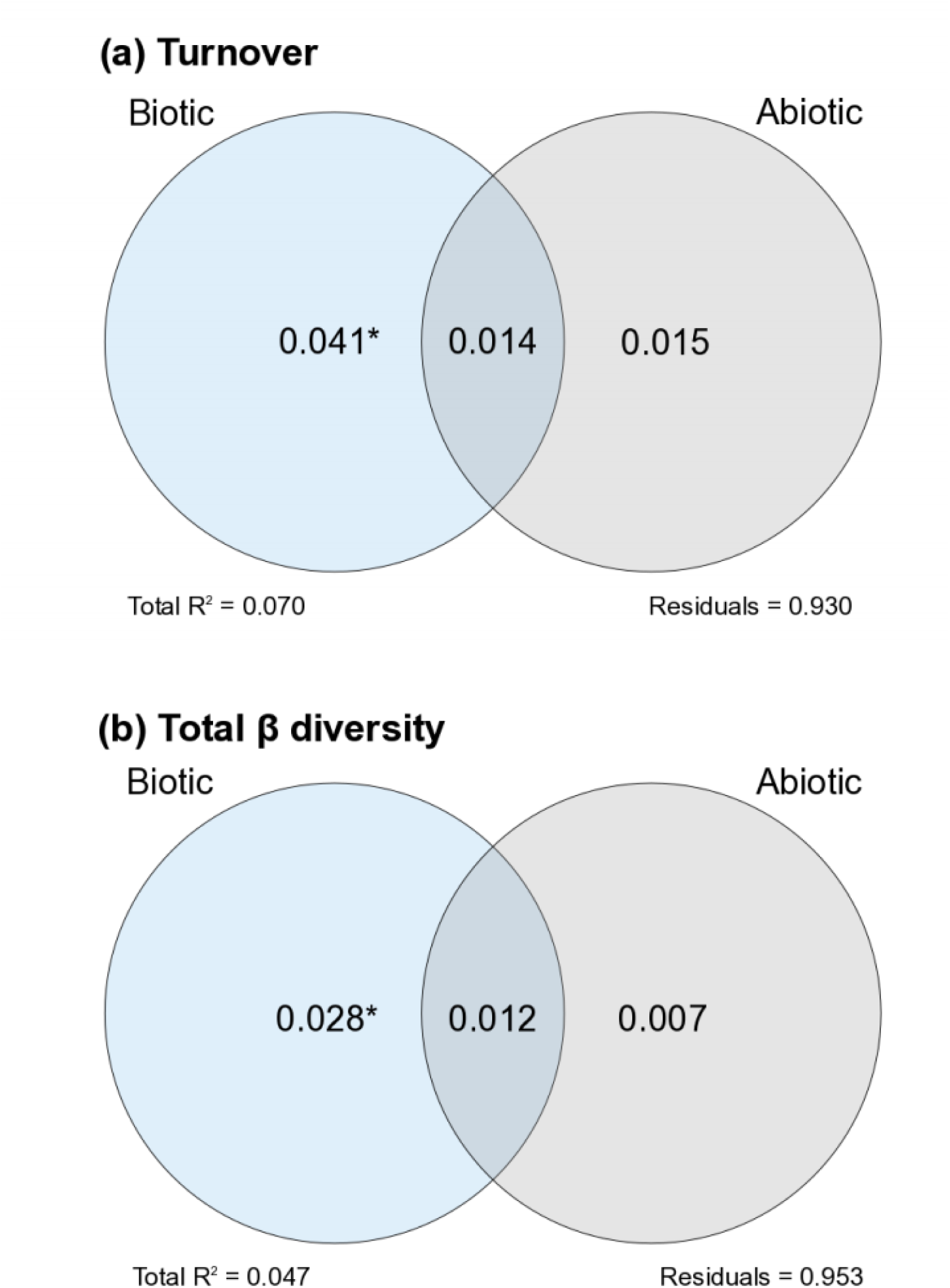
The relative contribution of biotic (crucian carp presence-absence) and abiotic variables (pond area) to species-level turnover **(a)** and total beta diversity **(b)** when the combined invertebrate data from all three standard sampling methods were considered. Values within circles and circle intersections represent the adjusted R^2^ values. Significant variables are indicated by asterisks (* = *P* < 0.05).

## 4. Discussion

### 4.1 Comparison of methods for freshwater invertebrate assessment

DNA (unbiased methods) or eDNA (standard methods) metabarcoding generated the highest species and family-level alpha diversity. Differences in alpha diversity were more pronounced between standard methods, but unique species were detected with unbiased methods nonetheless. Morphotaxonomic identification and DNA metabarcoding produced more similar communities, and performed best for Coleoptera, Hemiptera, Hirudinea, Megaloptera, and Odonata. eDNA metabarcoding generated a markedly different community, and detected aquatic taxa overlooked in pond net samples (Crustacea, Lepidoptera [i.e. Crambidae species with aquatic larvae such as *Cataclysta lemnata*], Trichoptera) alongside singleton terrestrial taxa (Arachnida, Hymenoptera, Psocoptera, Thysanoptera). Both metabarcoding approaches performed well for Annelida, Collembola, Diptera and Ephemeroptera, particularly at species-level.

Critically, we used a 1 mm mesh net which is designed to sample macroinvertebrates, whereas a 500 μm mesh net would also sample the meiofauna detected by metabarcoding. Meiofauna may inflate metabarcoding diversity estimates compared to morphotaxonomic identification, but may also prevent PCR amplification of macroinvertebrates (Beentjes et al., 2019; Deiner et al., 2016; Macher et al., 2018). To mitigate bias, we performed all analyses excluding meiofauna not captured by netting and excluding morphotaxonomically identified taxa without reference sequences that metabarcoding would not recover. Alpha diversity results from the standard and unbiased method comparisons were variable, but beta diversity results were more consistent. Both method comparisons echo other studies where metabarcoding captured greater diversity than morphotaxonomic approaches and resolved problematic groups (e.g. Diptera) for species-level morphotaxonomic identification (Andújar et al., 2017; Carew et al., 2018; Clarke, Beard, Swadling, & Deagle, 2017; Elbrecht et al., 2017a; Emilson et al., 2017; Klymus et al., 2017; Lobo et al., 2017). Standard and unbiased methods broadly revealed the same ecological relationships, although standard methods may yield more insights. Regardless, all three methods will be most powerful when used in combination for ecological investigation.

Each method has inherent biases that influence performance for freshwater invertebrate assessment. The UK National Pond Survey methodology recommends that pond net samples are placed into buckets/zip-lock bags, followed by sorting and identification in the laboratory as soon as possible. Samples are typically preserved in ethanol or refrigerated to prolong processing time (Biggs et al., 1998), but this standard methodology does not lend itself to molecular applications. After collection, samples were transported on ice then frozen at -20 °C to prevent predation within samples, minimise organismal decay and subsequent DNA degradation, and allow samples to be processed as required in the laboratory. Using this strategy, small or inconspicuous dead specimens would not have been recovered during sorting (Biggs et al., 1998) and therefore excluded from morphotaxonomic and DNA metabarcoding inventories. These losses are surplus to the estimated 29% of specimens already overlooked during sorting due to smaller body size (Haase et al., 2010). Sorting may also damage or completely destroy some recovered specimens, influencing morphotaxonomic identification (Lobo et al., 2017; Zizka, Leese, Peinert, & Geiger, 2018).

Human error is another source of discrepancy between morphotaxonomic identification and metabarcoding (Carew, Pettigrove, Metzeling, & Hoffmann, 2013; Elbrecht et al., 2017a; Haase et al., 2010). Some taxa were identified to genus or family-level and omitted from our species-level inventories, whereas other taxa may have been falsely identified and inventoried despite caution exercised to only assign taxa to species-level where there was high confidence in identification. Losses incurred by the preservation and sorting treatments for morphotaxonomic identification cannot be mitigated, and human error can only be reduced with taxonomic expertise. Conversely, taxon recovery by DNA metabarcoding can be improved with protocols that preserve and extract DNA from intact, unsorted samples (Elbrecht et al., 2017a; Nichols et al., 2020). For example, ethanol used to preserve specimens (Zizka et al., 2018), or replacement of ethanol with DNA extraction buffer followed by incubation then specimen removal (Carew, Coleman, & Hoffmann, 2018), offer alternative starting materials to bulk tissue for DNA metabarcoding. However, these alternative DNA sources tend to produce false negatives for schlerotised groups, such as Coleoptera, Trichoptera, and Mollusca (Carew et al., 2018; Zizka et al., 2018). More recently, Nichols et al. (2020) demonstrated that unsorted samples, including the sample matrix, recovered the same taxa as sorted samples, thus sorting may be unnecessary.

Despite successful application of DNA metabarcoding to bulk tissue samples for biomonitoring (Andújar et al., 2017; Carew et al., 2018; Elbrecht et al., 2017a; Emilson et al., 2017; Macher et al., 2018), recurring issues have been encountered. Arguably, the most pressing issue is size bias, where DNA from large and/or high biomass taxa can outcompete DNA of smaller and/or low biomass taxa during PCR amplification and sequencing (Elbrecht et al., 2017b). Although we used the size-sorting approach of Elbrecht *et al*. (2017b) and sequenced body size categories independently with data pooling downstream, DNA metabarcoding failed to detect several taxa which have reference sequences and were reliably identified by microscopy, including multiple coleopterans (*Agabus sturmii, Hydroporus erythrocephalu*s, *Rhantus frontalis, Haliplus confinis, Haliplus ruficollis*), a small mollusc (*Gyraulus crista*), a medium-sized hirudinean (*Erpobdella lineata*), and two large anisopterans (*Aeshna mixta, Anax imperator*). Primer bias may be partly responsible for these nondetections (see below), but non-delivery of desired results by size-sorting is problematic as it is a time-consuming, labour-intensive process that potentially increases cross-contamination risk between samples (Elbrecht et al., 2017b), and still requires samples to be sorted from vegetation and substrate (Elbrecht et al., 2017a). Size-sorting may therefore be a drain on resources and time allocated to DNA metabarcoding projects, but other sources of false negatives must be excluded.

In contrast, eDNA metabarcoding preferentially amplifies DNA from small, floating organisms (Beentjes et al., 2019; Deiner et al., 2016; Leese et al., 2020; Macher et al., 2018), and these organisms are often retained on filter membranes used for eDNA capture (*pers. obs*.). Consequently, their DNA is abundant in samples and overwhelms DNA from other taxa during sequencing. Pre-filtering with or only using large pore size filter membranes may reduce this bias (Macher et al., 2018). eDNA metabarcoding may also detect different taxa due to species variability in eDNA production and shedding rates. The species infrequently detected by eDNA metabarcoding were those that possess thicker exoskeletons composed of chitin and occasionally calcium carbonate, e.g. Coleoptera, Hemiptera. Exoskeletons may restrict the release of DNA into the water column (Tréguier et al., 2014) as opposed to organisms that are filter-feeders or produce slime (e.g. Mollusca), ectoparasites (i.e. Acari) that feed on blood or skin of other species, or use external versus internal fertilisation (Mächler et al., 2019; Wacker et al., 2019). Furthermore, species may differ in their habitat preferences and utilisation, potentially resulting in highly localised distributions (Klymus et al., 2017; Mächler et al., 2019). Therefore, more samples or greater volumes may be required to improve detection of pond biota (Harper et al., 2019).

Marker choice and primer design can substantially influence amplification success and taxonomic assignment. Although the *COI* region offers species resolution, has extensive database representation, and is used as standard in DNA barcoding (Elbrecht, Hebert, & Steinke, 2018), it lacks conserved primer-binding sites as a protein-coding gene (Clarke et al., 2017; Leese et al., 2020). This introduces high risk of primer mismatch and subsequent bias during metabarcoding (Clarke et al., 2017; Elbrecht et al., 2016; Elbrecht et al., 2018; Lobo et al., 2017). Most *COI* metabarcoding primers (Elbrecht & Leese, 2017; Leray et al., 2013; Meusnier et al., 2008; Zeale, Butlin, Barker, Lees, & Jones, 2011) are degenerate to enable binding, but degenerate primers may bind non-target regions (Elbrecht et al., 2018) and amplify non-metazoans, e.g. bacteria, algae, fungi (Brandon-Mong et al., 2015; Leese et al., 2020; Macher et al., 2018). Degenerate primers can experience primer slippage and produce variable length sequences across species, which has implications for bioinformatic processing and taxonomic assignment (Elbrecht et al., 2018). Using different (e.g. 16S ribosomal RNA) or multiple markers may counter these issues, but different markers often lack reference database representation, hindering taxonomic resolution, and use of multiple markers will increase PCR and sequencing costs (Clarke et al., 2017; Deiner et al., 2016; Elbrecht et al., 2016; Elbrecht et al., 2017a).

We obtained an excessive number (∼60%) of unassigned reads from eDNA metabarcoding, despite using a relaxed BLAST identity (90%) against custom and public reference databases. This suggests that sequences were of poor quality, could not be assigned to reference sequences, or lacked reference database representation (Macher et al., 2018). Public reference database records may be few, mislabelled, or have limited geographic coverage (Curry, Gibson, Shokralla, Hajibabaei, & Baird, 2018; Elbrecht et al., 2017a; Klymus et al., 2017; Weigand et al., 2019). We morphotaxonomically identified 19 species (Hemiptera, Mollusca, Odonata) without reference sequences whose DNA may have been sequenced, but would not have been taxonomically assigned to species-level. Future studies must develop more specific primers that have binding and flanking regions of high nucleotide diversity (Elbrecht et al., 2018; Leese et al., 2020) and/or target particular invertebrate orders or families (Klymus et al., 2017) as well as procure reference sequences for different markers (Curry et al., 2018; Elbrecht et al., 2016; Weigand et al., 2019). These are essential steps to improve the reliability and accuracy of molecular monitoring in freshwater ecosystems.

### 4.2 Impact of crucian carp stocking on pond invertebrates

Our findings indicate that the crucian carp can be an important driver of invertebrate community heterogeneity in ponds. Crucian carp had a negligible influence on alpha diversity and a positive influence on beta diversity of pond invertebrates. Alpha diversity in ponds with crucian carp was marginally reduced compared to alpha diversity in fishless ponds, but this difference was not significant across methods used at either taxonomic rank. Within the major invertebrate groups identified by all methods combined, species-level alpha diversity of Coleoptera and Mollusca was reduced in ponds containing crucian carp as opposed to fishless ponds. This may be because coleopterans are large, active swimmers that are visible and easy to hunt, and because molluscs are slow-moving prey. Across ponds, total beta diversity of invertebrate communities was driven by turnover (taxon replacement) rather than nestedness-resultant (taxon subsets) (Baselga & Orme, 2012). Our results revealed crucian carp to positively influence turnover and total beta diversity between ponds. Therefore, taxa in fishless ponds were replaced by different taxa in ponds with crucian carp, resulting in dissimilar community composition. Removal of top predators and dominant species, such as coleopterans, by crucian carp may allow other prey species to colonise but further research is needed to test this hypothesis.

Our results both echo and contradict Stefanoudis et al. (2017), where the presence of fish (including crucian carp) in ponds altered macrophyte and cladoceran community composition, but not water beetle composition. Hassall et al. (2011) also found that fish presence in ponds had a positive effect on species richness of most invertebrate taxa, but a negative effect on Coleoptera. Gee et al. (1997) observed no influence of fish stocking on macrophyte and macroinvertebrate species richness, although Odonata richness was lower and Trichoptera richness higher in stocked ponds. Conversely, other research found that managed/stocked ponds, some of which contained crucian carp, had lower invertebrate diversity than unmanaged sites (i.e. no human intervention to slow succession), which were characterised by Trichoptera, Coleoptera and Zygoptera larvae (Wood et al., 2001). Similarly, large, active and free-swimming taxa (Notonectidae, Corixidae, Gyrinidae, Dytiscidae, Aeshnidae, Libellulidae and Chaoboridae) were strongly associated with fish absence as well as more diverse and abundant in fishless lakes (Bendell & McNicol, 1995; Schilling et al., 2009a, 2009b).

Critically, only two of the aforementioned studies accounted for the identity of fish species assemblages present (Stefanoudis et al., 2017; Wood et al., 2001) and just one was based on standardised fish surveys (Stefanoudis et al., 2017), whereas other studies only considered the presence of a particular species (Schilling et al., 2009a) or fish presence-absence generally (Bendell & McNicol, 1995; Gee et al., 1997; Hassall et al., 2011; Schilling et al., 2009b). The contrasting results produced by these studies and our own indicate that the impact of fish on pond biodiversity will likely depend on the species present, population density, and management strategy. Wetland fishes vary in dietary preference and consume different proportions of invertebrate taxa, thus different fishes will suppress numbers of and confer benefits to different invertebrate taxa (Batzer, Pusateri, & Vetter, 2000). Invasive fish species may be more detrimental than non-invasive species, for example, the mosquitofish (*Gambusia affinis*) reduced zooplankton abundance and macroinvertebrate density by 90% and 50% respectively after introduction in a wetland ecosystem experiment (Preston et al., 2017). Additionally, invertebrate species richness and abundance was found to decrease as duration of stocking increased (Schilling et al., 2009a). Therefore, local-and regional-scale diversity may benefit most from pond mosaics that maintain fish-free and fish-containing (at low-moderate densities) ponds to prevent increases in fish biomass and predation pressure over time (Lemmens et al., 2013).

The effects of environmental variables on alpha and beta diversity of pond invertebrates must also be considered (Hassall et al., 2011; Hill, Heino, Thornhill, Ryves, & Wood, 2017). Our study ponds were selected to have similar physical and chemical properties, which may explain the minimal contribution of abiotic factors to variance in invertebrate community structure. Although pond area was retained by model selection, it did not significantly influence community structure and explained less variance than crucian carp presence-absence. Likewise, other studies have shown a weak or no effect of pond area on invertebrate species richness and community composition (Gee et al., 1997; Gledhill et al., 2008; Oertli et al., 2002; Wood et al., 2001). Critically, our environmental data were collected over a 7-year period, whereas contemporary data may have better explained variance in community structure. We omitted variables that experience high temporal variation from our analyses, but these may be linked to the Coleoptera and Mollusca alpha diversity reductions and different invertebrate communities observed in crucian carp ponds, such as water temperature, pH, nutrient concentration, and surface dissolved oxygen (Bendell & McNicol, 1995; Hassall et al., 2011; Menetrey, Sager, Oertli, & Lachavanne, 2005). Investigations examining contemporary physicochemical variables alongside crucian carp presence-absence will help disentangle the effects of stocking and habitat. The degree of connectivity between ponds may also substantially influence invertebrate community structure, both independently and in combination with crucian carp presence (Hill et al., 2017).

The crucian carp might be assumed to have negative impacts on pond biodiversity by having “carp” in its name and likely being non-native (Jeffries et al., 2017). Foraging activities of introduced common carp are known to reduce invertebrate density and macrophyte cover, with knock-on effects for amphibian and waterfowl species richness and abundance (Chan, 2011; Haas et al., 2007; Maceda-Veiga et al., 2017). However, all crucian carp ponds studied here were dominated by floating and/or submerged macrophyte beds as in Stefanoudis et al. (2017), and we observed no evidence for negative impacts on vegetation. Additionally, no impact of crucian carp on amphibian presence, oviposition, larval behaviour, or recruitment success has been found (Chan, 2011; Harper et al., 2019; Jarvis, 2012). Despite its probable non-native status, cumulative evidence suggests that the crucian carp does not have invasive potential. Instead, the crucian carp is naturalised and faces the same threats as native UK pond biodiversity, such as the great crested newt (*Triturus cristatus*), including terrestrialisation, dessication and infilling of ponds as well as predation by other fishes, e.g. pike (*Esox lucius*) and three-spined stickleback (*Gasterosteus aculeatus*) (Sayer et al., 2020). Additionally, the crucian carp is threatened by hybridisation with goldfish, common carp, and gibel carp across Europe (Hänfling et al., 2005). Gibel carp are though to be absent from UK waters and given the severe threat that this species poses to crucian carp persistence in ponds, England can be regarded as an important centre for crucian carp conservation (Jeffries et al., 2017; Sayer et al., 2020). Moreover, pond restoration as part of crucian carp conservation efforts in Norfolk has substantially benefitted macrophyte, invertebrate, amphibian, and farmland bird diversity (Sayer et al., 2013; Lewis-Phillips et al., 2019).

Our study demonstrates that crucian carp presence enhances invertebrate diversity across pond networks, and that current conservation management of established and stocked populations in England is appropriate. Crucian carp may contribute to taxon turnover and subsequent community heterogeneity in ponds by providing colonisation opportunities for smaller invertebrates through predator (e.g. coleopterans) removal. We conclude that management strategies to support high landscape-scale pond biodiversity should consider a patchwork of fish-containing and fishless ponds, although stocking should only occur in artificially created ponds or in ponds where fish were previously known to occur (Sayer et al., 2020). The crucian carp is likely to be an important species in this respect, but more research is required to verify that crucian carp presence increases community heterogeneity without contributing to the decline of endangered invertebrates, albeit this is unlikely. Furthermore, the impact of crucian carp on all pond biodiversity, especially amphibians, must be studied more broadly with respect to crucian carp population density, seasonality, and the aforementioned environmental variables to determine whether maintaining populations and stocking is invariably beneficial.

## Acknowledgements

This work was funded by the University of Hull. Additionally, D.S.R was supported by the Natural Environment Research Council award number NE/R016429/1 as part of the UK-SCAPE programme delivering National Capability. We thank the many landowners who allowed access to ponds. We are grateful to the Norfolk Crucian Project for providing data associated with Norfolk ponds. We thank Michael Lee and Matt Smith (Environment Agency) for selecting East Yorkshire ponds and providing associated data, and Jane Colley (Askham Bryan College) for providing invertebrate samples from these ponds. We are grateful to Jianlong Li and Cristina Di Muri (University of Hull) for assistance with water sampling and filtration as well as advice on library preparation and sequencing.

## Data accessibility

Raw sequence reads have been archived on the NCBI Sequence Read Archive (Study: SRP163672; BioProject: PRJNA494857; BioSamples: SAMN10181701 - SAMN10182084 [bulk tissue DNA] and SAMN10187732 - SAMN10188115 [eDNA]; SRA accessions: SRR7969394 - SRR796977 [bulk tissue DNA] and SRR7985814 - SRR7986197 [eDNA]). Jupyter notebooks, R scripts and corresponding data are deposited in a dedicated GitHub repository (https://github.com/HullUni-bioinformatics/Harper_et_al_2020_crucian_carp_impact_invertebrates_conventional_molecular_tools) which has been permanently archived (https://doi.org/10.5281/zenodo.3993125).

## Author contributions

L.R.H, B.H, and L.L.H conceived and designed the study. C.D.S selected the study ponds in Norfolk and provided associated environmental data. L.R.H, M.B, and R.C.B built the custom reference sequence database for invertebrates. L.R.H and R.C.B sampled the Norfolk ponds for invertebrates. L.R.H and M.B collected and filtered water samples from all study ponds. D.S.R assisted with water filtration of eDNA samples and provided advice on primers, PCR protocols, and sequencing. L.R.H performed laboratory work, and analysed the data with assistance from M.J.H. L.R.H wrote the manuscript, which all authors revised.

## Notes

https://doi.org/10.5281/zenodo.3993125

### Competing Interest Statement

The authors have declared no competing interest.

### Summary of Updates

This version of the manuscript has been revised for resubmission to a journal.

