## Supporting Information for "Assessing the impact of the threatened crucian carp (*Carassius carassius*) on pond invertebrate diversity - a comparison of conventional and molecular tools"

### Appendix 1: Materials and methods

#### 1.1 DNA reference database construction

Custom, phylogenetically curated reference databases were created for the mitochondrial cytochrome c oxidase subunit I (*COI*) region in UK invertebrate species, excluding Diptera. Public GenBank records for Diptera were missing record features (e.g. 'gene' or 'CDS') and/or names were not in the format required for custom reference database construction using the selected bioinformatic tools. A comprehensive list of recorded UK invertebrate species and their taxonomy was previously constructed by the Centre for Ecology and Hydrology (https://www.ceh.ac.uk/services/coded-macroinvertebrates-list). Database curation for each of the main invertebrate groups (e.g. Coleoptera, Odonata, Mollusca) was performed separately to ease data processing. Reference databases were constructed using ReproPhylo v1.3 (Szitenberg, John, Blaxter, & Lunt, 2015) in a Jupyter notebook (Jupyter Team, 2017). Jupyter notebooks detailing the processing steps for each data subset can be provided upon request.

Briefly, we used a BioPython script to perform a GenBank search based on the lists of binomial names and downloaded all available *COI* sequences for specified species. Where there were no records on GenBank for a UK species, the database was supplemented with downloaded sequences belonging to European sister species in the same genus. Species that had no *COI* records on Genbank are provided in Table S3. Redundant sequences were removed by clustering at 100% similarity using vsearch v1.1 (Rognes, Flouri, Nichols, Quince, & Mahé, 2016). Only sequences longer than 500 bp were processed to increase alignment robustness to large gaps. Sequences were aligned using MAFFT v7.123b (Katoh and Standley, 2013). Alignments were trimmed using trimAl v1.2 (Capella-Gutiérrez, Silla-Martínez, & Gabaldón, 2009), following which maximum likelihood trees were inferred with RAxML v8.0.12 (Stamatakis, 2006) using the GTR+gamma model of substitutions. The complete alignments were then processed using SATIVA v0.9-57-g8a99328 (Kozlov, Zhang, Yilmaz, Glöckner, & Stamatakis, 2016) for automated identification of 'mislabelled' sequences which could cause conflict in downstream analyses. Putatively mislabelled sequences were removed, whereupon alignment and phylogenetic tree construction were repeated for manual investigation of sequences.

The resultant databases (i.e. curated non-redundant reference databases) contained sequences from: 412/423 Coleoptera species, 54/59 Odonata species, 83/92 Ephemeroptera/Plecoptera/Nemoptera/Megaloptera species, 187/206 Trichoptera/Lepidoptera species, 53/114 Hemiptera/Hymenoptera species, 154/388 Crustacea species, 78/111 Mollusca species, 333/333 Arachnida species, and 129/152 Annelida species. The databases were supplemented by Sanger sequences obtained from tissue of a two-spotted assassin bug (*Platymeris biguttatus*) housed at the University of Hull. DNA from this species was used as our PCR positive control. Two-spotted assassin bug DNA was extracted from tissue samples using a DNeasy Blood & Tissue Kit (Qiagen®, Hilden, Germany). Reference sequences were generated using the standard *COI* primers for DNA barcoding of invertebrates (Folmer, Black, Hoeh, Lutz, & Vrijenhoek, 1994). PCR reactions were performed in 25 µL volumes containing 12.5 µL of MyTaq™ HS Red Mix (Bioline Reagents Limited, London, UK), 1 µL (10 µM, final concentration 0.04 µM) of forward and reverse primer (Integrated DNA Technologies, Belgium), 8.5 µL of molecular grade sterile water (Fisher Scientific UK Ltd, Loughborough, UK) and 2 µL DNA template. PCRs were performed on an Applied Biosystems® Veriti Thermal Cycler (Fisher Scientific UK Ltd, Loughborough, UK) with the following profile: 94°C for 3 min, 37 cycles of 94°C for 30 sec, 52°C for 60 sec and 72°C for 90 sec, followed by a final elongation step at 72°C for 10 min. Purified PCR products were Sanger sequenced directly (Macrogen Europe, Amsterdam, Netherlands) in both directions using the PCR primers. Sequences were edited using CodonCode Aligner (CodonCode Corporation, Centerville, MA, USA). The complete invertebrate reference databases compiled in GenBank or fasta format have been deposited in the GitHub repository for this study (https://github.com/HullUni-bioinformatics/Harper_et_al_2020_crucian_carp_impact_invertebrates_conventional_molecular_tools) which has been permanently archived (https://doi.org/10.5281/zenodo.3993125). These databases were used for *in silico* validation of metabarcoding primers.

#### 1.2 Primer validation

Invertebrate DNA from bulk tissue and environmental DNA (eDNA) samples was amplified with published *COI* metabarcoding primers, mICOIintF GGWACWGGWTGAACWGTWTAYCCYCC (Leray et al., 2013) and jgHCO2198 TAIACYTCIGGRTGICCRAARAAYCA (Geller, Meyer, Parker, & Hawk, 2013). These primers were validated for the present study *in silico* using ecoPCR software (Ficetola et al., 2010) against the custom reference databases for UK invertebrates. Parameters were set to allow a fragment size of 250-350 bp and maximum of three mismatches between each primer and each sequence in the reference database.

An additional five ponds (four with crucian carp and one fishless) in Norfolk were sampled for invertebrates in accordance with the UK National Pond Survey methodology (Biggs, Fox, & Nicolet, 1998) to obtain 38 macroinvertebrate species that represented 38 families and 10 major groups for *in vitro* primer validation. Specimens of newly inventoried species were removed for individual preservation in sterile 2 mL microtubes (Fisher Scientific UK Ltd, UK) with 100% ethanol, and stored at -20°C until DNA extraction. Each species was extracted individually, using a leg or foot as starting tissue material, with the DNeasy Blood & Tissue Kit (Qiagen®, Hilden, Germany) following the manufacturer's protocol. DNA extracts were stored at -20°C until PCR. These five ponds were not included for DNA metabarcoding.

During *in vitro* testing, mICOIintF/jgHCO2198 were compared to two other published primer sets for macroinvertebrate DNA metabarcoding: BF2/BR2 (Elbrecht & Leese, 2017) and fwhF1/fwhR1 (Vamos, Elbrecht, & Leese, 2017). Primer validation tests were performed at University of Hull in a separate laboratory situated on a different floor to the dedicated eDNA laboratory. All PCR reactions were performed in 25 µL volumes containing 12.5 µL of MyTaq™ HS Red Mix (Bioline Reagents Limited, London, UK), 1 µL (10 µM, final concentration 0.04 µM) of forward and reverse primer (Integrated DNA Technologies, Belgium), 8.5µL of molecular grade sterile water (Fisher Scientific UK Ltd, Loughborough, UK) and 2 µL DNA template. Thermocycling conditions were kept as consistent as possible across the different primer sets tested, bar annealing temperature. PCRs were performed on an Applied Biosystems® Veriti Thermal Cycler (Fisher Scientific UK Ltd, Loughborough, UK) with the following profile: 95°C for 3 min, 35 cycles of 95°C for 30 sec, 50-52°C for 30 sec, and 72°C for 60 sec, followed by a final elongation step at 72°C for 10 min. Annealing temperatures for mICOIintF/jgHCO2198, BF2/BR2, and fwhF1/fwhR1 were 51°C, 50°C, and 52°C respectively. Molecular grade sterile water (Fisher Scientific UK Ltd, Loughborough, UK) substituted template DNA for PCR negative controls.

#### 1.3 Sweep-net samples

The time allotted to sweep-netting was divided equally across identified mesohabitats (e.g. emergent macrophytes, submerged macrophytes, shaded water, marginal grasses, open water) unless ponds contained dominant mesohabitat. More sampling time was apportioned to dominant mesohabitat, i.e. time was quartered in a pond with one dominant and two subsidiary mesohabitats (Biggs et al., 1998). During the 1 min search, the water surface and hard substrate (e.g. rocks, logs) were inspected for aquatic invertebrates additional to those collected in the net. Collected material from sweeps and searches were pooled to create one sample for each pond, and deposited in a 1.2 L sterile Whirl-Pak® stand-up bag (Cole-Palmer, Hanwell, London).

#### 1.4 eDNA samples

Five 2 L surface water samples were collected from the shoreline of each pond using sterile Gosselin™ HDPE plastic bottles (Fisher Scientific UK Ltd, UK) and disposable gloves. Samples were taken at equidistant points around the pond perimeter where access permitted. All ponds without crucian carp were sampled on 22 August 2016. Water samples were transported on ice in sterile coolboxes to the Centre for Ecology & Hydrology (CEH), Wallingford, stored at 4°C, and vacuum-filtered within 24 hours of collection. Coolboxes were sterilised using 10% v/v chlorine-based commercial bleach solution (Elliot Hygiene Ltd, UK) and 70% v/v ethanol solution before ponds containing crucian carp were sampled on 25th August 2016. Samples were handled in the same way. For each pond, a full process blank (1 L molecular grade water) was taken into the field and stored in coolboxes with samples. Blanks were filtered and extracted alongside samples to identify contamination.

Where possible, the full 2 L of each sample was vacuum-filtered through sterile 0.45 µm cellulose nitrate membrane filters with pads (47 mm diameter; Whatman, GE Healthcare, UK) using Nalgene filtration units. One hour was allowed for each sample to filter but if filters clogged during this time, a second filter was used. After 2 L had been filtered or one hour had passed, filters were removed from pads using sterile tweezers and placed in sterile 47 mm petri dishes (Fisher Scientific UK Ltd, UK), which were sealed with parafilm (Sigma-Aldrich®, UK) and stored at -20°C. After each round of filtration (samples and blanks from two ponds), all equipment was sterilised in 10% v/v chlorine-based commercial bleach solution (Elliot Hygiene Ltd, UK) for 10 minutes, immersed in 5% v/v MicroSol detergent (Anachem, UK), and rinsed with purified water.

All filters were transported on ice in a sterile coolbox to the University of Hull and stored at -20°C until DNA extraction one week later. DNA was isolated from filters using the PowerWater® DNA Isolation Kit (MO BIO Laboratories, CA, USA), following the manufacturer's protocol in a dedicated eDNA facility at the University of Hull. This facility is devoted to pre-PCR processes with separate rooms for filtration, DNA extraction, and PCR preparation of environmental samples. Duplicate filters from the same sample were co-extracted by placing both filters in a single tube for bead milling. Eluted DNA (100 µL) concentration was quantified on a Qubit™ 3.0 fluorometer using a Qubit™ dsDNA HS Assay Kit (Invitrogen, UK). DNA extracts were stored at -20°C until further analysis.

#### 1.5 Metabarcoding workflow

A two-step PCR protocol was performed on bulk tissue DNA and eDNA samples at the University of Hull. For bulk tissue DNA samples, PCR reactions were set up in a UV and bleach sterilized laminar flow hood in a laboratory for analysis of tissue DNA with separate rooms for pre-PCR and post-PCR processes. eDNA samples were processed in the dedicated eDNA facility at the University of Hull with separate rooms for filtration, DNA extraction and PCR preparation of sensitive environmental samples. PCR reactions of eDNA samples were also set up in a UV and bleach sterilized laminar flow hood. For both sample types, eight-strip PCR tubes with individually attached lids were used instead of 96-well plates to minimise cross-contamination risk between samples (Port et al., 2016). PCR positive and negative controls were included on each PCR run (typically two positive and negative controls on each 96-well run), to screen for sources of potential contamination. The DNA used for the PCR positive control (tissue DNA $N$ = 9, eDNA $N$ = 11) was assassin bug (*Platymeris biguttatus*), as this is an exotic, terrestrial species not found in the UK whose DNA had not been handled in our laboratory prior to this study. The negative controls (tissue DNA $N$ = 9, eDNA $N$ = 11) substituted molecular grade sterile water (Fisher Scientific UK Ltd, Loughborough, UK) for template DNA.

During the first PCR, the target region was amplified using oligonucleotides including the locus primers described above, sequencing primers, and pre-adapters (Illumina, 2011). First PCR reactions were performed in triplicate in a final volume of 20 µL, using 2 µL of DNA extract as a template. The amplification mixture contained 10 µL of MyTaq™ HS Red Mix (Bioline Reagents Limited, London, UK), 1 µL (10 µM, final concentration 0.5 µM) of forward and reverse primer (Integrated DNA Technologies, Belgium) and 6 µL of molecular grade sterile water (Fisher Scientific UK Ltd, Loughborough, UK). PCRs were performed on an Applied Biosystems® Veriti Thermal Cycler (Fisher Scientific UK Ltd, Loughborough, UK) and PCR conditions for the first component of the two-step protocol consisted of: an incubation step at 95°C for 3 min, followed by 35 cycles of denaturation at 95°C for 30 s, annealing at 47°C for 30 s, and extension at 72°C for 1 min, with final extension at 72°C for 10 min. PCR products were stored at 4°C until fragment size was verified by visualising 2 µL of selected PCR products on 2% agarose gels (80 mL 1x Sodium Borate buffer (Brody & Kern, 2004), 1.6 g agarose powder). Gels were stained with ethidium bromide or GelRed® (Cambridge BioScience, UK) and imaged using Image Lab Software (Bio-Rad Laboratories Ltd, Watford, UK). A PCR product was deemed positive where there was an amplification band on the gel that was of the expected size (400-500 bp). PCR replicates for each eDNA sample were pooled in preparation for the addition of Illumina indexes in the second PCR, which resulted in 60 µL of PCR product for each sample. PCR replicates for bulk tissue samples, PCR positive controls, and PCR negative controls were not pooled to allow individual purification and sequencing. All PCR products were purified to remove excess primer using magnetic bead clean-up. Mag-Bind® RxnPure Plus beads (Omega Bio-tek, GA, USA) were used while following a double size selection protocol from Bronner *et al*. (2009). Magnetic bead ratios of 0.5x and 0.12x to 20 µL of first PCR product were used. Eluted DNA (15 µL) was stored at -20 °C until the second PCR could be performed.

In the second PCR, Multiplex Identification (MID) tags (unique 8-nucleotide sequences) and Illumina MiSeq adapter sequences were bound to the amplified product. For each second PCR run, 96 unique tag combinations were created by combining eight unique forward tags with 12 unique reverse tags or vice versa (Kitson et al., 2019). A total of 384 unique tag combinations were achieved, allowing samples to be distinguished during bioinformatics analysis. Second step PCR reactions were performed in eight-strip PCR tubes with individually attached lids in a final volume of 50 µL, using 5 µL of purified DNA from the first PCR product as a template. The amplification mixture contained 25 µL of MyTaq™ HS Red Mix (Bioline Reagents Limited, London, UK), 5 µL (10 µM, final concentration 0.4 µM) of tagged primer mix (Integrated DNA Technologies, Belgium) and 15 µL of molecular grade sterile water (Fisher Scientific UK Ltd, Loughborough, UK). PCR was performed on an Applied Biosystems® Veriti Thermal Cycler (Fisher Scientific UK Ltd, Loughborough, UK) with the following profile: denaturation at 95°C for 3 min, followed by 8 cycles of denaturation at 95°C for 15 s, annealing at 55°C for 30 s, and extension at 72°C for 30 s, with final extension at 72°C for 10 min. PCR products were stored at 4°C before they were visualised on 2% agarose gels (80 mL 1x Sodium Borate buffer, 1.6 g agarose powder) using 2 µL PCR product. Gels were stained with ethidium bromide or GelRed® (Cambridge BioScience, UK) and imaged using Image Lab Software (Bio-Rad Laboratories Ltd, Watford, UK). Again, PCR products were deemed positive where there was an amplification band on the gel that was of the expected size (500-600 bp). Second PCR products (25 µL) were pooled according to PCR run, and the pooled PCR products purified to remove excess primer using magnetic bead clean-up. Mag-Bind® RxnPure Plus beads (Omega Bio-tek, GA, USA) were used while following a double size selection protocol from Bronner *et al*. (2009). Magnetic bead ratios of 0.5x and 0.12x to 200 µL of pooled PCR product were used. Eluted DNA (30 µL) was stored at -20°C until library quality control.

The pooled PCR products were quantified on a Qubit™ 3.0 fluorometer using a Qubit™ dsDNA HS Assay Kit (Invitrogen, UK) and pooled proportional to sample size and concentration. The pooled libraries were then quantified on a Qubit™ 3.0 fluorometer using a Qubit™ dsDNA HS Assay Kit (Invitrogen, UK). Based on Qubit™ concentration, the libraries were diluted to 4 nM for quantification by real-time quantitative PCR (qPCR) using the NEBNext® Library Quant Kit for Illumina® (New England Biolabs® Inc., MA, USA). The libraries were also checked using an Agilent 2200 TapeStation and High Sensitivity D1000 ScreenTape (Agilent Technologies, CA, USA) to verify secondary product had been removed successfully and a fragment of the expected size (531 bp) remained. The bulk tissue library was sequenced at 15 pM with 10% PhiX Control and eDNA library sequenced at 8pM with 20% PhiX Control on an Illumina MiSeq® using 2 x 300 bp V3 chemistry (Illumina Inc., CA, USA).

Illumina data was converted from raw sequences to taxonomic assignment using a custom pipeline for reproducible analysis of metabarcoding data: metaBEAT (metaBarcoding and eDNA Analysis Tool) v0.97.11 (https://github.com/HullUni-bioinformatics/metaBEAT). Raw reads were quality trimmed using Trimmomatic v0.32 (Bolger, Lohse, & Usadel, 2014), both from the read ends (minimum per base phred score Q30), as well as across sliding windows (window size 5bp; minimum average phred score Q30). Reads were clipped to a length of 200 bp and reads shorter than 200 bp after quality trimming were discarded. To reliably exclude adapters and PCR primers, the first 26 bp of all remaining reads were also removed. Sequence pairs were merged into single high quality reads using FLASH v1.2.11 (Magoč & Salzberg, 2011), if a minimum of 10 bp overlap with a maximum of 10% mismatch was detected between pairs. For reads that were not successfully merged, only forward reads were kept. To reflect our expectations with respect to fragment size, a final length filter was applied and only sequences of length 313 bp were retained. These were screened for chimeric sequences against our custom reference database using the uchime algorithm (Edgar, Haas, Clemente, Quince, & Knight, 2011), as implemented in vsearch v1.1.0 (Rognes et al., 2016). Redundant sequences were removed by clustering at 97% identity ('--cluster_fast' option) in vsearch v1.1.0 (Rognes et al., 2016). Clusters represented by less than three sequences were considered sequencing error and omitted from further analyses. Non-redundant sets of query sequences were then compared against our custom reference database using BLAST (Zhang, Schwartz, Wagner, & Miller, 2000). For any query matching with at least 90% identity to a reference sequence across more than 80% of its length, putative taxonomic identity was assigned using a lowest common ancestor (LCA) approach based on the top 10% BLAST matches. Sequences that could not be assigned (non-target sequences) were subjected to a separate BLAST search against the complete NCBI nucleotide (nt) database at 90% identity to determine the source via LCA as described above. To ensure reproducibility of analyses, the described workflow has been deposited in the GitHub repository.

#### 1.6 Data manipulation and analysis

Total read counts per sample were calculated and retained for false positive threshold determination. Assignments from the custom and public databases were merged, and spurious assignments (i.e. non-metazoans) removed from the datasets. Assignments corresponding to ambiguous BOLD records were renamed as the genus or family stated in the record name. We identified several taxa that had been misassigned based on absence of UK occurrence records in public databases (NBN Atlas, GBIF, UK Natural History Museum). *Erythromma humerale* was detected by the BLAST of unassigned reads from bulk tissue DNA metabarcoding against the entire NCBI nucleotide database. This species was corrected to *Erythromma* as both *E. najas* and *E. viridulum* occur in the UK but *E. viridulum* does not have reference sequences. *Radix peregra* was identified by microscopy using standard keys, but has recently been reclassified as *Radix balthica* which was identified by DNA and eDNA metabarcoding. Similarly, *Gyraulus crista* was identified by microscopy using standard keys, but DNA and eDNA metabarcoding identified its synonym, *Armiger crista*. Therefore, we corrected *Radix peregra* to *Radix balthica* and *Gyraulus crista* to *Armiger crista* in the morphotaxonomic dataset to match the metabarcoding datasets.

Other misassignments were species that did not occur in the UK, but had sister species in the same genus that do occur in the UK. These misassignments largely belonged to Gastrotricha, Rotifera and Annelida and were as follows: *Abacarus lolii* reassigned to *Abacarus*, *Bichromomyia flaviscutellata* reassigned to Psychodidae, *Callicorixa audeni* reassigned to *Callicorixa*, *Chaetonotus aemilianus* reassigned to *Chaetonotus*, *Chaetonotus antrumus* reassigned to *Chaetonotus*, *Chaetonotus daphnes* reassigned to *Chaetonotus*, *Chironomus muratensis* reassigned to *Chironomus*, *Chydorus brevilabris* reassigned to *Chydorus*, *Enochrus ater* reassigned to *Enochrus*, *Glossiphonia concolor* reassigned to *Glossiphonia*, *Haplothrips tenuipennis* reassigned to *Haplothrips*, *Helobdella modesta* reassigned to *Helobdella*, *Heterolepidoderma macrops* reassigned as *Heterolepidoderma*, *Kurzia media* reassigned to *Kurzia*, *Mimeoma maculata* reassigned to Scarabaeidae, *Parachironomus major* reassigned to *Parachironomus*, *Stenostomum sthenum* reassigned to *Stenostomum*, *Stylochaeta scirtetica* reassigned to *Stylochaeta*, and *Thermonectus* reassigned to Dytiscidae.

Reads from the same assignments were then merged. To minimise risk of false positives, taxa were only classed as present in samples if their sequence frequency exceeded set thresholds. The DNA metabarcoding dataset threshold was defined using the maximum sequence frequency of the PCR positive control (assassin bug DNA) in the community DNA samples (0.0236%). However, there was no assassin bug contamination of eDNA samples, thus taxon-specific thresholds were applied to the eDNA metabarcoding dataset instead (Harper et al., 2018). These thresholds were defined using the maximum sequence frequency of each taxon in the PCR positive controls ($N$ = 11). Only *Homo sapiens* (0.0753%) required a threshold. After threshold application, non-invertebrate assignments and invertebrate assignments above family-level were removed. Semi-aquatic and terrestrial taxa detected by eDNA metabarcoding were not removed before downstream analyses as these taxa signify the high sensitivity of this approach for biomonitoring. For the DNA metabarcoding dataset, we pooled the sequence data for PCR/sequencing replicates belonging to the same size category, and pooled size categories according to pond sampled. For the eDNA metabarcoding dataset, we pooled the sequence data for biological replicates belonging to the same pond.

### Appendix 2: Results

#### 2.1 Primer validation

The *in silico* analysis indicated poor taxonomic coverage and resolution of the *COI* primers, where only 9.24% of target invertebrate species amplified. A small range of UK invertebrate taxa were amplified, with fragment length ranging from 307-313 bp. The primers amplified 18/78 Coleoptera species, 16/54 Odonata species, 9/83 Ephemeroptera/Plecoptera/Nemoptera/Megaloptera species, 8/187 Trichoptera/Lepidoptera species, 4/53 Hemiptera/Hymenoptera species, 20/154 Crustacea species, 18/78 Mollusca species, 10/333 Arachnida species, and 29/129 Annelida species (Table S3; Fig. S2). However, an important caveat of these results is available reference sequences on GenBank (Fig. S3). The majority of invertebrate *COI* sequences on GenBank were generated using the Folmer primers, LCO1490 and HCO2198 (Folmer et al., 1994). After Sanger sequencing, primer regions are often removed due to low quality sequence produced at the start of sequencing. Therefore, primer sequences are typically not included in invertebrate reference sequences uploaded to GenBank. Our forward metabarcoding primer mICOIintF (Leray et al., 2013) lies within the 658 bp fragment amplified by the Folmer primers. However, our reverse metabarcoding primer jgHCO2198, a modified version of HCO2198 (Geller et al., 2013), lies outwith this fragment. Consequently, ecoPCR is unable to find any match between the reverse primer (jgHCO2198) and sequences in the invertebrate reference databases, causing *in silico* amplification failure. During *in vitro* tests, bands were observed by agarose gel electrophoresis for all invertebrate tissue tested (representing 38 species from 38 families within 10 major groups), and no bands were observed in PCR negative controls (Fig. S4).

#### 2.2 Taxonomic composition by sampling method

##### *2.2.1 Sweep-netting and morphotaxonomic identification*

Using the standard dataset, we identified 2,281 specimens belonging to 38 families across samples from 18 ponds. From this total, 1,404 specimens were identified as belonging to 91 species (see Tables S4 and S5 for lists). Overall, the most abundant taxa were *Asellus aquaticus* (11.68%), *Pisidium casertanum* (7.69%), *Erpobdella octoculata* (5.91%), *Coenagrion puella* (5.41%), and *Radix balthica* (5.27%) at species-level, and Chironomidae (33.14%), Asellidae (8.24%), Coenagrionidae (8.20%), Sphaeriidae (7.15%), and Dytiscidae (4.91%) at family-level. However, *Notonecta glauca* (*n* = 12 ponds), *C. puella* (*n* = 10), *Enallagma cyathigerum* (*n* = 10), *R. balthica* (*n* = 9), and *A. aquaticus* (*n* = 8) occurred in the most ponds at species-level, and Dytiscidae (*n* = 17), Chironomidae (*n* = 16), Coenagrionidae (*n* = 15), Corixidae (*n* = 13), and Notonectidae (*n* = 12) occurred in the most ponds at family-level.

Using the unbiased dataset, we identified 2,149 specimens belonging to 36 families across samples from 18 ponds. From this total, 1,272 specimens were identified as belonging to 72 species. The most abundant taxa were still *A. aquaticus* (12.89%), *P. casertanum* (8.49%), *E. octoculata* (6.53%), *C. puella* (5.97%), and *R. balthica* (5.82%) at species-level, and Chironomidae (35.18%), Asellidae (8.75%), Coenagrionidae (8.66%), Sphaeriidae (6.70%), and Dytiscidae (5.21%) at family-level. *N. glauca* (*n* = 12 ponds), *C. puella* (*n* = 10), *E. cyathigerum* (*n* = 10), *R. balthica* (*n* = 9), and *A. aquaticus* (*n* = 8) still occurred in the most ponds at species-level, and Dytiscidae (*n* = 17), Chironomidae (*n* = 16), Coenagrionidae (*n* = 15), Corixidae (*n* = 12), and Notonectidae (*n* = 12) still occurred in the most ponds at family-level.

*2.2.2 DNA metabarcoding*

The sequencing run generated 34,473,112 raw sequence reads. In total, 12,024,697 sequences remained after trimming, merging, chimera removal, and clustering (average read count of 32,324 per sample). From these sequences, 7,281,801 (60.56%) were assigned to a metazoan or non-metazoan taxonomic rank, but 4,742,896 were not assigned a taxonomic identity (39.44%). Assignments were then corrected (i.e. removed, renamed, or merged), the false positive sequence threshold applied, and coarse assignments and samples from other projects removed. Across the study ponds, 2,906,869 sequence reads were assigned to 55 families, of which 2,448,078 sequence reads were assigned to 131 species (see Tables S4 and S5 for lists).

Using the standard dataset, the majority of reads were assigned to *N. glauca* (14.85%), *A. aquaticus* (8.22%), *E. octoculata* (5.89%), *Chironomus luridus* (5.29%), and *Ilyocoris cimicoides* (4.09%) at species-level, and Notonectidae (14.60%), Chironomidae (13.06%), Corixidae (9.33%), Glossiphoniidae (7.50%), and Asellidae (7.26%) at family-level. The taxa that inhabited the most ponds were *C. puella* (*n* = 16 ponds), *N. glauca* (*n* = 15), *A. aquaticus* (*n* = 12), *C. luridus* (*n* = 12), and *Helobdella stagnalis* (*n* = 9) at species-level, and Chironomidae (*n* = 18), Naididae (*n* = 17), Coenagrionidae (*n* = 16), Dytiscidae (*n* = 16), Corixidae (*n* = 15), and Notonectidae (*n* = 15) at family-level.

Using the unbiased dataset 2,900,282 sequence reads were assigned to 46 families, of which , 2,447,317 sequence reads were assigned to 118 species. The majority of reads were assigned to *N. glauca* (14.86%), *A. aquaticus* (8.23%), *E. octoculata* (5.89%), *C. luridus* (5.29%), and *I. cimicoides* (4.10%) at species-level, and Notonectidae (14.64%), Chironomidae (13.09%), Corixidae (9.35%), Glossiphoniidae (7.51%), and Asellidae (7.27%) at family-level. The taxa that inhabited the most ponds were *C. puella* (*n* = 16 ponds), *N. glauca* (*n* = 15), *A. aquaticus* (*n* = 12), *C. luridus* (*n* = 12), and *H. stagnalis* (*n* = 9) at species-level, and Chironomidae (*n* = 18), Coenagrionidae (*n* = 16), Dytiscidae (*n* = 16), Corixidae (*n* = 15), and Notonectidae (*n* = 15) at family-level.

*2.2.3 eDNA metabarcoding*

The sequencing run generated 11,019,530 raw sequence reads. In total, 4,267,530 sequences remained after trimming, merging, chimera removal, and clustering (average read count of 16,075 per sample). From these sequences, 1,726,801 (40.46%) were assigned to a metazoan or non-metazoan taxonomic rank, but 2,540,729 were not assigned a taxonomic identity (59.54%). Assignments were then corrected (i.e. removed, renamed, or merged), the false positive sequence threshold applied, and coarse assignments and samples from other projects removed. Across the study ponds, 813,376 sequence reads were assigned to 90 families, of which 346,163 sequence reads were assigned to 145 species (see Tables S4 and S5 for lists).

Using the standard dataset, the majority of reads were assigned to *Cyclops strenuus* (13.75%), *Cloeon dipterum* (12.93%), *Keratella cochlearis* (11.68%), *Cypridopsis vidua* (7.56%), and *Pristina longiseta* (7.49%) at species-level, and Cyclopidae (42.43%), Chironomidae (15.26%), Brachionidae (6.82%), Naididae (6.72%), and Baetidae (5.50%) at family-level. However, the most common taxa across the study ponds were *Rotaria rotatoria* ($n$ = 16 ponds), *Chaetogaster diastrophus* ($n$ = 14), *C. dipterum* ($n$ = 14), *Eucyclops serrulatus* ($n$ = 14), and *Simocephalus vetulus* ($n$ = 13) at species-level, and Chironomidae ($n$ = 18), Cyclopidae ($n$ = 18), Macrotrichidae ($n$ = 17), Philodinidae ($n$ = 17), and Chaetonotidae ($n$ = 16) at family-level.

Using the unbiased dataset, 586,416 sequence reads were assigned to 61 families, of which 215,312 sequence reads were assigned to 93 species. The majority of reads were assigned to *C. strenuus* (22.10%), *C. dipterum* (20.80%), *P. longiseta* (12.04%), *Libellula quadrimaculata* (11.76%), and *C. diastrophus* (6.54%) at species-level, and Cyclopidae (51.68%), Chironomidae (21.17%), Naididae (7.92%), Baetidae (7.64%) and Libellulidae (4.33%) at family-level. However, the most common taxa across the study ponds were *C. diastrophus* ($n$ = 14), C. *dipterum* ($n$ = 14 ponds), *S. vetulus* ($n$ = 13), *Macrocyclops albidus* ($n$ = 11), and *C. strenuus* ($n$ = 9) at species-level, and Chironomidae ($n$ = 18), Cyclopidae ($n$ = 18), Naididae ($n$ = 16), Baetidae ($n$ = 14), and Daphniidae ($n$ = 14) at family-level.

*2.2.4 Combined methods*

The three standard methods of invertebrate assessment combined identified 249 species and 110 families across the study ponds. The combined data including taxa without reference sequences that were detected by microscopy and meiofauna detected by metabarcoding indicated that *C. puella* ($n$ = 16 ponds), *R. rotatoria* ($n$ = 16), *C. dipterum* ($n$ = 15), *N. glauca* ($n$ = 15), and *C. diastrophus* ($n$ = 15) were the most common species, and Chironomidae ($n$ = 18), Dytiscidae ($n$ = 18), Cyclopidae ($n$ = 18), Naididae ($n$ = 18), and Coenagrionidae ($n$ = 17) were the most common families.

The three unbiased methods of invertebrate assessment combined identified 176 species and 77 families across the study ponds. The combined data excluding taxa without reference sequences that were detected by microscopy and meiofauna detected by metabarcoding indicated that *C. puella* ($n$ = 16), *C. dipterum* ($n$ = 15), *N. glauca* ($n$ = 15), *C. diastrophus* ($n$ = 15), and *C. luridus* ($n$ = 14) were the most common species, and Chironomidae ($n$ = 18), Dytiscidae ($n$ = 18), Cyclopidae ($n$ = 18), Naididae ($n$ = 18), and Coenagrionidae ($n$ = 17) were the most common families.

#### 2.3 Impact of crucian carp stocking on pond macroinvertebrates

Independently and combined, methods revealed alpha diversity of macroinvertebrates was marginally reduced in ponds containing crucian carp at species (Figs. S13ai-iv) and family-level (Figs. S13bi-iv), but not significantly so ($P$ > 0.05, Table S11). Detailed examination of alpha diversity within the major macroinvertebrate groups (Dobson, Pawley, Fletcher, & Powell, 2012) identified by all three methods combined revealed that Coleoptera and Mollusca diversity was significantly reduced (albeit Coleoptera was borderline) in ponds with crucian carp at species-level (GLM $\chi^2_{18}$ = 23.407, $P$ = 0.175; Coleoptera -0.481 ± 0.249, $Z$

= -1.930, *P* = 0.054; Mollusca -0.821 ± 0.283, *Z* = -2.904, *P* = 0.004), but not family-level (GLM $\chi^2_{18}$ = 14.681, *P* = 0.684; Coleoptera -0.544 ± 0.296, *Z* = -1.834, *P* = 0.067; Mollusca -0.499 ± 0.308, *Z* = -1.623, *P* = 0.105). However, differences in alpha diversity between ponds with or without crucian carp were not significant for other invertebrate groups at either taxonomic rank (Fig. S14).

Total beta diversity of ponds was consistently high at species and family-level for independent and combined methods. Variation in macroinvertebrate community composition was predominantly driven by turnover rather than nestedness-resultant (Table S12). Using morphotaxonomic identification and DNA metabarcoding, MVDISP did not significantly differ between ponds for any component of beta diversity at species or family-level. Using eDNA metabarcoding, MVDISP significantly differed between ponds for species turnover and total beta diversity, but not nestedness-resultant. No significant difference in MVDISP was observed at family-level for any beta diversity component. All methods combined indicated MVDISP significantly differed between ponds for turnover at species-level and total beta diversity at species and family-level, but not any other beta diversity component at either taxonomic rank (Table S13).

At species-level, all methods combined revealed a weak or moderate positive influence of crucian carp presence on turnover (Table S14; Fig. S15aiv) and total beta diversity (Table S14; Fig. S15civ) between ponds, but not nestedness-resultant (Table S14; Fig. S15biv). eDNA metabarcoding also revealed a weak positive influence of crucian carp presence on total beta diversity (Table S14; Fig. S15ciii). In contrast, morphotaxonomic identification and DNA metabarcoding did not identify a significant effect of crucian carp presence on turnover, nestedness-resultant, or total beta diversity (Table S14; Figs. S15ai, aii, bi, bii, ci, cii). At family-level, crucian carp did not influence turnover, nestedness-resultant, or total beta diversity according to methods independently or combined (Table S14; Fig. S16). Overall, eDNA metabarcoding and all methods combined produced concurrent results at species-level, but all methods independently and combined were in agreement at family-level.

Additional analyses undertaken on the combined method dataset for macroinvertebrates supported an effect of crucian carp presence-absence and excluded the influence of abiotic variables on pond invertebrate diversity. Forward selection did not identify any significant abiotic variables for any beta diversity component at either taxonomic rank. Consequently, variance partitioning analysis was not undertaken for any beta diversity component. At species-level, RDA of each beta diversity component minus abiotic data revealed crucian carp presence-absence influenced turnover ($F_1$ = 1.707, *P* = 0.015) and total beta diversity ($F_1$ = 1.559, *P* = 0.014), but not nestedness-resultant ($F_1$ = 0.495, *P* = 0.769). At family-level, RDA of each beta diversity component minus abiotic data revealed crucian carp presence-absence did not influence turnover ($F_1$ = 1.206, *P* = 0.247), nestedness-resultant ($F_1$ = 2.756, *P* = 0.060) or total beta diversity ($F_1$ = 1.376, *P* = 0.105).

### Appendix 3: Tables

**Table S1.** Available fyke net survey data for study ponds in Norfolk, eastern England. Rows containing data for the last year of fyke net survey prior to this study are highlighted in yellow. Number of crucian carp individuals and other fish caught during fyke net surveys are highlighted in blue. Provided as an Excel spreadsheet.

**Table S2.** Results of size-sorting for invertebrate specimens collected from each pond. Provided as an Excel spreadsheet.

**Table S3.** Tables containing: **(a)** list of UK invertebrate species that amplified during *in silico* PCR using ecoPCR software; **(b)** list of UK invertebrate species that did not amplify during *in silico* PCR using ecoPCR software; **(c)** List of UK invertebrate species that did not have any *COI* reference sequences on the NCBI nucleotide database at the time of batch sequence download (April 2017); and **(d)** summary of *in silico* amplification and reference database representation for UK invertebrate species living in or associated with freshwater habitats that are expected to occur in the UK. Provided as an Excel spreadsheet.

**Table S4.** Number of ponds where each invertebrate species was detected by sweep-netting and microscopy, DNA metabarcoding, and eDNA metabarcoding respectively. Provided as an Excel spreadsheet.

**Table S5.** Number of ponds where each invertebrate family was detected by sweep-netting and microscopy, DNA metabarcoding, and eDNA metabarcoding respectively. Provided as an Excel spreadsheet.

**Table S6.** Number of species in each invertebrate group that were detected by the different standard methods.

| Group | Sweep-netting and microscopy | DNA metabarcoding | eDNA metabarcoding |
|---|---|---|---|
| Annelida | 0 | 17 | 17 |
| Arachnida | 0 | 1 | 4 |
| Coleoptera | 21 | 26 | 4 |
| Collembola | 0 | 1 | 1 |
| Crustacea | 4 | 6 | 24 |
| Diptera | 0 | 23 | 27 |
| Ephemeroptera | 1 | 2 | 2 |
| Gastrotricha | 0 | 0 | 3 |
| Hemiptera | 19 | 10 | 7 |
| Hirudinea | 7 | 6 | 2 |
| Hymenoptera | 0 | 0 | 1 |
| Lepidoptera | 0 | 0 | 1 |
| Megaloptera | 1 | 1 | 0 |
| Mollusca | 24 | 17 | 13 |
| Nematoda | 0 | 0 | 1 |
| Odonata | 11 | 10 | 4 |
| Platyhelminthes | 0 | 0 | 1 |
| Psocoptera | 0 | 0 | 1 |
| Rotifera | 0 | 7 | 27 |
| Tardigrada | 0 | 1 | 1 |
| Trichoptera | 3 | 3 | 4 |
| **Total** | 91 | 131 | 145 |

**Table S7.** Number of families in each invertebrate group that were detected by the different standard methods.

| **Group** | **Sweep-netting and microscopy** | **DNA metabarcoding** | **eDNA metabarcoding** |
|---|---|---|---|
| Annelida | 0 | 3 | 5 |
| Arachnida | 0 | 2 | 8 |
| Coleoptera | 7 | 7 | 5 |
| Collembola | 0 | 1 | 4 |
| Crustacea | 3 | 8 | 12 |
| Diptera | 6 | 7 | 13 |
| Ephemeroptera | 1 | 1 | 2 |
| Gastrotricha | 0 | 1 | 2 |
| Hemiptera | 6 | 4 | 5 |
| Hirudinea | 2 | 2 | 1 |
| Hymenoptera | 0 | 0 | 3 |
| Lepidoptera | 0 | 0 | 1 |
| Megaloptera | 1 | 1 | 0 |
| Mollusca | 7 | 8 | 8 |
| Nematoda | 0 | 0 | 1 |
| Odonata | 3 | 3 | 2 |
| Platyhelminthes | 0 | 2 | 2 |
| Psocoptera | 0 | 0 | 1 |
| Rotifera | 0 | 2 | 10 |
| Tardigrada | 0 | 1 | 2 |
| Thysanoptera | 0 | 0 | 1 |
| Trichoptera | 2 | 2 | 2 |
| **Total** | 38 | 55 | 90 |

**Table S8.** Summary of analyses (ANOVA) statistically comparing homogeneity of multivariate dispersions (MVDISP) between the invertebrate communities produced by each standard and unbiased sampling method at species-level and family-level.

| | Homogeneity of multivariate dispersions (ANOVA) | | | | | | | |
|---|---|---|---|---|---|---|---|---|
| | **Species-level** | | | | **Family-level** | | | |
| | **Mean distance to centroid ± SE** | **df** | ***F*** | ***P*** | **Mean distance to centroid ± SE** | **df** | ***F*** | ***P*** |
| **Standard methods** | | | | | | | | |
| *Turnover* | | 2 | 2.467 | 0.095 | | 2 | 0.271 | 0.764 |
| Microscopy | 0.559 ± 0.005 | | | | 0.379 ± 0.010 | | | |
| DNA metabarcoding | 0.552 ± 0.004 | | | | 0.385 ± 0.011 | | | |
| eDNA metabarcoding | 0.507 ± 0.007 | | | | 0.360 ± 0.014 | | | |
| *Nestedness-resultant* | | 2 | 0.752 | 0.477 | | 2 | 0.707 | 0.498 |
| Microscopy | 0.042 ± 0.001 | | | | 0.101 ± 0.007 | | | |
| DNA metabarcoding | 0.039 ± 0.002 | | | | 0.067 ± 0.005 | | | |
| eDNA metabarcoding | 0.055 ± 0.003 | | | | 0.089 ± 0.011 | | | |
| *Total beta diversity* | | 2 | 1.817 | 0.173 | | 2 | 0.214 | 0.828 |
| Microscopy | 0.595 ± 0.003 | | | | 0.466 ± 0.007 | | | |
| DNA metabarcoding | 0.585 ± 0.001 | | | | 0.458 ± 0.001 | | | |
| eDNA metabarcoding | 0.559 ± 0.006 | | | | 0.452 ± 0.004 | | | |
| **Unbiased methods** | | | | | | | | |
| *Turnover* | | 2 | 0.022 | 0.978 | | 2 | 0.327 | 0.723 |
| Microscopy | 0.543 ± 0.006 | | | | 0.374 ± 0.010 | | | |
| DNA metabarcoding | 0.543 ± 0.005 | | | | 0.380 ± 0.009 | | | |
| eDNA metabarcoding | 0.538 ± 0.009 | | | | 0.402 ± 0.016 | | | |
| *Nestedness-resultant* | | 2 | 0.184 | 0.832 | | 2 | 0.504 | 0.607 |
| Microscopy | 0.052 ± 0.002 | | | | 0.102 ± 0.006 | | | |
| DNA metabarcoding | 0.045 ± 0.002 | | | | 0.070 ± 0.006 | | | |
| eDNA metabarcoding | 0.054 ± 0.002 | | | | 0.101 ± 0.023 | | | |
| *Total beta diversity* | | 2 | 0.035 | 0.966 | | 2 | 1.702 | 0.193 |
| Microscopy | 0.585 ± 0.004 | | | | 0.464 ± 0.006 | | | |
| DNA metabarcoding | 0.580 ± 0.002 | | | | 0.452 ± 0.002 | | | |
| eDNA metabarcoding | 0.581 ± 0.007 | | | | 0.492 ± 0.005 | | | |

**Table S9.** Summary of analyses (PERMANOVA) statistically examining variation in species-level and family-level invertebrate community composition across standard and unbiased methods.

| | Community similarity (PERMANOVA) | | | | | | | |
|---|---|---|---|---|---|---|---|---|
| | Species-level | | | | Family-level | | | |
| | **df** | ***F*** | $R^2$ | ***P*** | **df** | ***F*** | $R^2$ | ***P*** |
| **Standard methods** | | | | | | | | |
| Turnover | 2 | 6.619 | 0.206 | **0.001** | 2 | 15.686 | 0.381 | **0.001** |
| Nestedness-resultant | 2 | -10.146 | -0.661 | 1.000 | 2 | -8.242 | -0.478 | 1.000 |
| Total beta diversity | 2 | 4.922 | 0.162 | **0.001** | 2 | 10.563 | 0.293 | **0.001** |
| **Unbiased methods** | | | | | | | | |
| Turnover | 2 | 5.566 | 0.179 | **0.001** | 2 | 15.686 | 0.381 | **0.001** |
| Nestedness-resultant | 2 | -5.673 | -0.286 | 0.999 | 2 | -8.201 | -0.474 | 1.000 |
| Total beta diversity | 2 | 4.234 | 0.142 | **0.001** | 2 | 10.563 | 0.293 | **0.001** |

**Table S10.** Summary of analyses (GLM) statistically comparing macroinvertebrate alpha diversity at species-level and family-level between ponds with and without crucian carp using independent and combined unbiased methods.

| | **Species-level** | | | | | | **Family-level** | | | | | |
|---|---|---|---|---|---|---|---|---|---|---|---|---|
| | **GLM** | | | | **LRT** | | **GLM** | | | | **LRT** | |
| | **df** | **Estimate ± SE** | ***Z*** | ***P*** | **$\chi^2$** | ***P*** | **df** | **Estimate ± SE** | ***Z*** | ***P*** | **$\chi^2$** | ***P*** |
| Sweep-netting and microscopy | 1 | -0.174 ± 0.185 | -0.939 | 0.348 | 0.882 | 0.348 | 1 | -0.123 ± 0.162 | -0.756 | 0.450 | 0.572 | 0.449 |
| DNA metabarcoding | 1 | 0.058 ± 0.147 | 0.391 | 0.696 | 0.153 | 0.696 | 1 | 0.016 ± 0.127 | 0.127 | 0.899 | 0.016 | 0.899 |
| eDNA metabarcoding | 1 | -0.250 ± 0.203 | -1.230 | 0.219 | 1.515 | 0.218 | 1 | -0.243 ± 0.155 | -1.561 | 0.119 | 2.446 | 0.118 |
| Combined methods | 1 | -0.139 ± 0.114 | -1.216 | 0.224 | 1.480 | 0.224 | 1 | -0.155 ± 0.099 | -1.571 | 0.116 | 2.476 | 0.116 |

**Table S11.** Relative contribution of taxon turnover and nestedness to total macroinvertebrate beta diversity (Jaccard dissimilarity) produced by independent and combined unbiased methods. A value of 1 corresponds to all sites containing different species.

| | **Species-level** | | | **Family-level** | | |
|---|---|---|---|---|---|---|
| | **Turnover** | **Nestedness-resultant** | **Total beta diversity** | **Turnover** | **Nestedness-resultant** | **Total beta diversity** |
| Sweep-netting and microscopy | 0.927 (97.68%) | 0.022 (2.32%) | 0.949 (100%) | 0.863 (94.42%) | 0.051 (5.58%) | 0.914 (100%) |
| DNA metabarcoding | 0.934 (98.32%) | 0.016 (1.68%) | 0.950 (100%) | 0.878 (96.38%) | 0.033 (3.62%) | 0.911 (100%) |
| eDNA metabarcoding | 0.928 (97.58%) | 0.023 (2.42%) | 0.951 (100%) | 0.890 (96.01%) | 0.037 (3.99%) | 0.927 (100%) |
| Combined methods | 0.927 (98.41%) | 0.015 (1.59%) | 0.942 (100%) | 0.875 (96.58%) | 0.031 (3.42%) | 0.906 (100%) |

**Table S12.** Summary of analyses (ANOVA) statistically comparing homogeneity of multivariate dispersions (MVDISP) between the species-level and family-level macroinvertebrate communities in ponds with and without crucian carp according to the independent and combined unbiased methods.

| | Homogeneity of multivariate dispersions (ANOVA) | | | | | | | |
|---|---|---|---|---|---|---|---|---|
| | Species-level | | | | Family-level | | | |
| | **Mean distance to centroid ± SE** | **df** | ***F*** | ***P*** | **Mean distance to centroid ± SE** | **df** | ***F*** | ***P*** |
| **Sweep-netting and microscopy** | | | | | | | | |
| *Turnover* | | 1 | 2.303 | 0.149 | | 1 | 1.144 | 0.301 |
| Crucian carp | 0.544 ± 0.003 | | | | 0.387 ± 0.008 | | | |
| No crucian carp | 0.491 ± 0.008 | | | | 0.333 ± 0.015 | | | |
| *Nestedness-resultant* | | 1 | 1.114 | 0.307 | | 1 | 0.881 | 0.362 |
| Crucian carp | 0.042 ± 0.001 | | | | 0.077 ± 0.003 | | | |
| No crucian carp | 0.063 ± 0.002 | | | | 0.115 ± 0.012 | | | |
| *Total beta diversity* | | 1 | 1.179 | 0.294 | | 1 | 0.555 | 0.467 |
| Crucian carp | 0.577 ± 0.001 | | | | 0.464 ± 0.006 | | | |
| No crucian carp | 0.545 ± 0.007 | | | | 0.432 ± 0.011 | | | |
| **DNA metabarcoding** | | | | | | | | |
| *Turnover* | | 1 | 0.033 | 0.858 | | 1 | 0.003 | 0.959 |
| Crucian carp | 0.523 ± 0.006 | | | | 0.351 ± 0.015 | | | |
| No crucian carp | 0.517 ± 0.005 | | | | 0.354 ± 0.011 | | | |
| *Nestedness-resultant* | | 1 | 0.948 | 0.345 | | 1 | 0.055 | 0.817 |
| Crucian carp | 0.056 ± 0.003 | | | | 0.062 ± 0.006 | | | |
| No crucian carp | 0.035 ± 0.001 | | | | 0.071 ± 0.007 | | | |
| *Total beta diversity* | | 1 | 0.452 | 0.511 | | 1 | 0.194 | 0.666 |
| Crucian carp | 0.567 ± 0.003 | | | | 0.424 ± 0.004 | | | |
| No crucian carp | 0.552 ± 0.002 | | | | 0.437 ± 0.004 | | | |
| **eDNA metabarcoding** | | | | | | | | |
| *Turnover* | | 1 | 27.253 | **<0.001** | | 1 | 0.002 | 0.966 |
| Crucian carp | 0.589 ± 0.004 | | | | 0.383 ± 0.030 | | | |
| No crucian carp | 0.443 ± 0.003 | | | | 0.380 ± 0.010 | | | |
| *Nestedness-resultant* | | 1 | 0.162 | 0.693 | | 1 | 1.201 | 0.289 |
| Crucian carp | 0.045 ± 0.001 | | | | 0.145 ± 0.035 | | | |
| No crucian carp | 0.054 ± 0.003 | | | | 0.072 ± 0.006 | | | |
| *Total beta diversity* | | 1 | 21.273 | **<0.001** | | 1 | 1.233 | 0.283 |
| Crucian carp | 0.617 ± 0.003 | | | | 0.493 ± 0.008 | | | |
| No crucian carp | 0.497 ± 0.003 | | | | 0.455 ± 0.003 | | | |
| **Combined methods** | | | | | | | | |

| | Homogeneity of multivariate dispersions (ANOVA) | | | | | | | |
|---|---|---|---|---|---|---|---|---|
| | Species-level | | | | Family-level | | | |
| | **Mean distance to centroid ± SE** | **df** | ***F*** | ***P*** | **Mean distance to centroid ± SE** | **df** | ***F*** | ***P*** |
| *Turnover* | | 1 | 6.385 | **0.022** | | 1 | 1.062 | 0.318 |
| Crucian carp | 0.527 ± 0.002 | | | | 0.376 ± 0.005 | | | |
| No crucian carp | 0.465 ± 0.004 | | | | 0.343 ± 0.005 | | | |
| *Nestedness-resultant* | | 1 | 0.032 | 0.860 | | 1 | 0.135 | 0.718 |
| Crucian carp | 0.033 ± 0.001 | | | | 0.056 ± 0.003 | | | |
| No crucian carp | 0.035 ± 0.001 | | | | 0.047 ± 0.002 | | | |
| *Total beta diversity* | | 1 | 9.344 | **0.008** | | 1 | 4.891 | **0.042** |
| Crucian carp | 0.554 ± 0.001 | | | | 0.437 ± 0.001 | | | |
| No crucian carp | 0.503 ± 0.002 | | | | 0.393 ± 0.002 | | | |

**Table S13.** Summary of analyses (PERMANOVA) statistically examining variation in community composition between ponds with and without crucian carp at species-level and family-level using independent and combined unbiased methods.

| | Community similarity (PERMANOVA) | | | | | | | |
|---|---|---|---|---|---|---|---|---|
| | Species-level | | | | Family-level | | | |
| | **df** | ***F*** | $R^2$ | ***P*** | **df** | ***F*** | $R^2$ | ***P*** |
| **Netting and microscopy** | | | | | | | | |
| Turnover | 1 | 1.628 | 0.092 | 0.055 | 1 | 0.987 | 0.058 | 0.477 |
| Nestedness-resultant | 1 | -2.048 | -0.147 | 0.857 | 1 | 0.357 | 0.022 | 0.561 |
| Total beta diversity | 1 | 1.408 | 0.081 | 0.056 | 1 | 1.108 | 0.065 | 0.341 |
| **DNA metabarcoding** | | | | | | | | |
| Turnover | 1 | 1.372 | 0.079 | 0.112 | 1 | 2.039 | 0.113 | 0.052 |
| Nestedness-resultant | 1 | -0.175 | -0.011 | 0.615 | 1 | -0.113 | -0.007 | 0.728 |
| Total beta diversity | 1 | 1.197 | 0.070 | 0.176 | 1 | 1.520 | 0.087 | 0.085 |
| **eDNA metabarcoding** | | | | | | | | |
| Turnover | 1 | 1.335 | 0.077 | 0.160 | 1 | 1.348 | 0.078 | 0.239 |
| Nestedness-resultant | 1 | 2.388 | 0.130 | 0.358 | 1 | 2.238 | 0.123 | 0.313 |
| Total beta diversity | 1 | 1.341 | 0.077 | **0.047** | 1 | 1.245 | 0.072 | 0.155 |
| **Combined methods** | | | | | | | | |
| Turnover | 1 | 1.724 | 0.097 | **0.010** | 1 | 1.191 | 0.069 | 0.315 |
| Nestedness-resultant | 1 | -1.061 | -0.071 | 0.915 | 1 | 5.065 | 0.240 | 0.079 |
| Total beta diversity | 1 | 1.559 | 0.089 | **0.012** | 1 | 1.379 | 0.079 | 0.087 |

**Table S14.** Summary of analyses (ANOVA) statistically comparing homogeneity of multivariate dispersions (MVDISP) between the species-level and family-level invertebrate communities in ponds with and without crucian carp according to the independent and combined standard methods.

| | Homogeneity of multivariate dispersions (ANOVA) | | | | | | | |
|---|---|---|---|---|---|---|---|---|
| | Species-level | | | | Family-level | | | |
| | **Mean distance to centroid ± SE** | **df** | ***F*** | ***P*** | **Mean distance to centroid ± SE** | **df** | ***F*** | ***P*** |
| **Sweep-netting and microscopy** | | | | | | | | |
| *Turnover* | | 1 | 3.706 | 0.072 | | 1 | 1.522 | 0.235 |
| Crucian carp | 0.562 ± 0.002 | | | | 0.393 ± 0.009 | | | |
| No crucian carp | 0.502 ± 0.007 | | | | 0.331 ± 0.014 | | | |
| *Nestedness-resultant* | | 1 | 2.210 | 0.157 | | 1 | 0.748 | 0.400 |
| Crucian carp | 0.032 ± 0.001 | | | | 0.080 ± 0.003 | | | |
| No crucian carp | 0.060 ± 0.003 | | | | 0.115 ± 0.012 | | | |
| *Total beta diversity* | | 1 | 1.576 | 0.227 | | 1 | 0.744 | 0.401 |
| Crucian carp | 0.588 ± 0.001 | | | | 0.468 ± 0.007 | | | |
| No crucian carp | 0.554 ± 0.006 | | | | 0.428 ± 0.013 | | | |
| **DNA metabarcoding** | | | | | | | | |
| *Turnover* | | 1 | 0.024 | 0.879 | | 1 | 0.148 | 0.706 |
| Crucian carp | 0.532 ± 0.006 | | | | 0.354 ± 0.021 | | | |
| No crucian carp | 0.527 ± 0.004 | | | | 0.375 ± 0.005 | | | |
| *Nestedness-resultant* | | 1 | 0.858 | 0.368 | | 1 | 0.049 | 0.827 |
| Crucian carp | 0.047 ± 0.002 | | | | 0.071 ± 0.007 | | | |
| No crucian carp | 0.030 ± 0.001 | | | | 0.063 ± 0.005 | | | |
| *Total beta diversity* | | 1 | 0.441 | 0.516 | | 1 | 0.133 | 0.720 |
| Crucian carp | 0.572 ± 0.003 | | | | 0.436 ± 0.002 | | | |
| No crucian carp | 0.558 ± 0.001 | | | | 0.444 ± 0.002 | | | |
| **eDNA metabarcoding** | | | | | | | | |
| *Turnover* | | 1 | 2.634 | 0.124 | | 1 | 0.027 | 0.871 |
| Crucian carp | 0.505 ± 0.014 | | | | 0.344 ± 0.026 | | | |
| No crucian carp | 0.435 ± 0.003 | | | | 0.334 ± 0.007 | | | |
| *Nestedness-resultant* | | 1 | 0.280 | 0.604 | | 1 | 0.868 | 0.365 |
| Crucian carp | 0.070 ± 0.005 | | | | 0.111 ± 0.017 | | | |
| No crucian carp | 0.054 ± 0.003 | | | | 0.064 ± 0.005 | | | |
| *Total beta diversity* | | 1 | 5.580 | **0.031** | | 1 | 3.829 | 0.068 |
| Crucian carp | 0.570 ± 0.007 | | | | 0.458 ± 0.005 | | | |
| No crucian carp | 0.488 ± 0.004 | | | | 0.405 ± 0.002 | | | |

| | Homogeneity of multivariate dispersions (ANOVA) | | | | | | | |
|---|---|---|---|---|---|---|---|---|
| | Species-level | | | | Family-level | | | |
| | **Mean distance to centroid ± SE** | **df** | ***F*** | ***P*** | **Mean distance to centroid ± SE** | **df** | ***F*** | ***P*** |
| **Combined methods** | | | | | | | | |
| *Turnover* | | 1 | 4.052 | 0.061 | | 1 | 0.956 | 0.343 |
| Crucian carp | 0.518 ± 0.002 | | | | 0.365 ± 0.009 | | | |
| No crucian carp | 0.463 ± 0.005 | | | | 0.328 ± 0.004 | | | |
| *Nestedness-resultant* | | 1 | 0.008 | 0.930 | | 1 | 0.356 | 0.559 |
| Crucian carp | 0.032 ± 0.001 | | | | 0.062 ± 0.004 | | | |
| No crucian carp | 0.034 ± 0.002 | | | | 0.046 ± 0.002 | | | |
| *Total beta diversity* | | 1 | 6.987 | **0.018** | | 1 | 8.085 | **0.012** |
| Crucian carp | 0.549 ± 0.002 | | | | 0.435 ± 0.002 | | | |
| No crucian carp | 0.502 ± 0.001 | | | | 0.378 ± 0.002 | | | |

#### Appendix 4: Figures

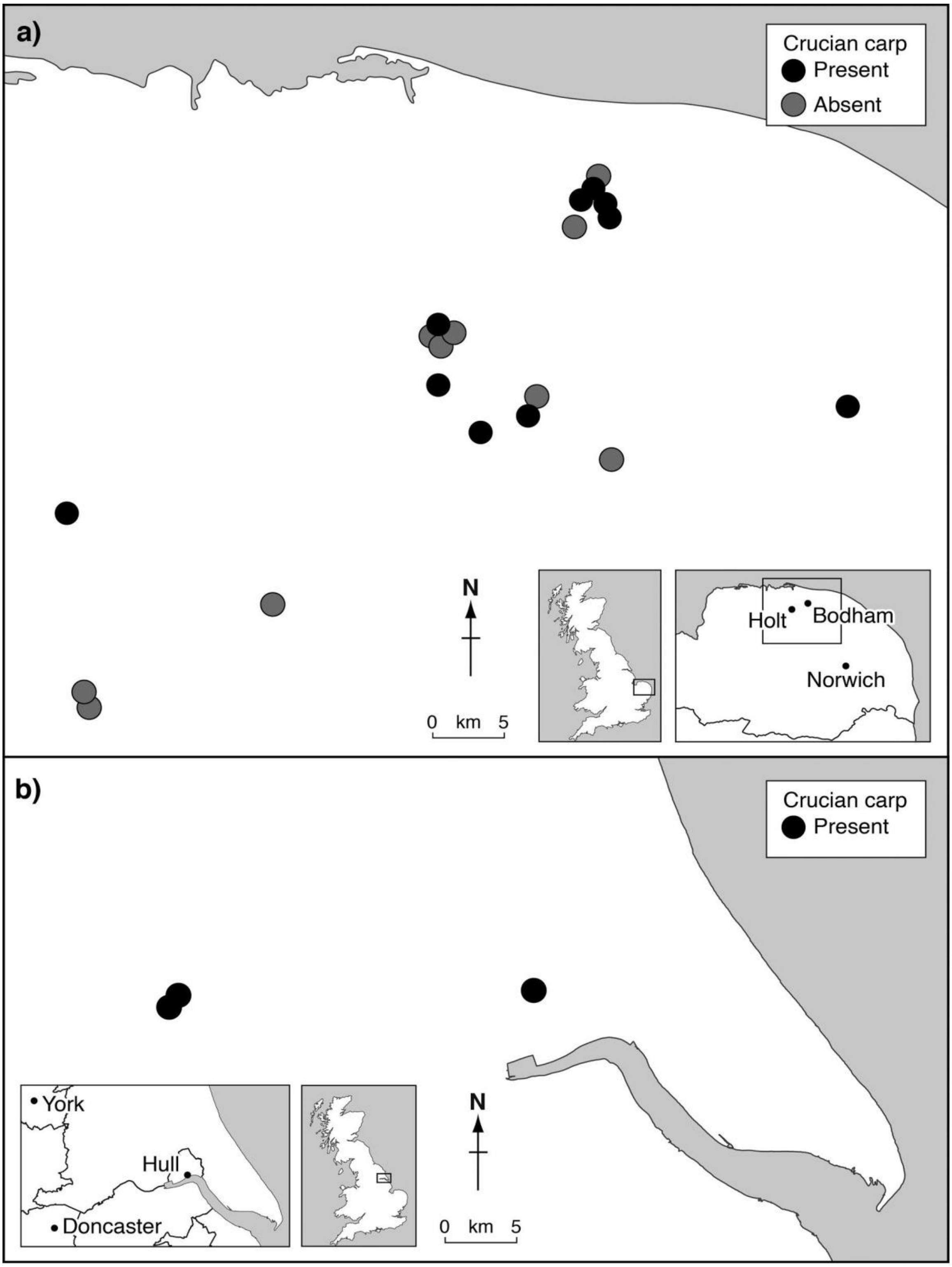


**Figure S1.** Map showing the location of study ponds across Norfolk (*n* = 10 with crucian, *n* = 10 without crucian carp) and East Yorkshire (*n* = 3 with crucian carp). Five of the Norfolk ponds (*n* = 3 with crucian carp, *n* = 2 without crucian carp) were not included for metabarcoding.

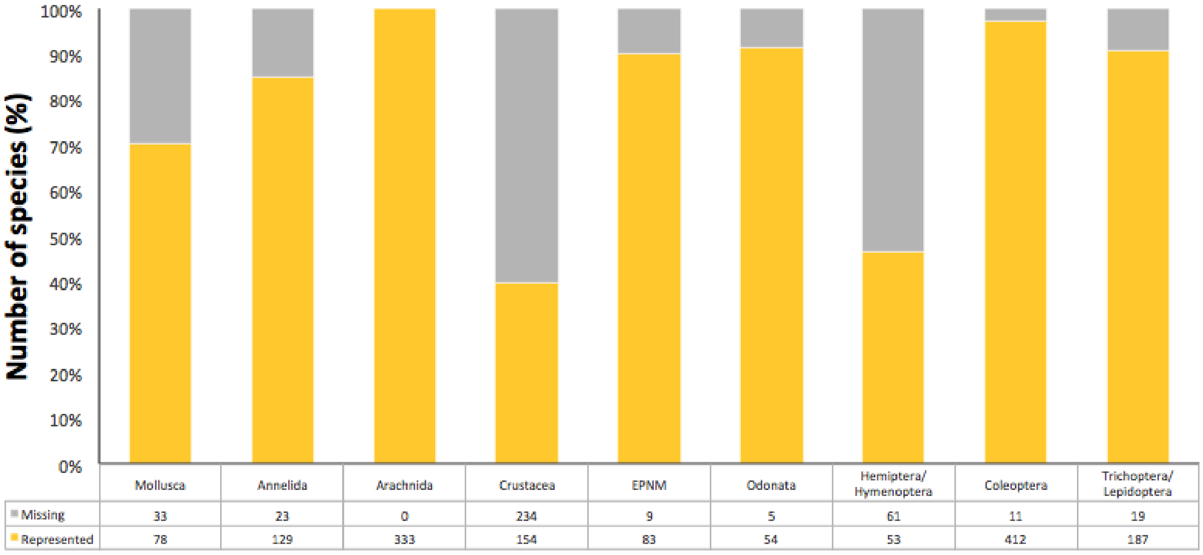


**Figure S2.** Barplot summarising the number and proportion of species with and without reference sequences from GenBank for each custom invertebrate database. The extent of reference sequence representation for species varied across the invertebrate databases: Coleoptera 97.40% (*N* = 423), Odonata 91.53% (*N* = 59), Hemiptera and Hymenoptera 46.49% (*N* = 114), Trichoptera and Lepidoptera 90.78% (*N* = 206), Ephemeroptera, Plecoptera, Neuroptera, and Megaloptera 90.22% (*N* = 92), Crustacea 39.69% (*N* = 388), Mollusca 70.27% (*N* = 111), Arachnida 100% (*N* = 333), and Annelida 84.87% (*N* = 152).

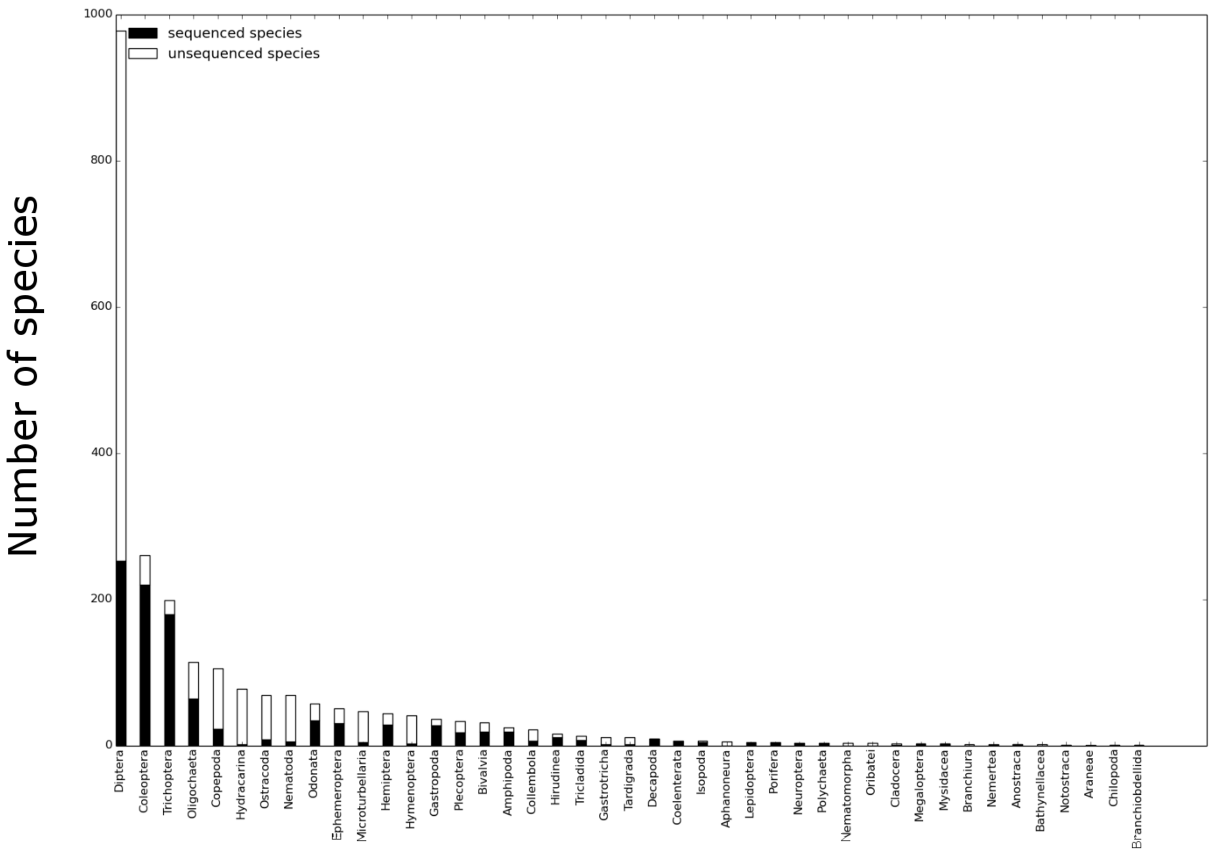


**Figure S3.** Barplot summarising the number of species with and without records on GenBank according to invertebrate groups.

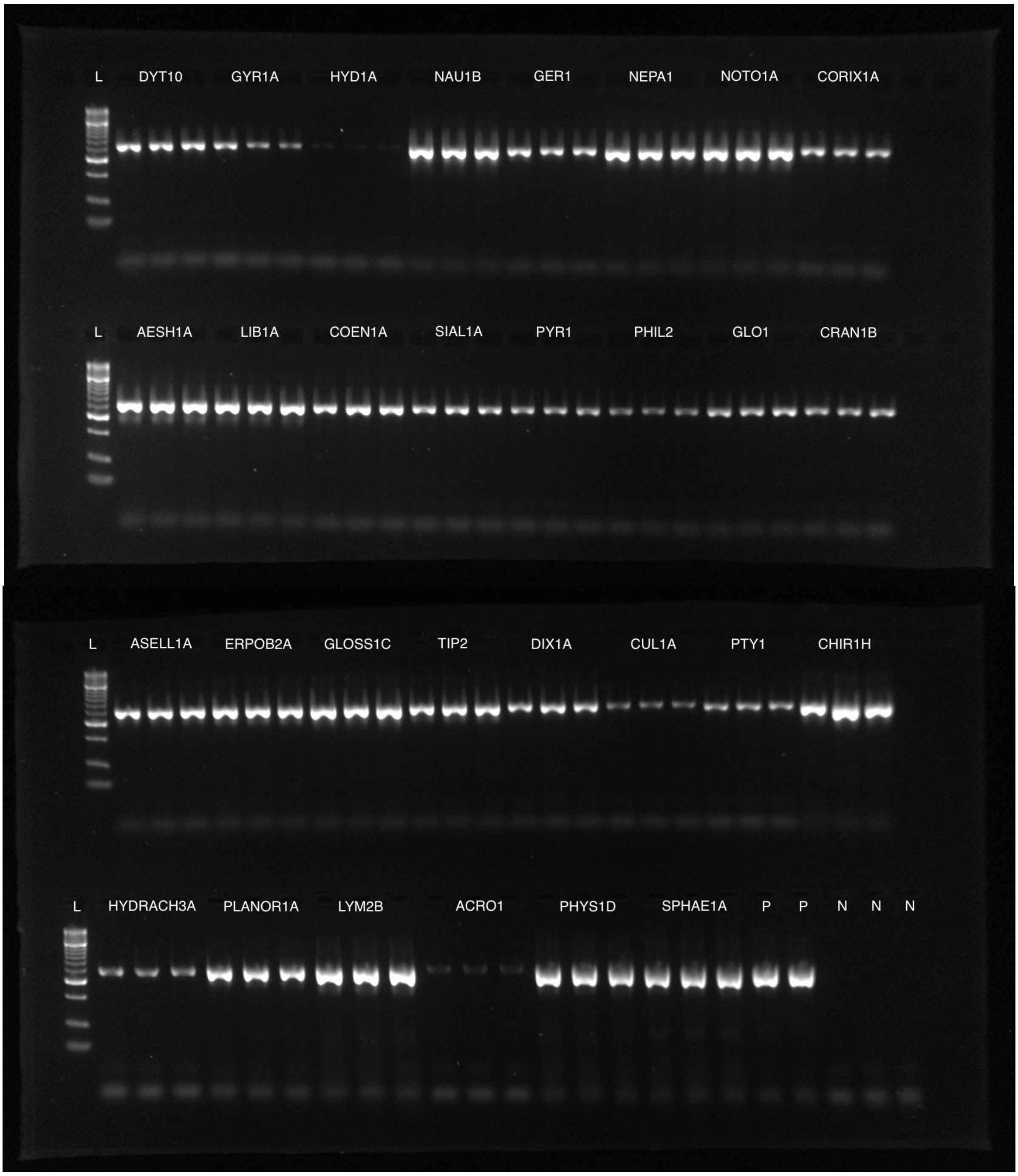
L DYT10 GYR1A HYD1A NAU1B GER1 NEPA1 NOTO1A CORIX1A
L AESH1A LIB1A COEN1A SIAL1A PYR1 PHIL2 GLO1 CRAN1B
L ASELL1A ERPOB2A GLOSS1C TIP2 DIX1A CUL1A PTY1 CHIR1H
L HYDRACH3A PLANOR1A LYM2B ACRO1 PHYS1D SPHAE1A P P N N N

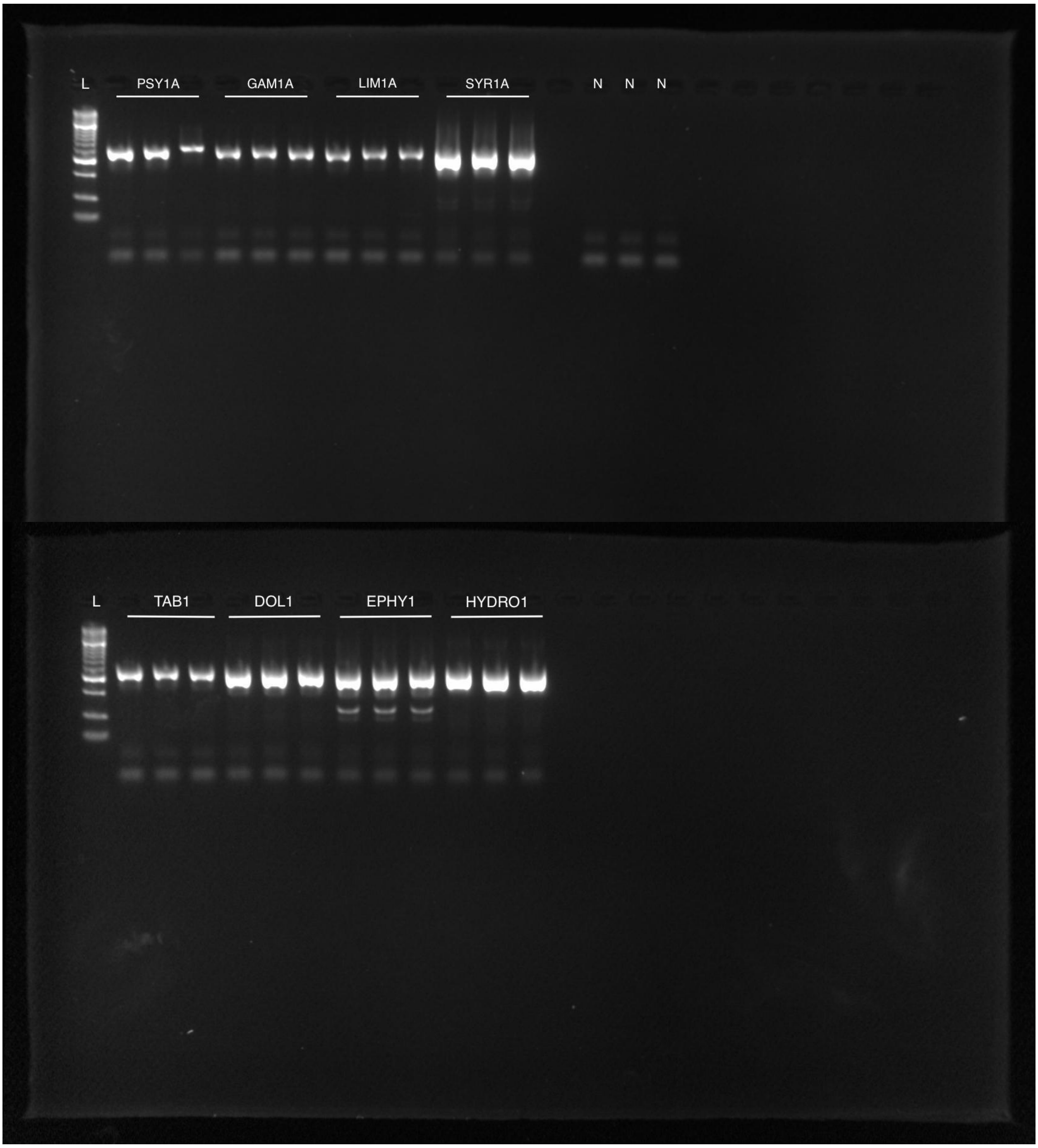


**Figure S4.** Gel images showing results of *in vitro* primer validation for primers mICOIintF and jgHCO2198. PCR products were run on 2% agarose gels with Hyperladder™ 50bp (Bioline®, London, UK) molecular weight marker (L). Tissue from the two-spotted assassin bug was used as the positive control.

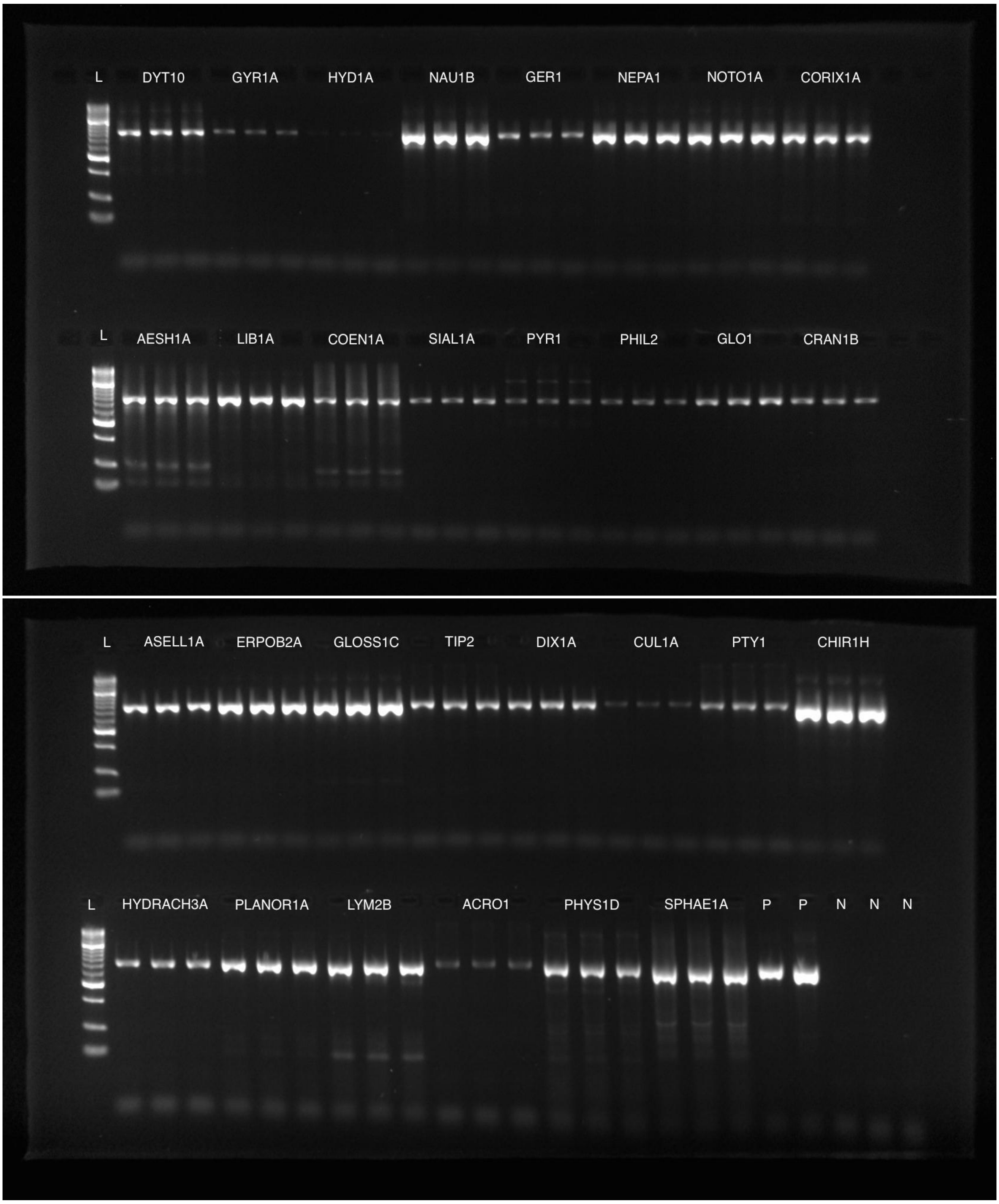


**Figure S5.** Gel images showing results of PCR for primers BF2 and BR2 (Elbrecht & Leese, 2017). PCR products were run on 2% agarose gels with Hyperladder™ 50bp (Bioline®, London, UK) molecular weight marker (L). Tissue from the two-spotted assassin bug was used as the positive control.

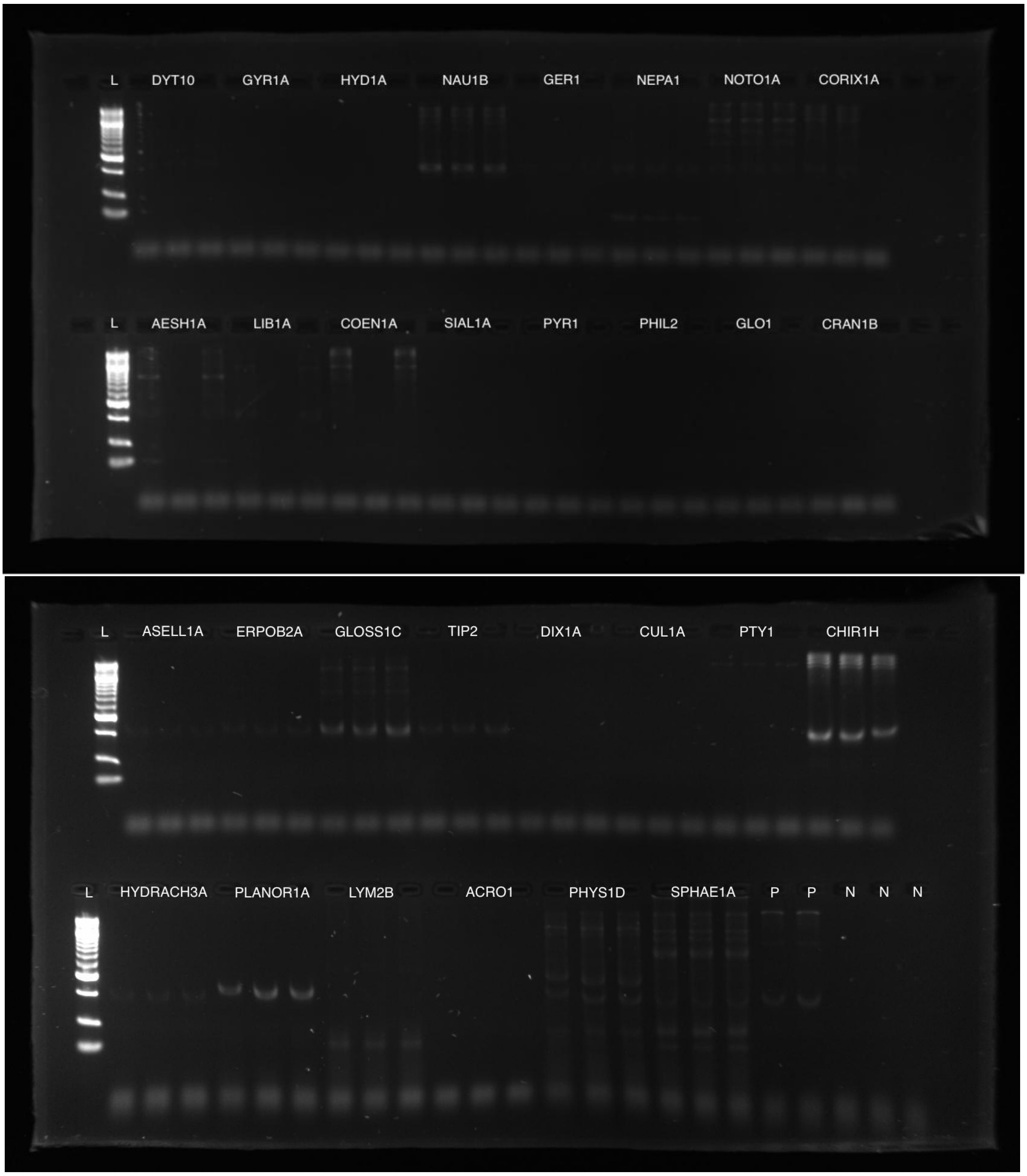


**Figure S6.** Gel images showing results of PCR for primers fwhF1 and fwhR1 (Vamos et al., 2017). PCR products were run on 2% agarose gels with Hyperladder™ 50bp (Bioline®, London, UK) molecular weight marker (L). Tissue from the two-spotted assassin bug was used as the positive control.

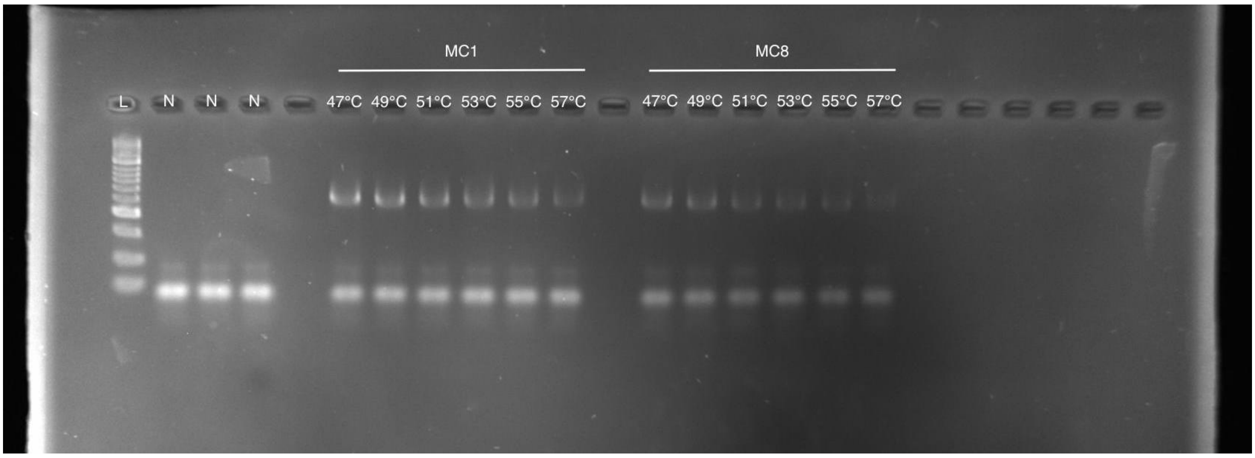


**Figure S7.** Gel image showing results of annealing temperature gradient PCR for primers mICOIintF and jgHCO2198. PCR products were run on 2% agarose gels with Hyperladder™ 50bp (Bioline®, London, UK) molecular weight marker (L).

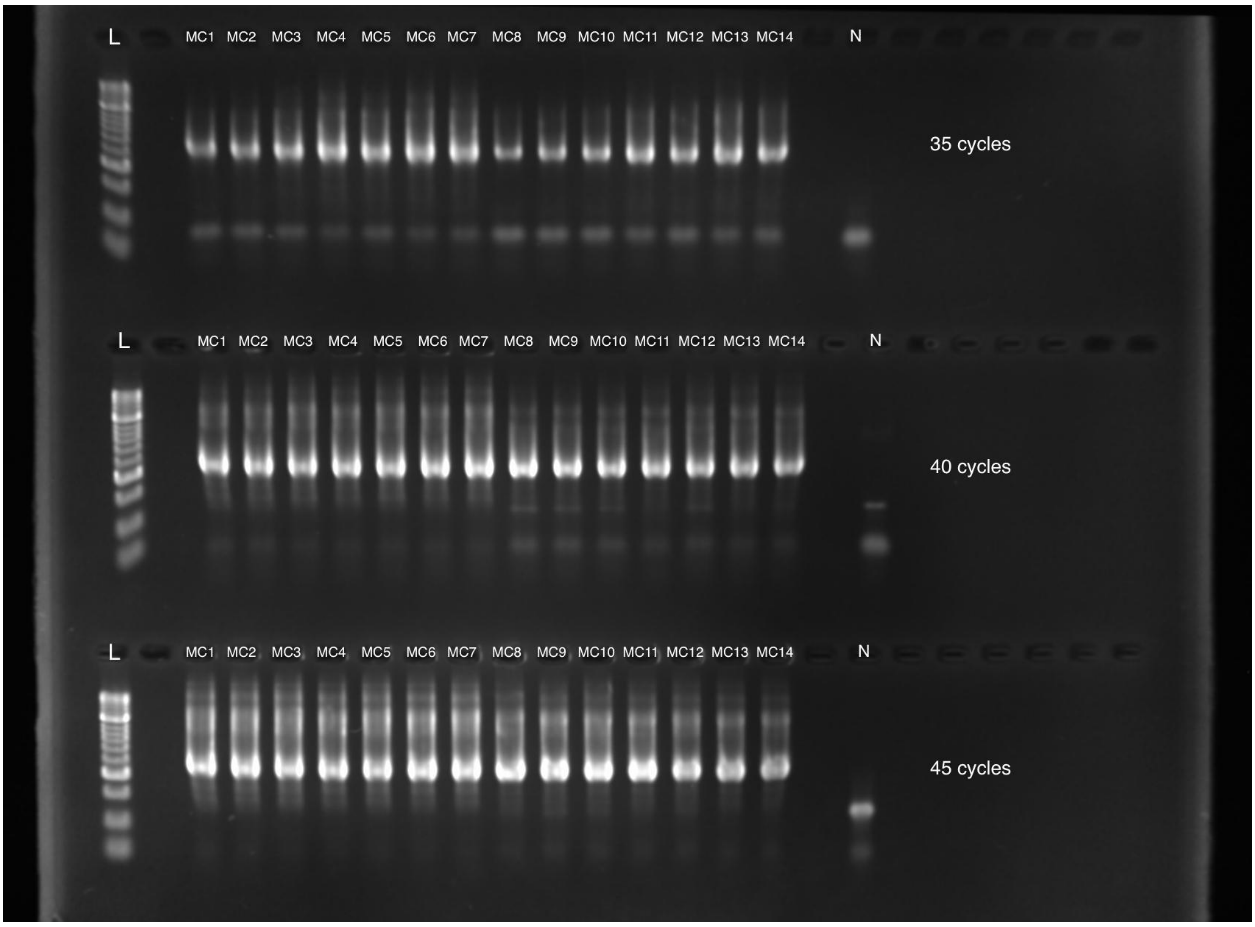


**Figure S8.** Gel images showing results of PCR cycle number optimisation for primers mICOIintF and jgHCO2198. PCR products were run on 2% agarose gels with Hyperladder™ 50bp (Bioline®, London, UK) molecular weight marker (L).

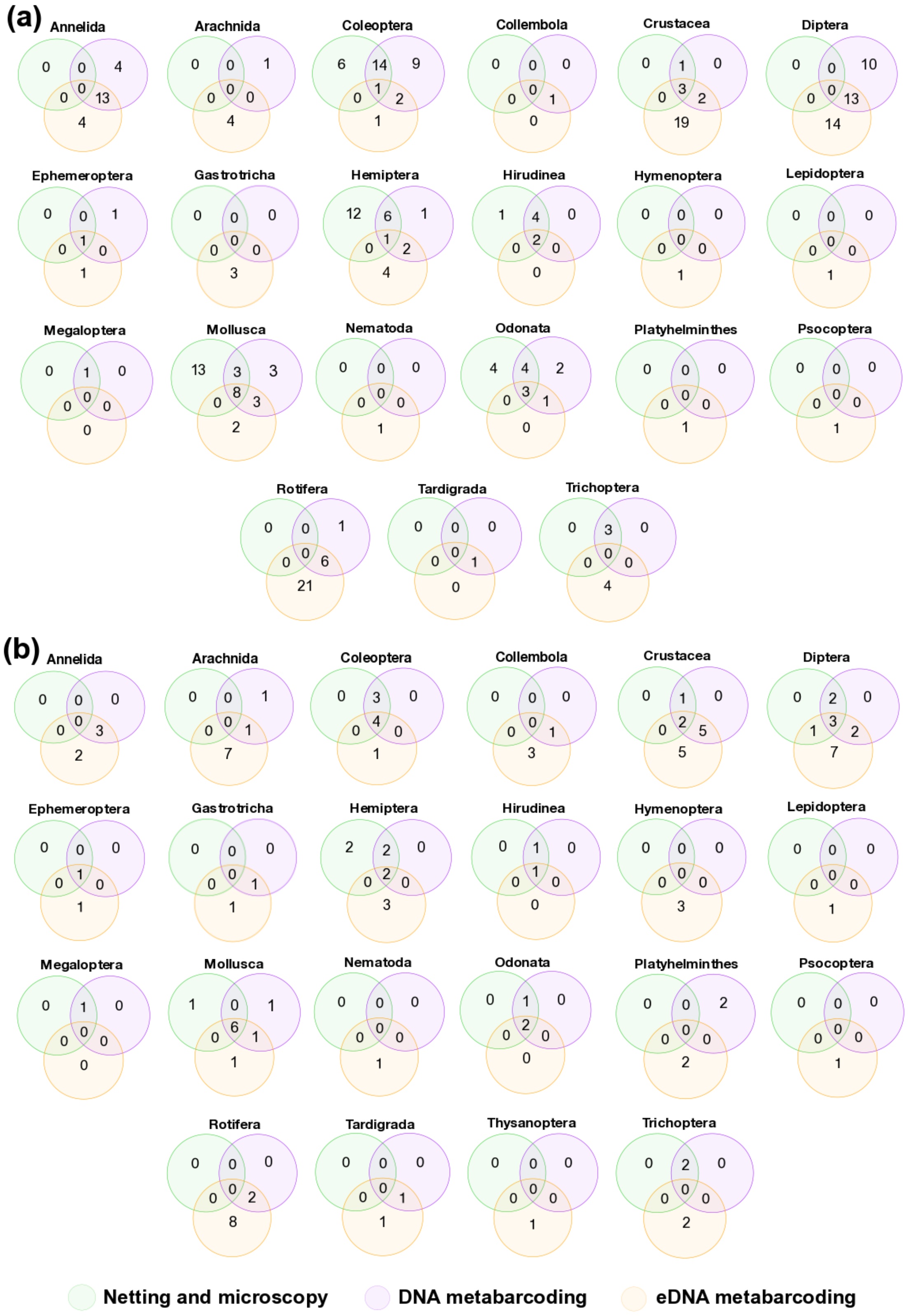


**Figure S9.** Venn diagrams which summarise the number of species **(a)** and families **(b)** detected within the major invertebrate groups across the 18 study ponds by standard methods of invertebrate assessment: sweep-netting and microscopy (green circle), DNA metabarcoding (purple circle), and eDNA metabarcoding (orange circle). Overlap in species or family detections between methods is displayed within circle intersections.

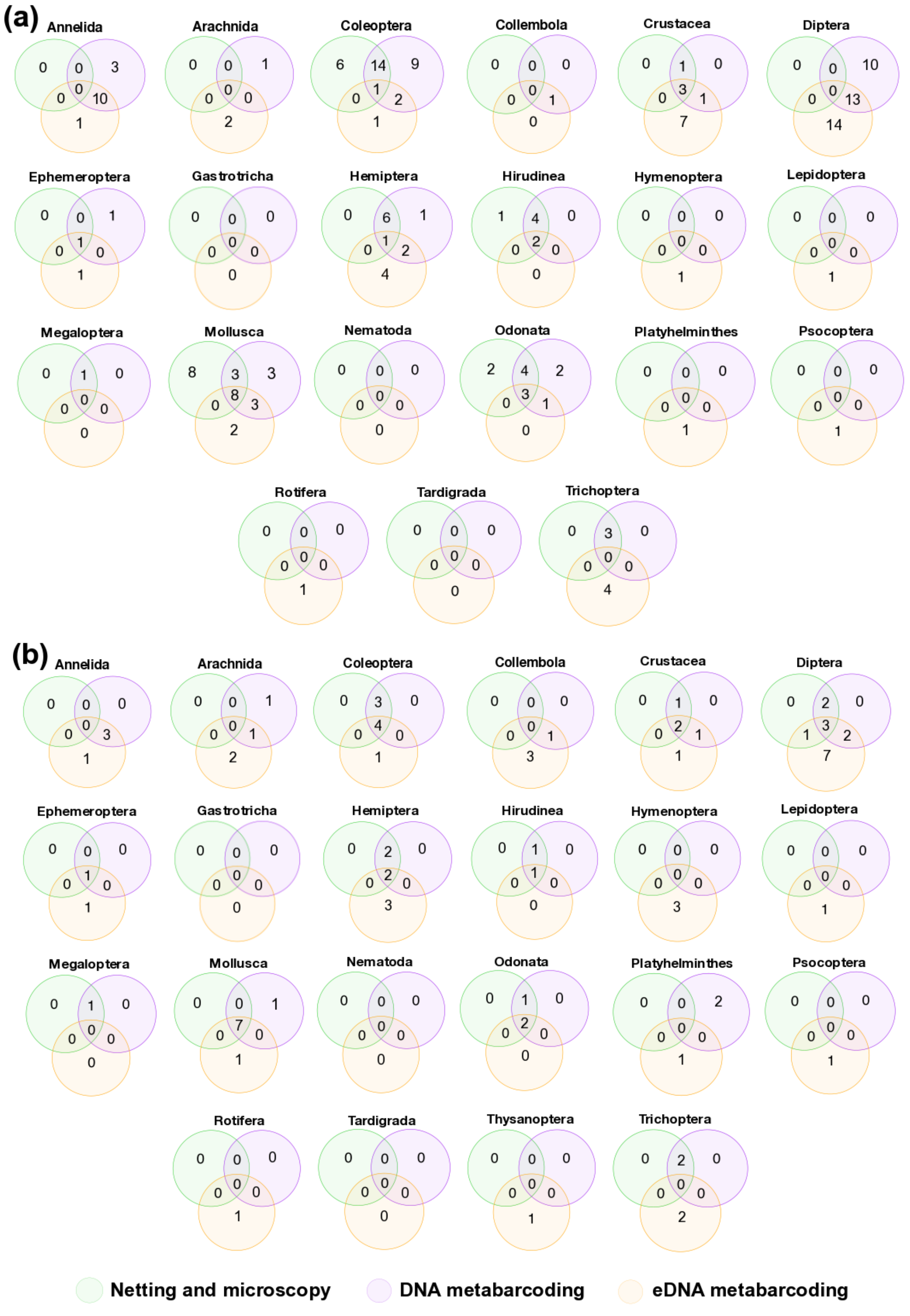


**Figure S10.** Venn diagrams which summarise the number of species **(a)** and families **(b)** detected within the major invertebrate groups across the 18 study ponds by unbiased methods of invertebrate assessment: sweep-netting and microscopy (green circle), DNA metabarcoding (purple circle), and eDNA metabarcoding (orange circle). Overlap in species or family detections between methods is displayed within circle intersections.

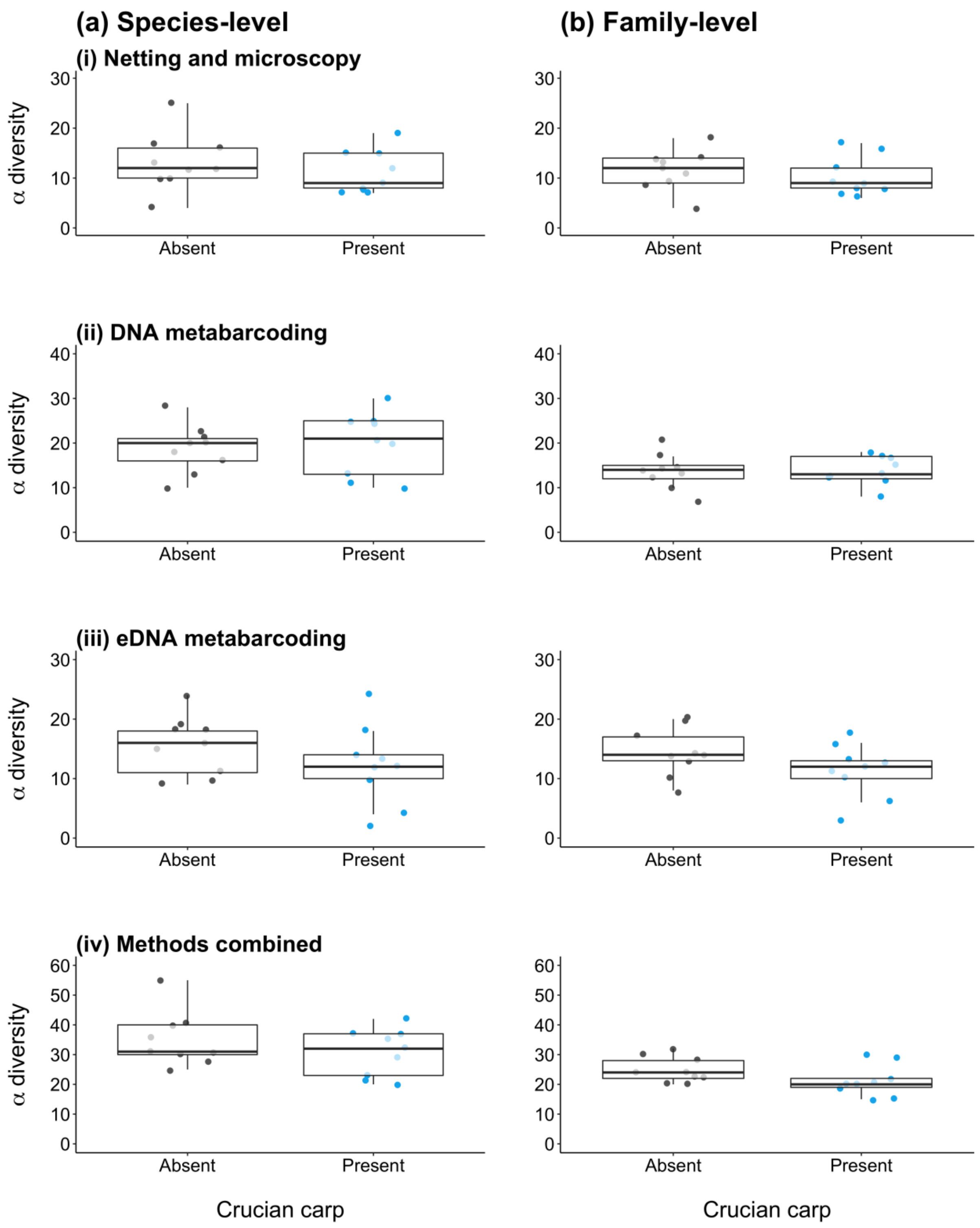


**Figure S11.** Mean alpha diversity of macroinvertebrates in ponds with crucian carp (blue points) and without fish (grey points) across Norfolk and East Yorkshire. Alpha diversity at species-level **(a)** and family-level **(b)** is shown according to unbiased methods of invertebrate assessment: sweep-netting and microscopy **(i)**, DNA metabarcoding **(ii)**, eDNA metabarcoding **(iii)**, and all methods combined **(iv)**. Boxes show 25th, 50th, and 75th percentiles, and whiskers show 5th and 95th percentiles.

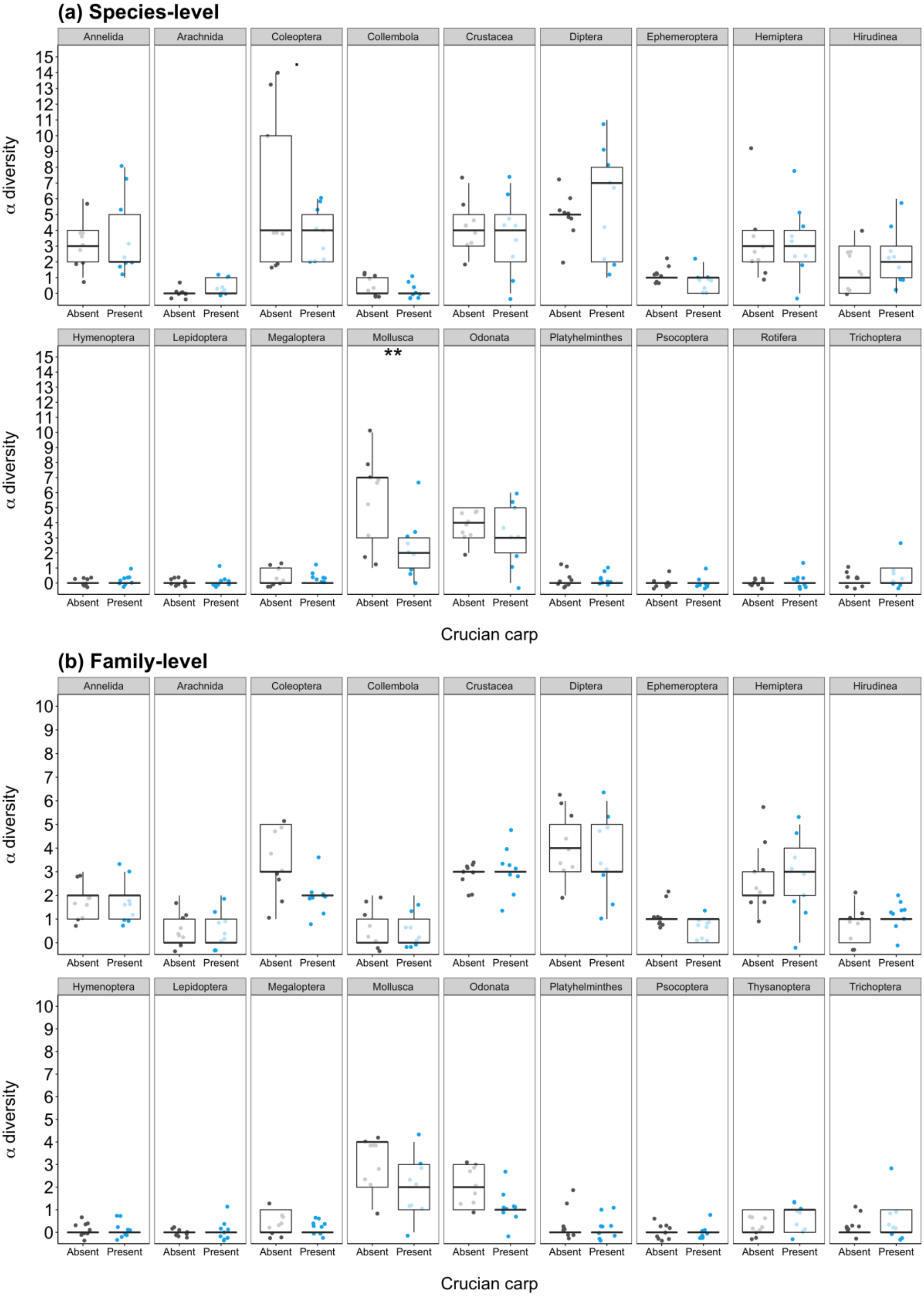


**Figure S12.** Mean alpha diversity (taxon richness) at species-level **(a)** and family-level **(b)** of the different macroinvertebrate groups identified by all three unbiased methods in ponds with crucian carp (blue points) and without fish (grey points) across Norfolk and East Yorkshire. Boxes show 25th, 50th, and 75th percentiles, and whiskers show 5th and 95th percentiles. Significant differences are indicated by asterisks (* = $P < 0.05$).

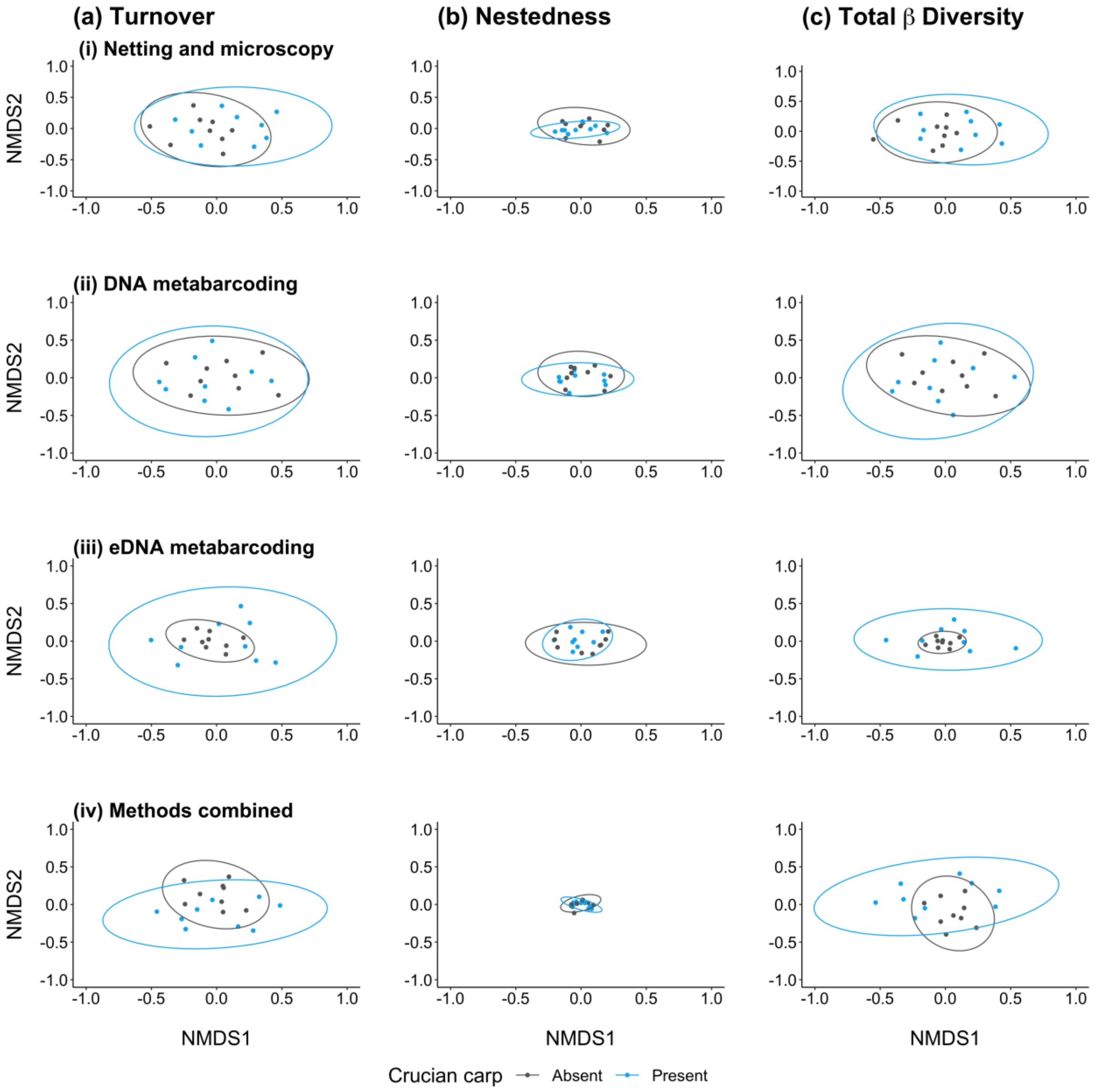


**Figure S13.** Non-metric Multidimensional Scaling (NMDS) plots of species-level macroinvertebrate communities (Jaccard dissimilarity) from ponds with crucian carp (blue points/ellipse) and without fish (grey points/ellipse) across Norfolk and East Yorkshire. The turnover **(a)** and nestedness-resultant **(b)** partitions of total beta diversity **(c)** are shown according to unbiased methods of invertebrate assessment: netting and microscopy **(i)**, DNA metabarcoding **(ii)**, eDNA metabarcoding **(iii)**, and all methods combined **(iv)**.

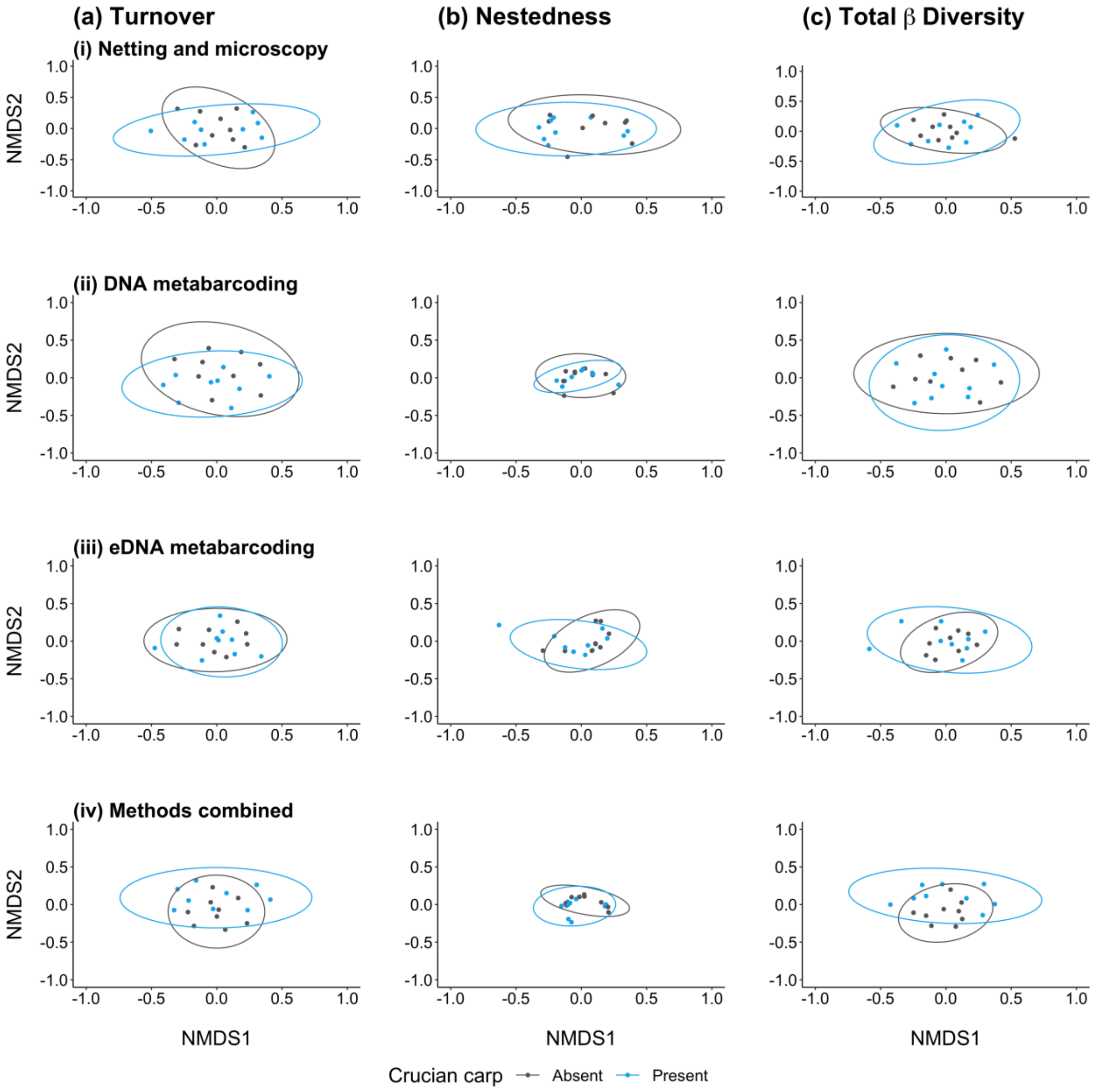


**Figure S14.** Non-metric Multidimensional Scaling (NMDS) plots of family-level macroinvertebrate communities (Jaccard dissimilarity) from ponds with crucian carp (blue points/ellipse) and without fish (grey points/ellipse) across Norfolk and East Yorkshire. The turnover **(a)** and nestedness-resultant **(b)** partitions of total beta diversity **(c)** are shown according to unbiased methods of invertebrate assessment: netting and microscopy **(i)**, DNA metabarcoding **(ii)**, eDNA metabarcoding **(iii)**, and all methods combined **(iv)**.

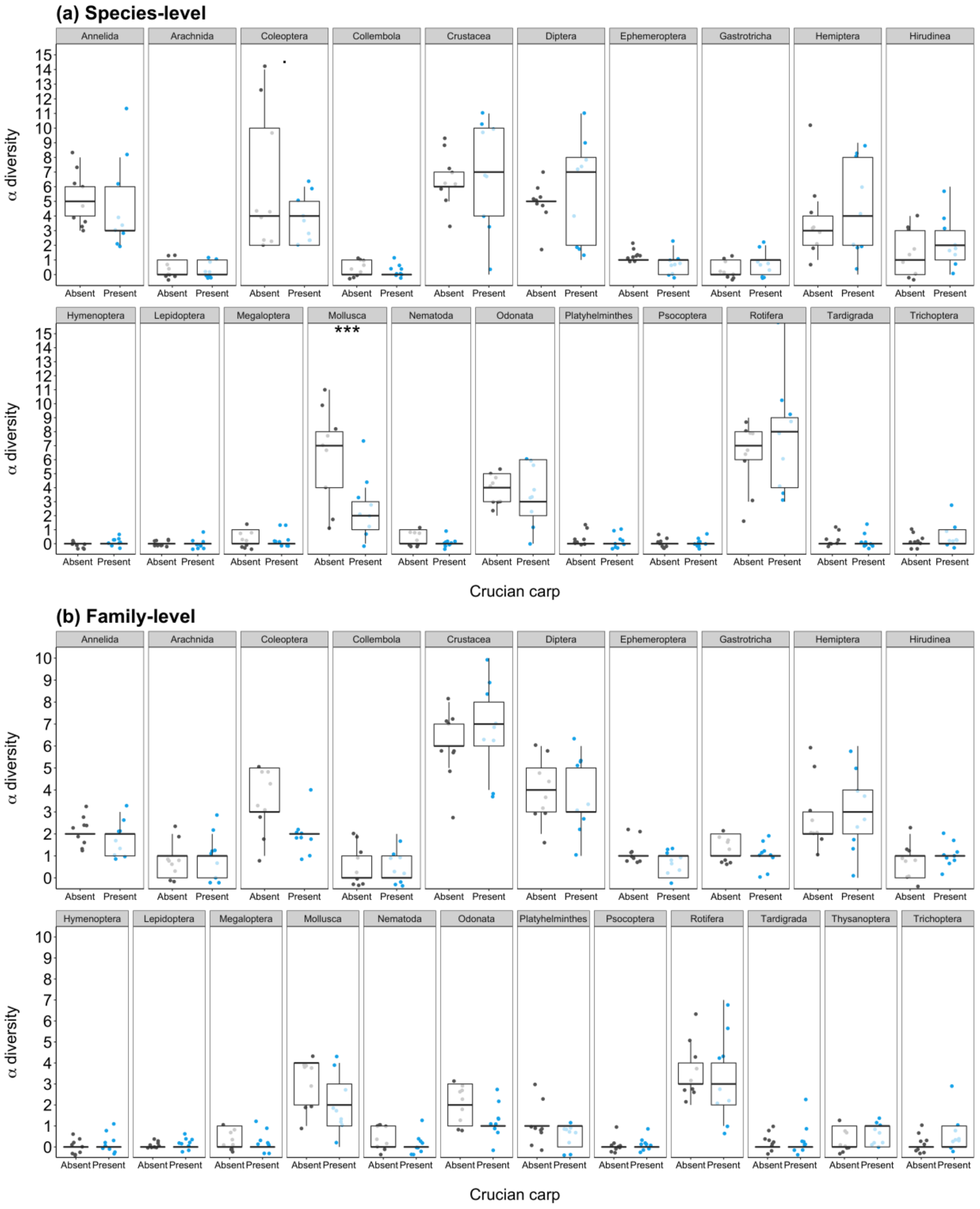


**Figure S15.** Mean alpha diversity (taxon richness) at species-level **(a)** and family-level **(b)** of the different invertebrate groups identified by all three standard methods in ponds with crucian carp (blue points) and without fish (grey points) across Norfolk and East Yorkshire. Boxes show 25th, 50th, and 75th percentiles, and whiskers show 5th and 95th percentiles. Significant differences are indicated by asterisks (* = $P < 0.05$, ** = $P < 0.01$).

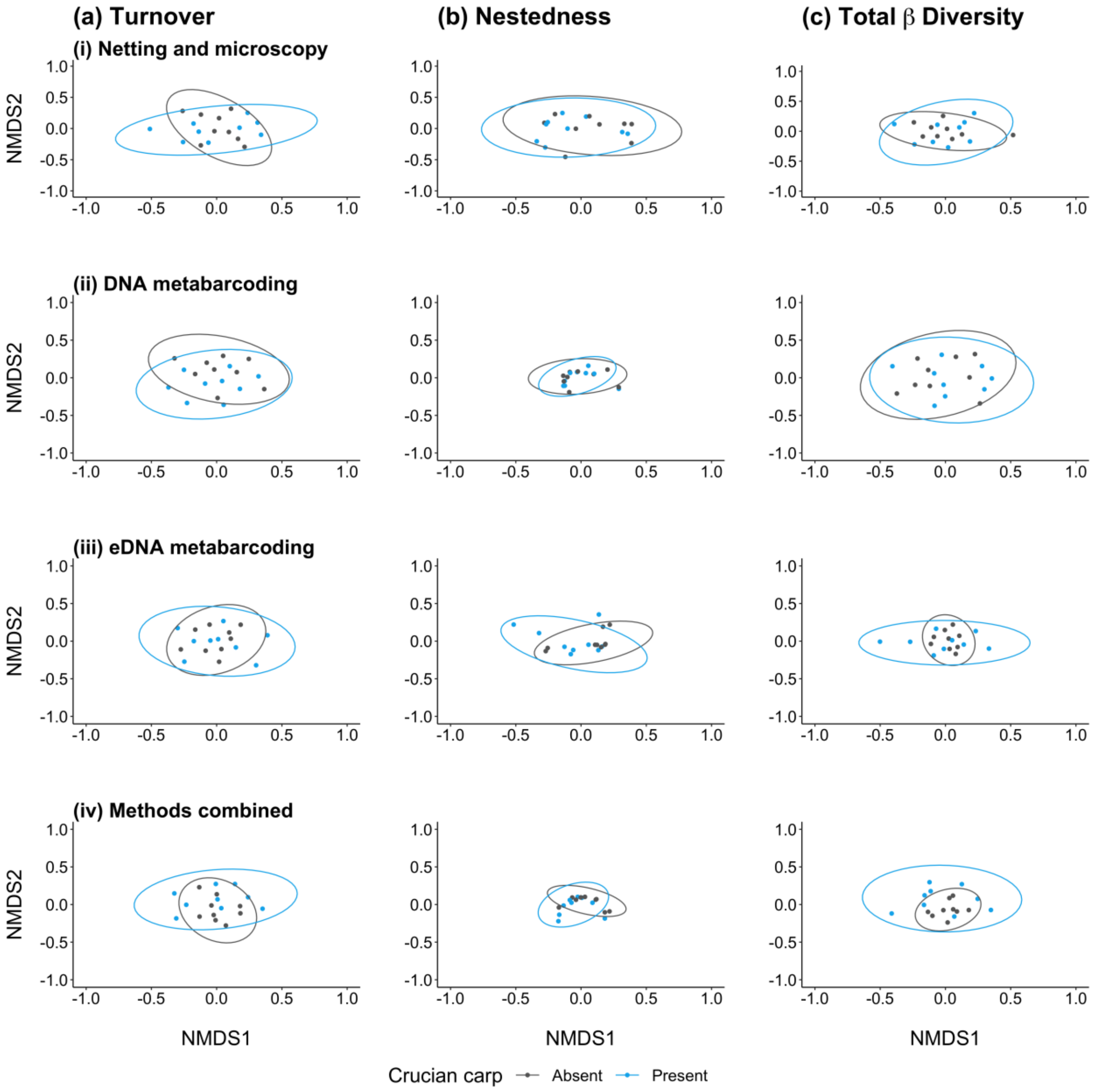


**Figure S16.** Non-metric Multidimensional Scaling (NMDS) plots of family-level invertebrate communities (Jaccard dissimilarity) from ponds with crucian carp (blue points/ellipse) and without fish (grey points/ellipse) across Norfolk and East Yorkshire. The turnover **(a)** and nestedness-resultant **(b)** partitions of total beta diversity **(c)** are shown according to standard methods of invertebrate assessment: netting and microscopy **(i)**, DNA metabarcoding **(ii)**, eDNA metabarcoding **(iii)**, and all methods combined **(iv)**.
